# Spatial co-enrichment of immature myeloid and T-cell populations predicts survival in NPM1-mutant AML

**DOI:** 10.64898/2026.11.17.767133

**Authors:** Jan C. Schroeder, Patrice Zeis, Borhan R. Saeed, Mohammed Zidane, Lanzhu Li, Ahmad Makky, Alexander Rochwarger, Christine Beschorner, Jing Zhao, Yannick A. Palmowski, Lucas Mix, Wolfgang A. Bethge, Claudia Lengerke, Christian M. Schürch

## Abstract

*NPM1*-mutant acute myeloid leukemia (AML) is the most common molecularly defined AML subtype. Outcomes vary considerably across co-mutational profiles, and the bone marrow (BM) microenvironment may shape treatment responses through immune cell interactions. Whether these interactions form spatially organized structures with independent prognostic relevance in the diagnostic BM is unknown.

We used CO-Detection by indEXing (CODEX) with a 54-marker panel and MERFISH spatial transcriptomics to profile 67 *NPM1*-mutant AML patients at diagnosis and 38 age- and sex-matched normal BM (NBM) controls on tissue microarrays from archival FFPE samples, profiling 1,494,397 cells and annotating 27 cell types.

Compared to NBM, *NPM1*-mutant AML showed striking architectural reorganization, including a loss of stroma association with adaptive immune cells and loss of NBM-specific mature myeloid and erythroid cellular neighborhoods. AML was characterized by expansion of phenotypically distinct immature myeloid (IM) populations with aberrant immunophenotypes, including CD11b+ VISTA+ and CD15+ subsets, alongside an expansion of CD8+ and PD1+ CD4+ T-cells.

While cell type and neighborhood cluster frequencies showed limited prognostic associations, spatial context-dependent interactions were more informative. Both enrichment of CD11b+ IM in adaptive immune neighborhoods, and co-enrichment with CD8+ T-cells were adversely associated with overall survival. Spatial transcriptomics identified upregulated genes associated with monocytic AML in CD11b+ IM cells and candidate immunosuppressive ligand-receptor interactions, including *TGFB1*–*TGFBR1* and *JAG1*–*NOTCH1*, with T-cells.

These findings indicate that spatial context of suppressor-immune cell interactions carries prognostic information not captured by genomics or cell type frequencies alone and is amenable to translation using routine diagnostic FFPE samples.

## INTRODUCTION

*NPM1* mutations are the most common genetic alteration in AML and define a distinct disease biology^1,2^. *NPM1*-mutant AML is more frequent in younger patients and generally associated with favorable outcomes after intensive chemotherapy^3^. However, prognosis varies with co-mutations, particularly concurrent *DNMT3A*^R882^ and *FLT3*-ITD^4^, and remains poor in refractory/relapsed disease^5^. Despite improved molecular classification, sensitive MRD monitoring^6,7^, and emerging targeted therapies including menin inhibitors^8,9^, allogeneic hematopoietic stem cell transplantation (alloHSCT) remains the only curative salvage option for refractory/relapsed patients.

Genomic and single-cell studies revealed diverse malignant differentiation states and immune/microenvironmental components in AML^10,11^. Single-cell RNA-seq has defined AML hierarchies, stem/progenitor-like states, differentiation states, immune remodeling^12,13^, and transcriptome-based *NPM1*-mutant AML subtypes potentially useful for stratification^14,15^. However, tissue dissociation destroys spatial architecture, precluding the analysis of organized leukemic and microenvironmental niches. The AML BM microenvironment shows profound immune remodeling, including immunosuppressive myeloid expansion and T-cell dysfunction linked to treatment resistance^16–19^. However, whether these populations form spatially organized niches in diagnostic BM with prognostic relevance beyond cell frequencies remains unknown. Leukemogenesis and post-treatment persistence occur within the spatially structured BM microenvironment. Normal hematopoiesis depends on local vascular, stromal, immune, megakaryocytic, erythroid, and myeloid niches that are insufficiently captured by aspiration^20,21^. AML likely remodels this architecture rather than simply replacing normal BM with blasts^22^. Recent studies, particularly of adaptive immune cells, demonstrated the importance of niche-specific spatial interactions for treatment response^23–26^, suggesting that spatial analysis provides information beyond cell type abundance or gene expression.

CO-Detection by indEXing (CODEX) combines highly granular protein phenotyping with preserved tissue architecture^20,23,27,28^, enabling single-cell identification of hematopoietic and microenvironmental compartments. CODEX and related approaches have revealed recurrent multicellular neighborhoods or ecotypes with functional and prognostic relevance^28–31^. Importantly, candidate spatial biomarkers can be translated into focused multiplex immunofluorescence assays compatible with extended routine diagnostics. *NPM1*-mutant AML represents a biologically defined but heterogeneous entity, and to date no comprehensive dataset is available for spatial biomarker discovery. Here, we performed spatial multi-omics on archival FFPE BM samples from 67 *NPM1*-mutant AML patients and identified prognostically relevant, context-dependent cell interactions not captured by genomics or cell type frequencies alone.

## METHODS

Informed consent was obtained in accordance with the Declaration of Helsinki. The study was approved by the Ethics Committee of the University of Tübingen (099/2023BO2).

The study consisted of (1) the assembly of diagnostic *NPM1*-mutant AML and age- and sex-matched normal BM (NBM) cohorts, followed by tissue microarrays (TMAs) construction; (2) the establishment and validation of a CODEX imaging and processing pipeline; (3) the development and validation of MERFISH spatial transcriptomics on FFPE BM TMAs; and (4) the design of an analysis pipeline for spatial phenotyping, cell type annotation and downstream analyses.

Detailed methodology, including experimental workflows, reagents, instrumentation, and computational analyses, is provided in the **Supplemental Information**.

## RESULTS

### High-throughput spatial profiling of *NPM1*-mutant AML BM

We constructed three TMAs from diagnostic FFPE BM biopsies of 67 treatment-naïve patients, comprising 202 cores (median, three 1-mm cores per patient; **Fig. 1A, Fig. S1**). Two NBM TMAs comprising 128 cores from 38 age- and sex-matched individuals with normal BM histology served as controls. All AML patients harbored *NPM1* mutations, and *FLT3*-ITD was detected in 26/67 patients (38.8%). First-line intensive anthracycline-based chemotherapy (7+3) was administered to 56/67 patients (83.6%), and 49/67 (73.1%) underwent alloHSCT (**Fig. 1B**; **Tables S1-S2**).

**Figure 1:**
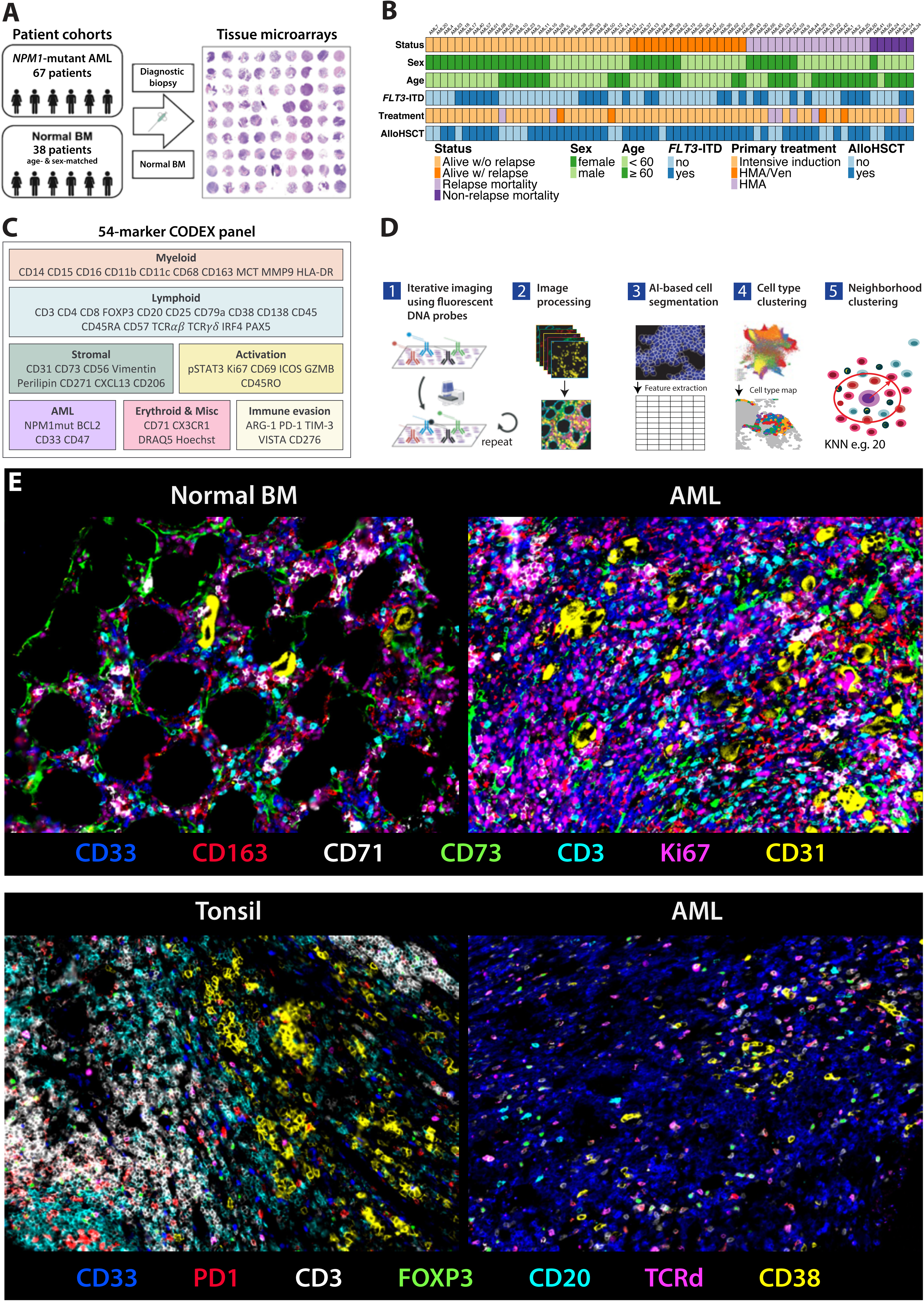
High-throughput spatial profiling of *NPM1*-mutant AML bone marrow. A: *NPM1*-mutant AML and normal bone marrow (NBM) cohorts and tissue microarray (TMA) construction for high-throughput analysis. B: Clinical annotation of *NPM1*-mutant AML cohort, including status at follow-up, patient sex and age at diagnosis, *FLT3*-ITD status, primary treatment, and receipt of allogeneic hematopoietic stem cell transplantation (alloHSCT). HMA, hypomethylating agent; Ven, venetoclax. C: 54-marker antibody panel used for CO-detection by inDEXing (CODEX) staining and iterative imaging. D: Computational image processing and analysis strategy including artificial intelligence (AI)-based segmentation, cell type annotation, and k-nearest-neighbor-based spatial window definition and Kmeans cellular neighborhood clustering. E: Seven marker-overlay images of representative *NPM1*-mutant AML, NBM, and tonsil regions depicting various hematopoietic, immune, and stromal markers.

We developed and validated a 54-marker CODEX panel covering hematopoietic, leukemic, stromal, and immune activation and evasion markers (**Fig. 1C, Table S3**). TMAs were imaged using the PhenoCycler Fusion 2.0, followed by DeepCell-based segmentation (**Fig. 1D**, **Tables S4-S5**). Representative CODEX images are shown in **Figs. 1E** and **S2**.

### BM cell type annotation

After imaging, segmentation, and artifact filtering, 1,214,491 AML and 279,906 NBM cells were recovered. Cell type (CT) annotation was achieved by unsupervised clustering with manual annotation combined with supervised classifiers, followed by harmonization and refinement using decision-tree algorithms (**Fig. 1D**; **Methods**). We identified 27 CTs in the AML dataset (**Figs. 2A**, **S3&S4**; **Table S6**). Adipocytes and megakaryocytes (MKs), which were inadequately segmented, were manually annotated using CODEX and H&E images (**Fig. 2B**), while eosinophil clusters were identified by H&E correlation. The NBM dataset comprised 35 CTs and showed the expected predominance of mature myeloid and erythroid cells (**Fig. S5**).

**Figure 2:**
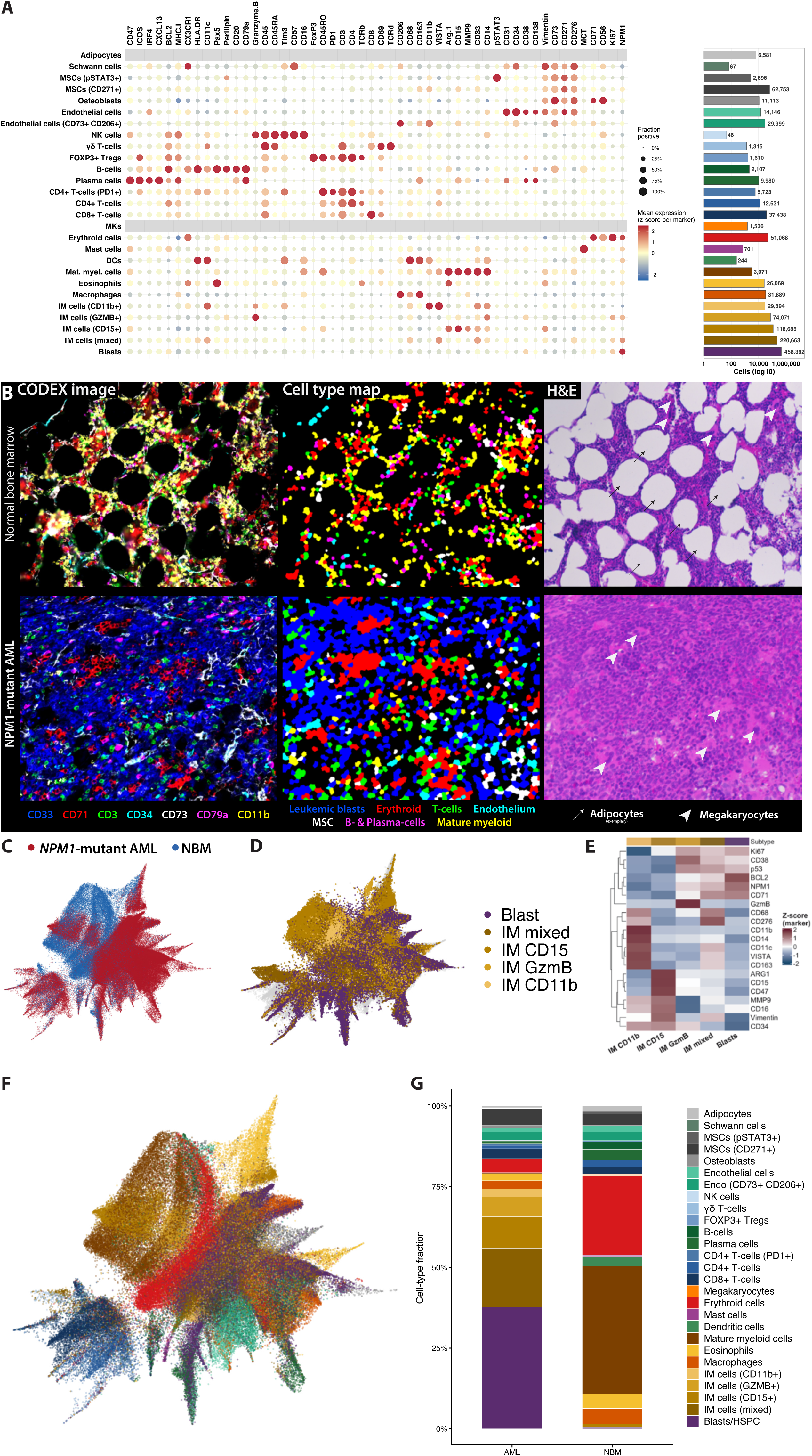
Cell type annotation and differences in BM microenvironment composition between *NPM1*-mutant AML and NBM. A: Dot plot visualizing marker expression intensity (z-score, color) and fraction of cells with detectable marker expression (dot size) for each of 27 annotated cell types (CTs) in the *NPM1*-mutant AML cohort. Bars show absolute cell number per CT on a log_10_-scale. B: Representative regions shown with CODEX staining, annotation-based spatial cell type maps, and corresponding H&E staining with indicated adipocytes and megakaryocytes. C: UMAP representation of AML versus normal bone marrow (NBM) cells in marker space. D: UMAP of NBM and AML cells in marker space highlighting leukemic blast and immature myeloid (IM) cell populations. E: Heatmap showing normalized expression of myeloid and AML markers in leukemic blast and IM cell populations. F: UMAP representation of pooled AML and NBM cells in marker space, overlaid with cell type annotation for harmonized, corresponding CTs present in both the AML and NBM annotations. G: Stacked bar chart depicting CT composition in AML and NBM cohorts.

UMAP in marker space revealed clear AML-NBM separation (**Fig. 2C**), driven by expanded leukemic blast and immature myeloid (IM) populations and independent of batch effects (**Figs. 2D**, **S6**). Leukemic blasts showed high NPM1, BCL2, and CD71 expression and lacked lineage (LIN) markers, whereas IM cells displayed monocytic (IM CD11b+), granulocytic (IM CD15+), granzyme-B-high (IM GZMB), or mixed differentiation (IM mixed). IM mixed cells most closely resembled leukemic blasts but showed partial maturation (**Fig. 2E**). Major myeloid, erythroid, lymphoid, mesenchymal and endothelial populations separated by marker expression, with some overlap between blasts and myeloid populations (**Fig. 2F**). As expected, diagnostic AML BM was enriched for blasts and IM cells, while most normal hematopoietic and lymphoid CTs were depleted (**Fig. 2G**). CTs were identified reliably across patients, including rare subtypes such as FOXP3+ regulatory T-cells (Tregs) or pSTAT3+ mesenchymal stromal cells (MSCs) (**Fig. S7**). We compared CT frequencies from CODEX with clinical BM aspiration cytology. Blast, erythroid, and myeloid frequencies correlated significantly between both methods, while lymphocytes, plasma cells, macrophages, and MKs did not, consistent with limited representation in aspirates (**Fig. S8**).

### Differences in cell type composition and proximity between AML and NBM

Analysis at the individual TMA-core level revealed near-complete loss of mature myeloid cells, dendritic cells (DCs), and B-cells in AML BM, accompanied by reduced MKs, plasma cells, and erythroid cells (**Fig. 3A**). In contrast, Treg, eosinophil, and adipocyte densities were unchanged, whereas macrophages, CD4+ T-cells, CD8+ T-cells, γδ T-cells, and stromal cells were increased. Given higher cellular density in AML BM, we additionally compared relative CT frequencies. This confirmed enrichment of CD8+ T-cells and MSCs, and unchanged γδ T-cell frequencies, whereas all other CTs were reduced (**Fig. S9**). Intercellular proximity provides a simple measure of tissue architecture, but the marked expansion of blasts and IM cells and depletion of mature myeloid and erythroid cells in AML can confound proximity analyses. We therefore compared both raw mean minimum distances and density-corrected proximity between CT pairs common to AML and NBM (**Fig. S10A**). As expected, changes in pairwise raw distance between AML and NBM were highly correlated with concordant changes in the abundance of a given CT pair (r=-0.91, **Fig. S10B**). For instance, mature myeloid cells and B-cells showed large decreases in abundance in AML and correspondingly increases in raw distance to most CTs. Conversely, CD8+ T-cells and MSCs became more abundant, associated with decreased raw distance to other CTs (**Fig. S10A**). To better distinguish spatial organization from compositional effects, we derived a density-corrected proximity metric comparing observed distances to null distributions generated by random label permutation (**Fig. 3B & S10C**; **Methods**), which exhibited moderate correlation with CT abundance (r=0.61). This revealed a module consisting of CD8+ T-cells, macrophages, MSCs, osteoblasts and endothelium with lower-than-expected mutual attraction in AML and conversely higher-than-expected self-association, especially of CD8+ T-cells. Erythroid and mature myeloid cells showed markedly reduced segregation in AML relative to NBM, whereas erythroid cells showed lower-than-expected proximity to the CD8/macrophages/stroma module. Erythroid cells and macrophages were the only pair showing a large, concordant decrease in both raw and density-corrected proximity (**Fig. S10D**), as previously reported in myelodysplastic syndrome^32,33^.

**Figure 3:**
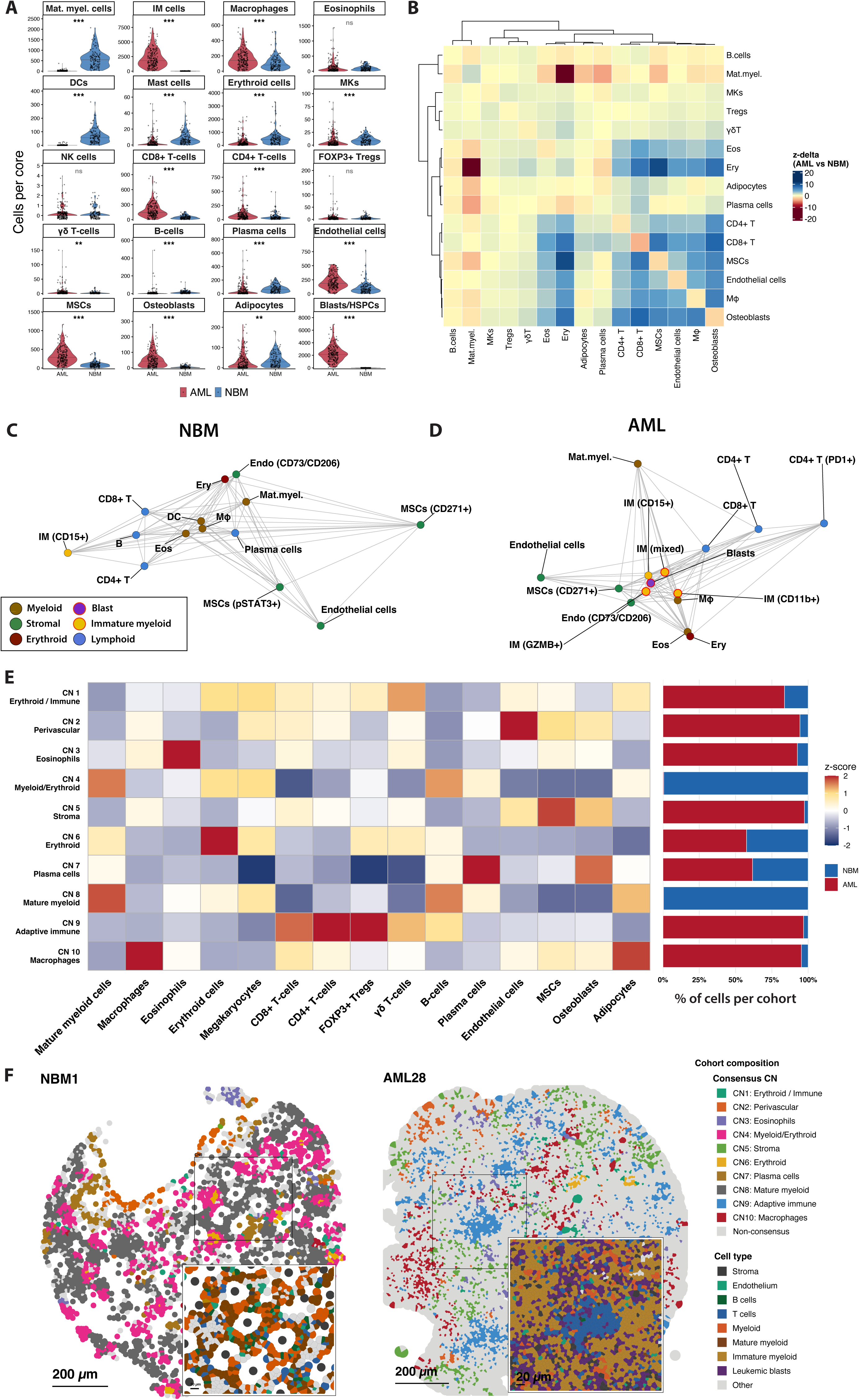
Cellular neighborhood analysis identifies shared and unique architectural features of NBM and *NPM1*-mutant AML BM microenvironment. A: Per-subject cell type (CT) numbers for CTs present in both AML and normal bone marrow (NBM) datasets per tissue microarray (TMA) core visualized as violin plots. Blasts in the AML dataset were matched to hematopoietic stem and progenitor cells (HSPCs) in the NBM dataset, respectively. Significance of frequency differences was assessed by two-sided Wilcoxon rank-sum test with Benjamini–Hochberg FDR correction. Annotation symbols: *** FDR < 0.001, ** < 0.01, * < 0.05, ns = not significant. B: Raw mean-minimum distance between two CTs was quantified per TMA core as the mean nearest-neighbor distance, then averaged across cores. To derive a density-corrected proximity metric, the observed distance was compared to a null distribution built by shuffling CT identity 500 times between the two CTs only, yielding a null mean and standard deviation per region. Density-corrected proximity was quantified as a z-score, z = (observed − null mean) / null SD. To compare shifts in pairwise density-corrected proximity between NBM and AML, the difference between z-scores (z-delta = z_AML − z_NBM) was computed and visualized in a heatmap for a set of 15 consensus CTs (105 cross-type pairs plus 15 self-associations on the diagonal). Blasts, IM cells, and three CTs with frequency below 0.1% in the AML dataset and insufficient representation across AML cores (dendritic cells, NK cells, and mast cells) were excluded. Negative z-delta indicates the pair is more attracted relative to chance in AML than in NBM. C & D: Multidimensional-scaling proximity map of NBM (C) and AML (D) CTs, embedded from pairwise raw mean-minimum nearest-neighbor distance (non-metric MDS, isoMDS, k=2). Grey edges connect each CT to up to its 4 nearest spatial neighbors (or any additional pair within 25% of that distance). A subset of low-abundance CTs is omitted from this view for visual clarity (full network in **Fig. S11**). Panels share a common physical scale, so relative spread/distance is comparable between NBM and AML. Absolute position and orientation of CT nodes are not comparable, as each cohort was embedded independently. E: Consensus Cellular neighborhood (CN) clustering analysis was performed to identify shared and specific architectural features. This analysis used only consensus CTs present in both NBM and AML datasets, excluding immature myeloid (IM) cells, blasts, and three CTs with frequency below 0.1% in the AML dataset and insufficient representation across AML cores (dendritic cells, NK cells, and mast cells). The heatmap shows relative CT composition of each consensus CN cluster, each frequency double-z normalized within CT and CN. Bar charts indicate the frequency of cells per consensus CN belonging to either the NBM or the AML cohort. F: Exemplary Voronoi maps for shared consensus CNs, with insets displaying consensus CTs labeled by broad lineage group plus leukemic blasts and immature myeloid cells in AML. Lower cell density in the NBM regions causes some cells to be plotted disproportionately large. To limit this, Voronoi maps were plotted from centroids with maximum radius set to 20 µm, resulting in adipocytes being plotted smaller than their true size. Mφ, macrophages.

To visualize changes induced by leukemic transformation, we performed non-metric multidimensional scaling on pairwise raw distances across consensus and tissue-specific CTs, including blasts and IM cells (**Fig. 3C-D**, full network, **Fig. S11**). In NBM, this revealed an adaptive immune module of CD4+/CD8+ T-cells, B-cells, DCs, macrophages and eosinophils, that also associated with endothelial cells and MSCs. Together with the density-corrected proximity analyses (**Fig. 3B**), this suggested that in AML, blasts and IM cells interposed between the adaptive immune module and MSCs and endothelium. Blasts and IM cells clustered tightly together and were consistently closer to macrophages, CD73+ CD206+ endothelium, and CD271+ MSCs than to plasma cells, osteoblasts, pSTAT3+ MSCs, or MKs (**Figs. 3D & S11**). Leukemic transformation was also associated with increased proximity of PD1+CD4+ T-cells to other immune populations. Together, these findings suggest that blast/IM expansion correlated with disrupted adaptive immune–stromal interaction in AML. Given the large number of CTs and demonstrated abundance-driven proximity changes, this network should be interpreted as a descriptive representation of global tissue architecture.

### Cellular neighborhood analysis reveals structural reorganization of the BM microenvironment in *NPM1*-mutant AML

To obtain an unbiased, data-driven view of tissue architecture, we performed cellular neighborhood (CN) clustering^28^. We first defined consensus CNs across NBM and AML using shared CTs, enabling direct comparison of spatial organization (**Fig. 3E**). This revealed NBM-specific, AML-dominated and shared CNs. Two NBM-exclusive CNs showed maturing myelopoiesis, with or without erythropoiesis, spatially associated with B-cells, MKs, and adipocytes (CN4, CN8).

Consensus CNs appeared less organized and sparser in AML than NBM, consistent with the predominance of blasts and IM cells, which were excluded from consensus CT-based neighborhood inference (**Fig. 3F**). An erythroid neighborhood enriched with MKs, Tregs and γδ T-cells (consensus CN6) and a plasma cell clusters neighborhood associated with osteoblasts (consensus CN7), were similarly represented in NBM and AML. However, distance analyses revealed altered spatial relationships within these CNs: in AML, erythroid cells showed increased proximity to mature myeloid cells but decreased proximity to macrophages (**Fig. 3B, S10D**). The remaining consensus CNs were predominantly AML-enriched, comprising adaptive immune cell clusters (consensus CN9); immune/erythroid-enriched and immune/macrophage-enriched neighborhoods (consensus CN1 and CN10), eosinophil-enriched regions (consensus CN3), and perivascular or periosteal stromal niches (consensus CN2 and CN5) (**Fig. 3E**).

To capture spatial heterogeneity within the *NPM1*-mutant AML BM microenvironment, we next performed CN clustering across all cells in the AML dataset only. Ten CNs were identified, exhibiting distinct enrichments of CTs (**Figs. 4A, S12**). As expected, most CNs were dominated by blasts and IM, whereas CN6 and CN10 showed pronounced erythroid and stromal enrichment, respectively (**Fig. 4B**). To determine whether CN identity was driven by CT composition, we defined CN units as all cells assigned to a given CN within an individual TMA core and clustered these units by CT composition (**Fig. 4C**). CN6 and CN10 units formed distinct erythroid and stromal clusters, whereas the remaining CNs formed a continuum. Thus, beyond the erythroid and stromal niches, CN identity appears to reflect recurrent spatial organization rather than CT composition alone. CN units were broadly distributed across TMA cores, indicating that CN assignment was not driven by individual patients or acquisition regions (**Fig. S13**).

**Figure 4:**
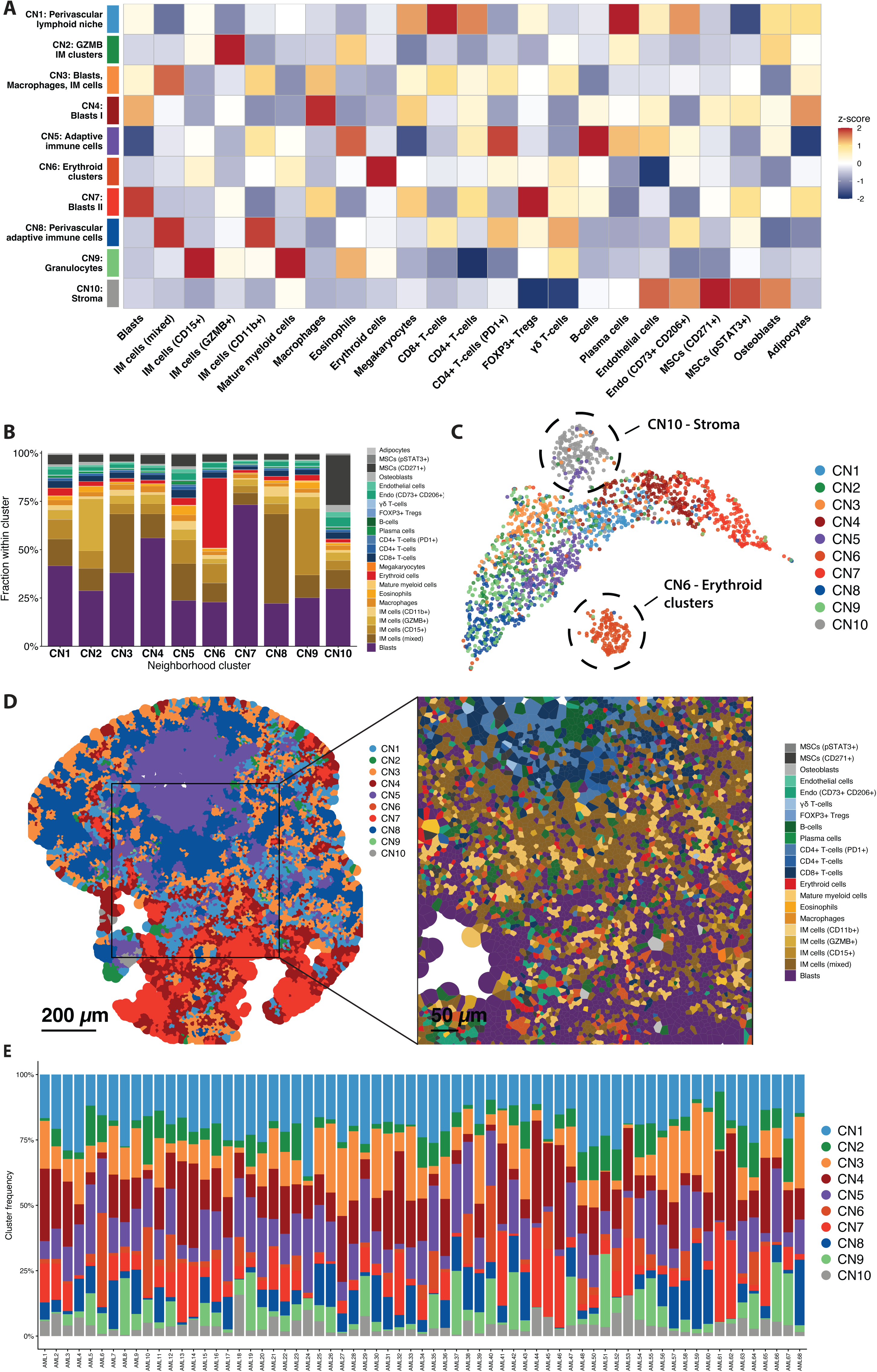
NPM1-mutant AML exhibits recurrent cellular neighborhoods. A: AML cellular neighborhood (CN) clustering performed on all cell types (CTs) in the AML dataset. The heatmap shows relative CT composition of each CN cluster, each frequency double-z normalized within CT and CN. B: Stacked bar charts depicting CT composition of each AML CN. C: UMAP showing distribution of CN units in CT composition space. CN units were defined as the entirety of cells belonging to one CN cluster in one TMA core. Each dot represents a CN unit projected on CT composition space by UMAP. CN10 (stroma) and CN6 (erythroid clusters) cluster separately and are highlighted by dashed lassos. D: Exemplary Voronoi maps for AML CNs (left), with an inset displaying corresponding CT positions (right). E: CN composition per patient in the *NPM1*-mutant AML cohort.

Beyond erythroid and stromal CNs, several neighborhoods captured patterns of immune cell colocalization. CN5 was enriched in adaptive immune cells, including FOXP3+ Tregs, B-cells and plasma cells. CN1 and CN8 represented perivascular niches: CN1 with perivascular lymphoid cells, MKs and stroma; CN8 enriched in CD73+CD206+ endothelium, PD1+CD4+ T-cells, and CD11b+ and mixed IM cells. CN9 represented a granulocytic niche enriched in CD15+ IM and mature myeloid cells, eosinophils, and some γδ T-cells. The remaining CNs were dominated by blasts and IM CTs. CN4 and CN7 were enriched in blasts, MKs and macrophages, while CN3 showed additional enrichment in mixed IM cells. CN2 was dominated by GZMB+ IM cells and enriched in osteoblasts and eosinophils. Representative regions enriched in blasts and distinct IM CTs are shown in **Fig. 4D**. All CNs were consistently detected across the cohort, supporting recurrent architectural elements in *NPM1*-mutant AML BM (**Fig. 4E**). These AML dataset-derived CNs were used for further analyses and prognostic associations.

### BM architecture is associated with clinical outcomes in *NPM1*-mutant AML

We next assessed associations between CN abundance, cellular composition, and clinical outcomes. As expected, blast-dominated CN4 and CN7 correlated positively with diagnostic BM blast counts, while CN5 (adaptive immune cells) and CN9 (granulocytes) showed inverse correlations (**Fig. 5A**). White blood cell counts correlated positively with CN4 (blasts I), CN3 (blasts, macrophages, IM cells) and, interestingly, CN8 (perivascular adaptive immune cells), and inversely with CN6 (erythroid clusters). Elevated C-reactive protein at diagnosis was associated with CN1 (perivascular lymphoid niche) and CN4. To establish a clinical and cellular reference for survival analyses, we first assessed associations of clinical variables and CT frequencies with overall survival (OS) using univariate Cox regression. As expected, age was strongly associated with inferior OS (HR 3.69, p<0.001), whereas other clinical variables in our cohort were not (**Fig. S14A**). Among CTs, higher frequencies of CD15+ and GZMB+ IM cells were associated with improved OS (HRs 0.63 and 0.65, respectively; p<0.05; **Fig. S14B**). We next performed Cox regression using normalized CN frequencies. Higher abundance of CN2 (GZMB+ IM cells) was significantly associated with improved OS, consistent with the corresponding CT-level association. In addition, enrichment of CN5 (adaptive immune cells) was associated with improved relapse-free survival (**Fig. 5B**), suggesting a potential role of adaptive immune cell interactions in limiting relapse. Overall, however, CT and CN frequencies alone showed limited prognostic associations with clinical outcomes.

**Figure 5:**
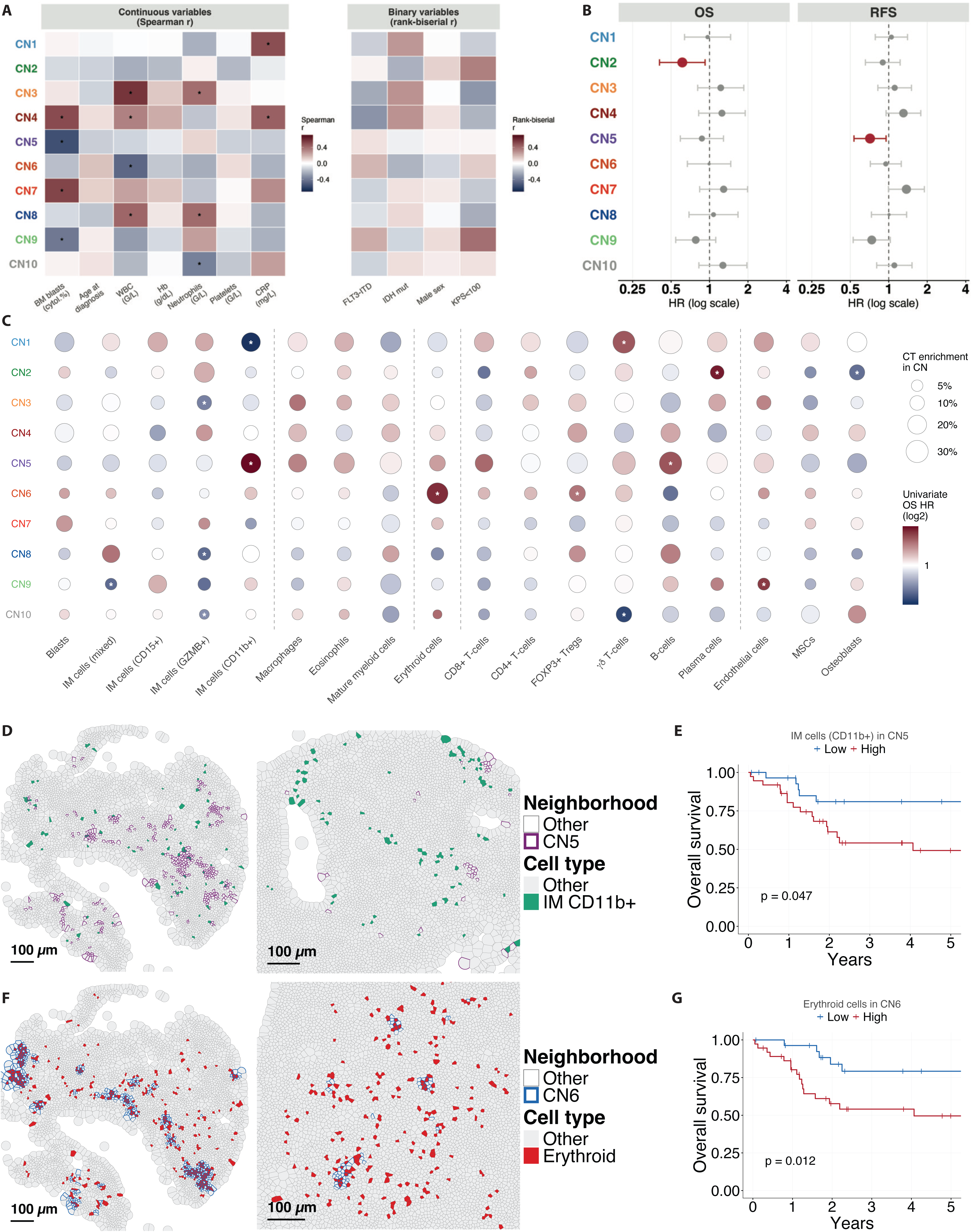
Association of architectural features with clinical baseline variables and outcomes. A: Correlation of normalized cellular neighborhood (CN) cluster enrichment with continuous and binary baseline variables. Continuous variables were tested by two-sided Spearman rank correlation; binary variables were tested by two-sided Wilcoxon rank-sum test. P-values were adjusted separately within the continuous and binary variable sets using the Benjamini–Hochberg method; asterisk denotes FDR < 0.05. B: Univariate Cox regression for association of CN frequency per patient with overall survival (OS) and relapse-free survival (RFS). C: Dot plot visualizing cell type (CT) enrichment in CNs (dot size) and corresponding univariate Cox regression hazard ratios for OS per 1 SD of normalized enrichment (color). Asterisk, p<0.05. D: CD11b+ immature myeloid (IM) cell enrichment in CN5 (adaptive immune cells), exemplary Voronoi maps illustrating low and high enrichment. E: CD11b+ IM cell enrichment in CN5, Kaplan-Meier estimated OS and log-rank p-value for low and high enrichment. F: Erythroid cell enrichment in CN6 (erythroid clusters), exemplary Voronoi maps illustrating low and high enrichment. G: Erythroid cell enrichment in CN6, Kaplan-Meier estimated OS and log-rank p-value for low and high enrichment.

Because intercellular interactions are context-dependent, we analyzed differential CT enrichment within individual CNs, which revealed several prognostically relevant spatial associations. CD11b+ IM cell enrichment in CN5 (adaptive immune cells) was associated with inferior survival, while enrichment in CN1 (perivascular lymphoid niche) predicted improved survival (**Fig. 5C-E**). In contrast, GZMB+ IM cell enrichment was associated with improved OS across multiple CNs, consistent with its favorable CT-level association (**Fig. 5C**). Erythroid enrichment in CN6 (erythroid cluster) was associated with inferior outcome (**Fig. 5F-G**), potentially reflecting disruption of the erythropoietic niche and altered myeloid/stromal interactions in AML versus NBM (**Fig. S10**). Importantly, several prognostic associations involved CT enrichments within CNs primarily defined by other cell populations, highlighting the context-dependence of these effects. For example, γδ T-cells were enriched in CN1, where their abundance was associated with inferior outcome, but showed a favorable association when enriched in CN10, despite being less frequent there. Similarly, plasma cells were relatively uncommon in CN2, yet their enrichment there was associated with inferior survival (**Fig. 5C**). These findings indicate that the prognostic relevance of individual CTs depends not only on their abundance, but also on their spatial context.

### Proximity and neighborhood co-enrichment analyses identify modules of IM and adaptive immune cells

Expanding on the CN analysis, we assessed pairwise CT interactions using raw and density-corrected distances (**Fig. 6A**). Raw distances showed a central module including IM cells, blasts, CD8+ T-cells, macrophages and CD271+ MSCs that showed high mutual proximity. Osteoblasts and plasma cells self-clustered but remained relatively distant from other CTs. Density-corrected proximity analysis suggested higher-than expected self-association across CTs and, interestingly, overall less-than-expected proximity between most CTs in the AML dataset (**Fig. S15A**). However, both raw and density-corrected proximity metrics strongly correlated with CT abundance (r=-0.91 and +0.74; **Figs. S15B-C**), suggesting that both proximity metrics are significantly confounded in this dataset. Interestingly, CD4+ and CD8+ T-cells showed concordant high raw and density-corrected proximity, while osteoblasts exhibited concordant low raw and corrected proximity to multiple other CTs (**Fig. S15D**), indicating that these observations are not primarily abundance-driven. Exploratory survival analysis identified several adverse associations, including the proximity of pSTAT3+ MSCs to Tregs and CD4+ PD1+ T-cells, and of Tregs and MKs, although these require cautious interpretation given the influence of CT abundance (**Fig. S16**). Proximity between GZMB+ and CD15+ IM cells was associated with increased survival, in line with the same-direction survival associations for the frequency of these CTs, and the frequency of CN2 (GZMB IM clusters).

**Figure 6:**
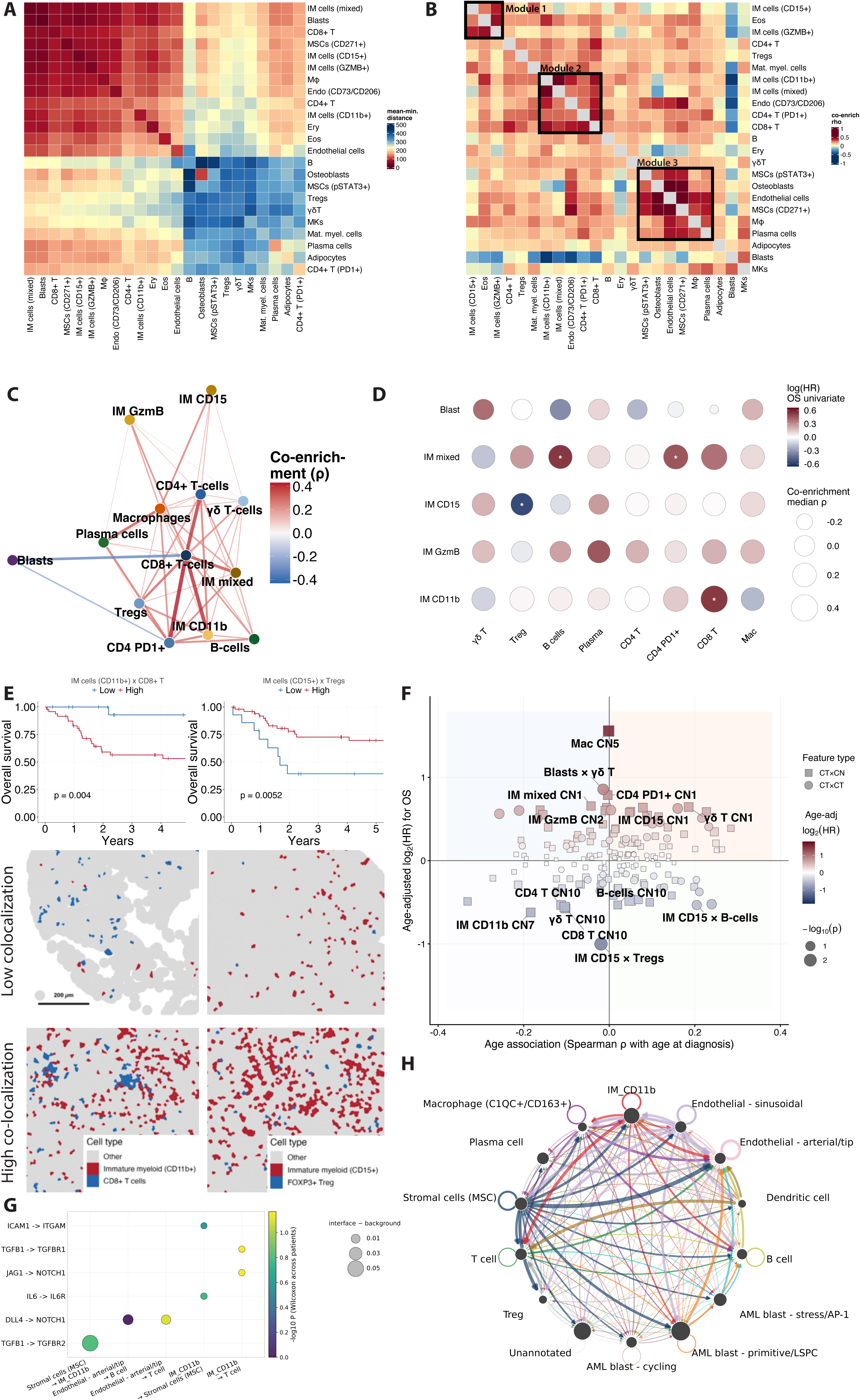
Spatial interaction of suppressive myeloid and exhausted immune cell populations correlates with treatment outcomes. A: Heatmap showing raw mean-minimum distance (µm) between cell type (CT) pairs, averaged across regions. B: Heatmap showing CT co-enrichment within the same CNs per patient. Black boxes indicate modules of highly co-enriched CTs. C: Force-directed network analysis based on CT x CT co-enrichment results, restricted to associations between adaptive immune cells, immature myeloid (IM) cells and leukemic blasts. D: Dot plot visualizing CT x CT co-enrichment (dot size) and univariate hazard ratios (HRs) for overall survival (OS) per 1 SD of normalized co-enrichment (color). Asterisk, p<0.05. E: Kaplan-Meier estimated OS with log-rank p-value (top) and exemplary Voronoi maps for low (center) vs. high (bottom) co-enrichment of CD8+ T-cells with CD11b+ IM cells (left), and of FOXP3+ Tregs with CD15+ IM cells (right). F: Scatterplot showing age-adjusted HRs for OS and age-correlation for CT x CN enrichment and CT x CT co-enrichment to identify age-independent prognostic features. G: Ligands and receptors co-expressed on spatially associated CTs, based on neighborhood radius (90^th^ percentile of the distance to each object’s 16^th^ nearest neighbor, median 31.2 µm). For each ligand-receptor pair, the per patient values were tested against zero with the Wilcoxon signed rank test and p-values were corrected for multiple testing across all pairs with the Benjamini-Hochberg procedure. H: Summed ligand-receptor interaction strength for a curated list of 49 ligand-receptor pairs (45 ligands and 40 receptors) between CTs identified by MERFISH spatial transcriptomics. Edges weighted by mean ligand x mean receptor expression, filtered by Benjamini-Hochberg correction (q<0.05). Mφ, macrophages.

Building on the recurrent CN architecture across *NPM1*-mutant AML, we examined the co-enrichment of CT pairs within CNs. Interestingly, CN co-enrichment was weakly correlated to CT abundance (r=-0.27) and unrelated to pairwise raw distance (r=0.11), consistent with these metrics capturing distinct spatial features: physical adjacency, which is strongly influenced by CT abundance, versus compositional co-variation across CNs. Neighborhood co-enrichment analysis identified three modules (**Fig. 6B**): CD15+ IM, GZMB+ IM, and eosinophils (module 1); CD11b+ IM, mixed IM, CD73+ endothelial cells, CD4+ T-cells and CD8+ T-cells (module 2), with additional associations to Tregs and mature myeloid cells; and stromal cells, plasma cells and macrophages (module 3). Notably, despite their high raw proximity to many CTs, blasts showed little CN co-enrichment except with MKs, consistent with their clustering in blast-dominated neighborhoods.

### Prognostically relevant spatial interactions between IM cells and T-cells

The co-enrichment of adaptive immune and IM cells (module 2), and the adverse prognostic association of CD11b+ IM enrichment in the adaptive immune CN5 (**Fig. 5D-E**), prompted us to further investigate IM-adaptive immune cell interactions. Expansion and exhaustion of CD8+ T-cell subsets have previously been described in *NPM1*-mutant AML^19,23^. Accordingly, we observed increased densities of CD8+ T-cells, PD1+CD4+ T-cells, γδ T-cells and macrophages, accompanied by decreased B-cells and plasma cells (**Figs. 3A, S17**). T-cells further showed increased VISTA and PD1 expression (**Fig. S17**). Among the expanded IM populations, monocytic CD11b+ IM cells expressed high levels of VISTA (**Fig. 2E**), and their frequency weakly but significantly correlated with PD1^hi^CD8+ T-cells across patients (**Fig. S18**).

Focusing on blasts, IM cells, and adaptive immune cells, force-directed network analysis confirmed co-enrichment of CD11b+ IM cells, CD8+ T-cells, PD1+CD4+ T-cells, and Tregs (**Fig. 6C**). We next assessed the prognostic relevance of specific IM–adaptive immune cell interactions (**Fig. 6D**). Higher CD11b+ IM–CD8+ T-cell co-enrichment was significantly associated with inferior survival in univariate analysis (**Fig. 6E**), potentially reflecting local suppression of antileukemic immunity. Similarly, while PD1+CD4+ T-cell frequency was not associated with OS, co-enrichment with mixed IM cells and with CD8+ T-cells predicted inferior survival. In contrast, CD15+ IM–Treg co-enrichment was associated with favorable OS (**Figs. 6E, S19**).

Because age was strongly associated with survival in our cohort (**Fig. S14A**), we examined age-related patterns in CT and CN frequencies, CT-CN enrichment, and CT-CT co-localization. While CD11b+ IM cells decreased with age, their co-enrichment with CD8+ T-cells in CN5 showed a weak positive association with age (**Fig. S20**). Furthermore, CD8+ T-cells preferentially co-localized with CD11b+ IM cells in older patients, but with blasts in younger patients. This suggests that the adverse prognostic associations of CD11b+ IM cell enrichment in CN5 and CD11b+ IM– CD8+ T-cell co-localization reflect age-associated spatial phenotypes. We therefore performed age-adjusted regression of the previously identified spatial features (**Fig. 6F**). This revealed macrophage enrichment in CN5 as a predictor of inferior OS and CD15+ IM–Treg co-enrichment as a predictor of improved OS, neither of which was associated with patient age.

To complement the cellular and spatial architecture identified by CODEX, we performed MERFISH spatial transcriptomics using an 815-gene Immuno-Oncology panel on 22 BM samples from the same AML cohort. After quality control, 38,260 cells from 18 patients were included. Unsupervised clustering identified 13 CTs, including blast subtypes, stromal cells, T-cells and IM cells. Consistent with CODEX, transcriptomics revealed an IM cell population co-expressing *ITGAM, ITGAX* and *VSIR* (CD11b, CD11c, and VISTA) and *LYZ*, *CEBPB*, *CIITA*, and *CYBB* (**Fig. S21A**), consistent with monocytic AML differentiation^34^. Ligand-receptor analysis revealed candidate interactions between CD11b+ IM cells and T-cells (**Figs. 6G-H, S21B**), including *TGFB1*–*TGFBR1* and *JAG1*–*NOTCH1*, accompanied by correlated expression of *VSIR* in CD11b+ IM cells and *PDCD1* in T-cells (**Fig. S21C**). CD11b+ IM cells also interacted with MSCs via *TGFB1*–*TGFBR2, ICAM1*– *ITGAM*, *IL6*–*IL6R*, and *CXCL12*–*CXCR4* (**Figs. 6G-H, S21D**). Together, these findings provide a potential mechanistic basis for the prognostically relevant interactions between IM and adaptive immune cells identified by CODEX.

## DISCUSSION

We established CODEX and MERFISH to characterize the BM microenvironment in routine FFPE samples of *NPM1*-mutant AML and NBM. CN clustering revealed architectural reorganization, including loss of immune-stromal interactions and emergence of AML-associated adaptive immune neighborhoods. AML showed marked expansion of phenotypically distinct IM populations with monocytic and immunosuppressive features. Importantly, spatial context-dependent enrichment of CD11b+ IM cells and macrophages with adaptive immune cells provided stronger prognostic signals than CT or CN frequencies alone. Our findings provide spatial context for previously described immunosuppressive mechanisms in AML and may inform prognostic and therapeutic stratification.

Decalcified BM poses major challenges for spatial profiling due to epitope loss and RNA degradation. Previous high-resolution studies therefore used fresh BM or restricted multiplex panels^20,21,35,36^. Using CODEX in 67 *NPM1*-mutant AML patients and 38 NBM controls, we annotated 27 CTs, including plasma cells, endothelial cells, MKs and MSCs, which are poorly captured by aspiration^35^. Consistent with previous reports^20,37^, AML showed increased MSC and CD8+ T-cell frequencies, and increased densities of CD4+ T-cells, γδ T-cells, macrophages, and endothelial cells.

CN and distance analyses revealed substantial architectural remodeling in AML, including separation of CD8+ T-cells from MSCs and osteoblasts and increased CD8+ T-cell self-association. NBM-specific myelopoietic and erythro-myeloid niches were associated with B-cells and MKs, whereas erythroid/Treg and plasma cell niches were shared between NBM and AML. The absence of AML-exclusive CNs among consensus CTs may reflect fragmentation of NBM architecture, although leukemia-specific populations were excluded from this comparison. Surprisingly, in AML, depletion of non-erythroid cells from erythroid neighborhoods was associated with adverse survival. This may reflect functional disruption, e.g., by leukemia-associated cytokines rather than simple displacement, consistent with leukemia-mediated suppression of erythroposiesis^38,39^ and potentially altered local oxygen and HIF-1α signaling^40–43^. AML establishes an immunosuppressive BM microenvironment that promotes leukemogenesis, treatment resistance, and relapse^16–18,44–46^. Dysfunctional Treg, γδ T-cell, and CD8+ T-cell phenotypes have been linked to chemotherapy response^17,47^, while immunosuppressive transcriptional programs predict early relapse in predominantly *NPM1*-mutant AML^19^. Additional immune evasion mechanisms include AML-induced M2 macrophage polarization^48^, immune checkpoint signaling^48–51^, and monocytic differentiation programs associated with treatment resistance^10,34,52^. In particular, CD11b+ VISTA+ monocytic myeloid-derived suppressor cells suppress anti-leukemic T-cell immunity, and their VISTA expression correlates with PD-1 expression on T-cells^53–57^.

Our spatial transcriptomic data identified a monocytic population corresponding to CODEX-defined CD11b+ IM cells and recapitulated the VISTA(*VISR*)–PD-1(*PDCD1*) relationship between CD11b+ IM and T-cells. Importantly, ligand-receptor analysis further implicated *TGFB1*–*TGFBR1* and *JAG1*–*NOTCH1* signaling. TGF-β suppresses cytotoxic T-cell function and contributes to chemoresistance^58–60^, while Jagged-1–Notch signaling has been linked to immune evasion, Treg induction, and tumorigenesis^61–63^. Notch signaling can also induce CD11b+ myeloid-derived suppressor cells^64^, supporting reciprocal interactions within the leukemic BM microenvironment.

Importantly, the adverse prognostic association of CD11b+VISTA+ IM cells depended on their spatial co-enrichment with exhausted T-cells rather than their overall frequency. Thus, our findings may represent an *in situ* correlate of previously described monocytic differentiation and immune-evasion states and suggest that their clinical impact depends on spatial organization.

Spatial studies in myeloid neoplasms remain limited and have largely focused on specific treatment settings^23–26,65^. CODEX analysis linked donor-lymphocyte infusion responses to spatial coordination of anti-leukemic CD8+ T-cell responses, while spatial transcriptomics associated PD-1 inhibitor response with altered blast–myeloid–T-cell interactions^24^.

In *TP53*-mutant AML residual disease, spatial proteomics similarly demonstrated niche-level immunosuppressive interactions not captured by bulk or single-cell analysis^25^. More recently, spatial profiling across diagnostic AML identified interactions among immunosuppressive myeloid progenitors, macrophages, and T-cells^66^. Our findings extend these observations to diagnostic *NPM1*-mutant AML, demonstrating that suppressor–immune interactions are spatially organized at diagnosis and carry prognostic relevance.

Our study has several limitations. FFPE immunostaining and nucleus-based segmentation are susceptible to nonspecific binding, variable detection, imperfect capture of irregular cell shapes, and doublets or signa-bleedover -mitigated, but not eliminated, by computational filtering and manual annotation. Spatial transcriptomics was additionally limited by RNA degradation in archival decalcified FFPE tissue, resulting in low transcript detection and reduced resolution of cell type and interaction analyses. TMA-based profiling enabled high throughput but restricted analysis to 1-mm tissue regions. Finally, the survival associations are hypothesis-generating and require prospective validation. Nevertheless, CODEX is substantially more scalable and cost-efficient than spatial transcriptomics, supporting its potential clinical translation.

In conclusion, diagnostic *NPM1*-mutant AML is genetically defined but phenotypically and microenvironmentally heterogeneous. Spatial single-cell proteomics transforms routine BM biopsies into quantitative maps of leukemic differentiation, residual hematopoiesis, immune suppression, and neighborhood remodeling that provide information beyond genotype and cell abundance. In particular, the spatial association of VISTA+ monocytic IM cells with PD1+ T-cells may help explain the limited efficacy of bulk checkpoint blockade and support spatially informed combination strategies. Beyond providing a spatial atlas of *NPM1*-mutant AML, our approach establishes a framework for translating spatial biomarker discovery in routine FFPE biopsies into scalable phenotyping for prognostication and treatment guidance.

## Acknowledgements

We thank the patients and their families for their informed consent and donations of tissues. C.M.S. was supported by the Faculty of Medicine, University of Tübingen; the Institute of Pathology, University Hospital Tübingen; the State of Baden-Württemberg within the Centers for Personalized Medicine Baden-Württemberg (ZPM); the Cluster of Excellence iFIT (EXC 2180) ‘Image-Guided and Functionally Instructed Tumor Therapies’, University of Tübingen; the Mach-Gaensslen-Stiftung Schweiz; the Dres. Bayer-Foundation; Swiss Cancer Research (KFS-5114-08-2020); the European Union (ERC, 101116768; MSCA, 101226364); the German Research Foundation (EXC 2180 -390900677; INST 37/1228-1 FUGG; INST 37/1302-1 FUGG; INST 37/1310-1 FUGG; iRTG 101226364); the Brigitte und Dr. Konstanze Wegener-Stiftung; the Werner Jackstädt-Stiftung; the Bundesinstitut für Risikobewertung (Bf3R, 60-0102-01.P641-12572660); the Wilhelm Sander Foundation (2024.096.1; 2025.216.1), the German Cancer Aid (70117323; 70117411), the Rolf M. Schwiete Foundation (2025-076), the Alternatives Research & Development Foundation (1650881), the German Academic Exchange Service (Postdoc-NeT-AI), and the Leukemia Research Foundation (1077776). J.C.S. was supported by the MINT-Clinician Scientist program of the Faculty of Medicine, University of Tübingen, funded by the German Research Foundation (493665037); by a Walter Benjamin Fellowship from the German Research Foundation (557326522); and by the Werner Jackstädt-Stiftung. J.Z. was supported by the China Scholarship Council (202108370099). C.L. was supported by the European Union (ERC, 866548) and the German Research Foundation (EXC 2180 – 390900677; 566352518).

## Authorship

J.C.S., C.L. and C.M.S. conceptualized the study; J.C.S., P.Z., B.R.S., M.Z., L.L., and C.B. performed the methodology; J.C.S., P.Z., B.R.S., M.Z., and L.L. performed the formal analysis; J.C.S., P.Z., B.R.S., and M.Z. performed the investigation; J.C.S., P.Z., B.R.S., M.Z., A.M., A.R., J.Z., Y.A.P., L.M., W.A.B., C.L., and C.M.S. were responsible for the resources; P.Z., J.C.S., B.R.S., M.Z., and L.M. performed the data curation; J.C.S. wrote the original draft; J.C.S., P.Z., B.R.S., and M.Z. visualized the study; C.L. and C.M.S. supervised the study; J.C.S. and C.M.S. performed the project administration; J.C.S., C.L., and C.M.S. acquired funding; and all authors reviewed and edited the manuscript.

## Disclosure of Conflicts of Interest

C.M.S. is a cofounder, shareholder and employee of Vicinity Bio GmbH, and is a scientific advisor to Enable Medicine Inc., all outside the current work. The other authors declare that no conflicts of interest exist.

