## Supplemental Figures and Tables for "Spatial co-enrichment of immature myeloid and T-cell populations predicts survival in NPM1-mutant AML"

##### Figure S1

Normal BM TMA 1

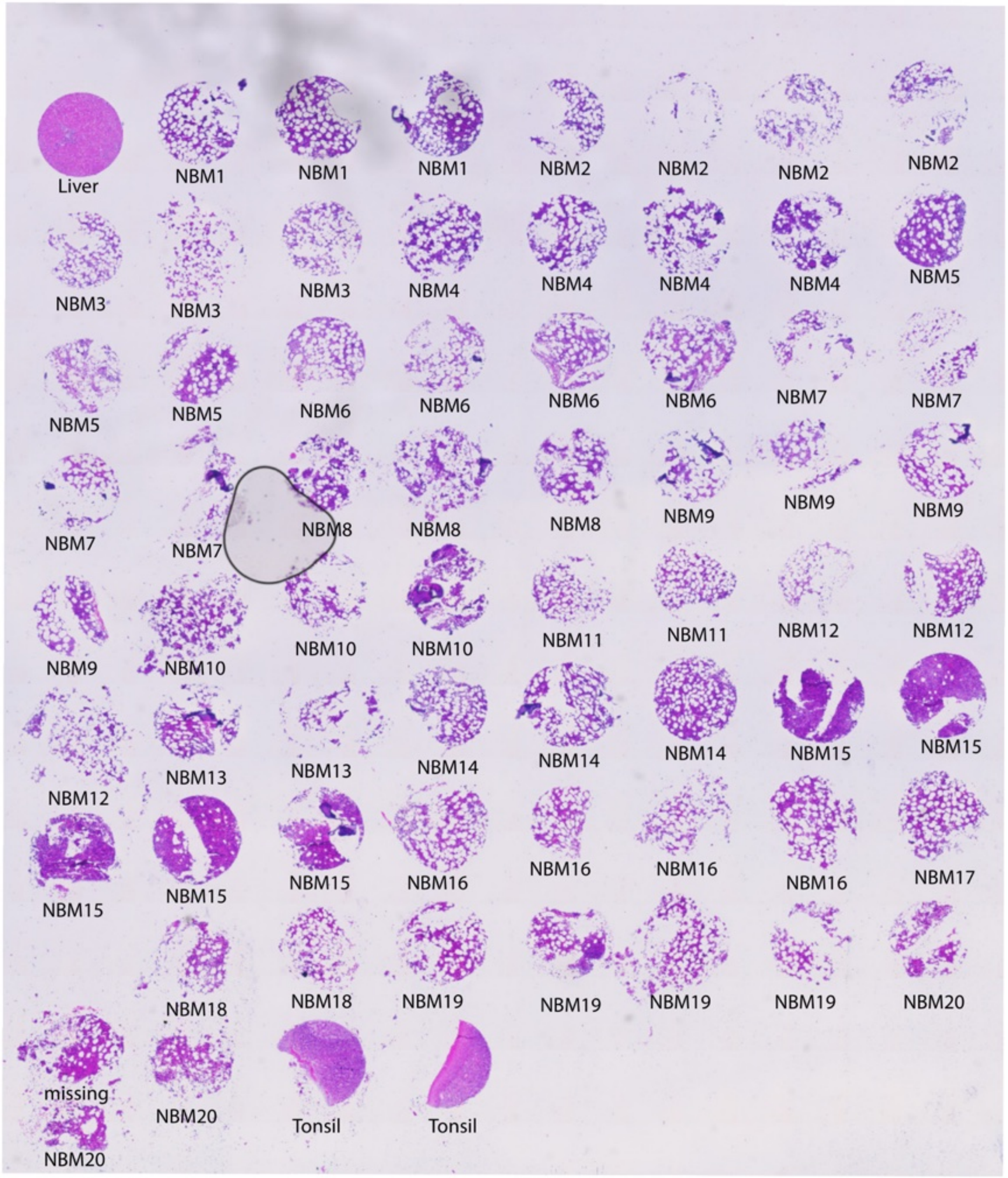

Normal BM TMA 2

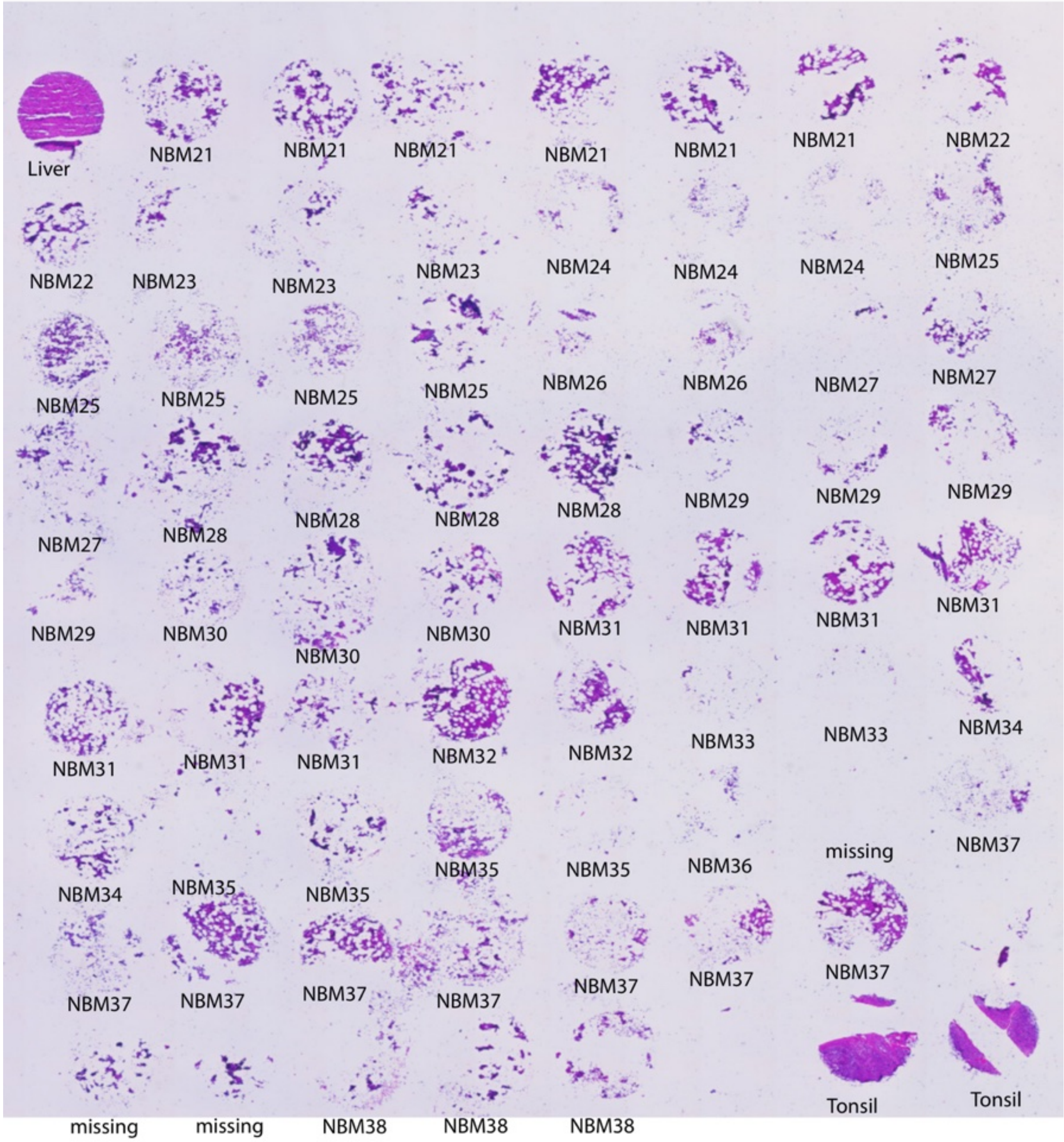

NPM1-mutant AML TMA 1

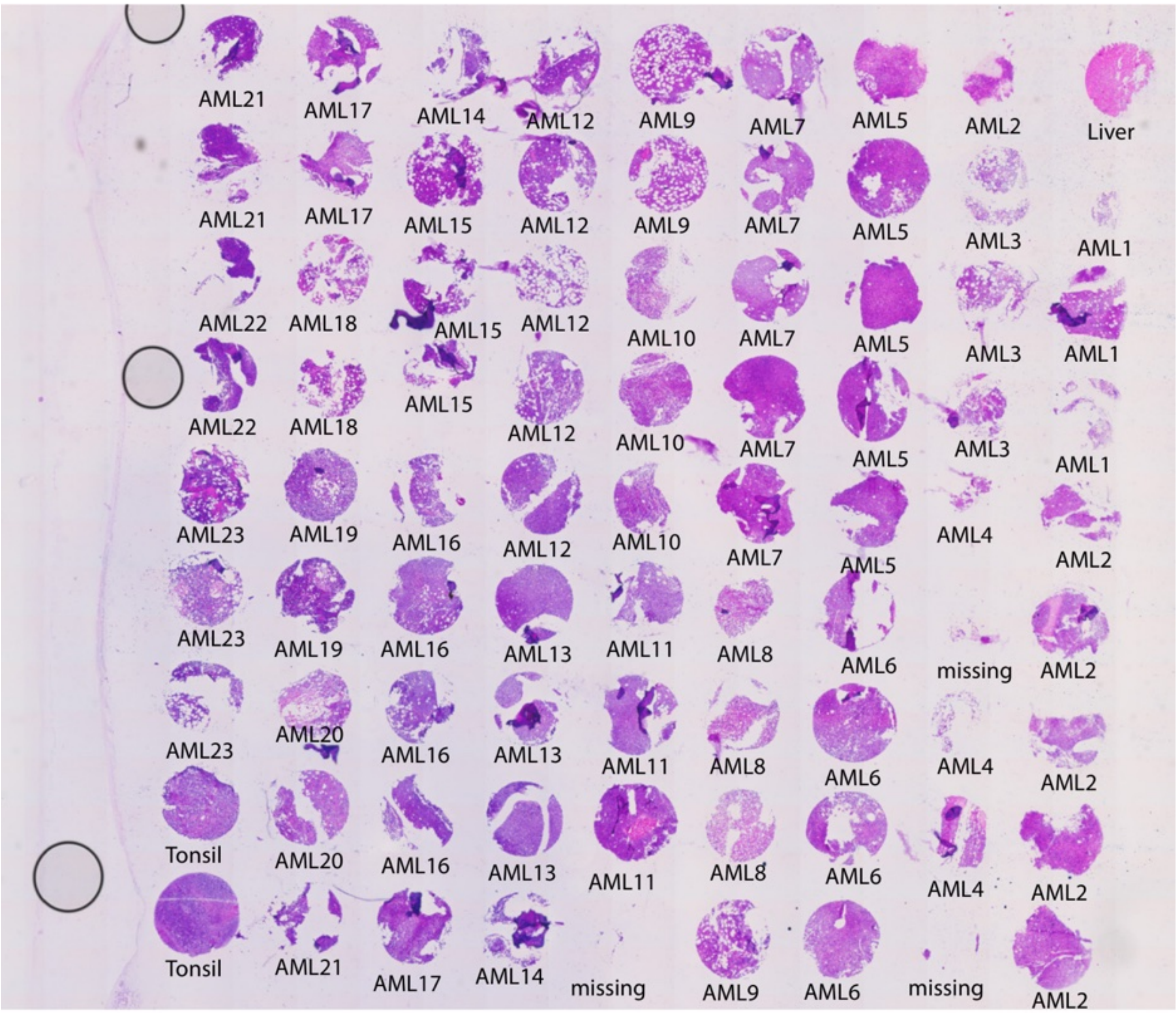

NPM1-mutant AML TMA 2

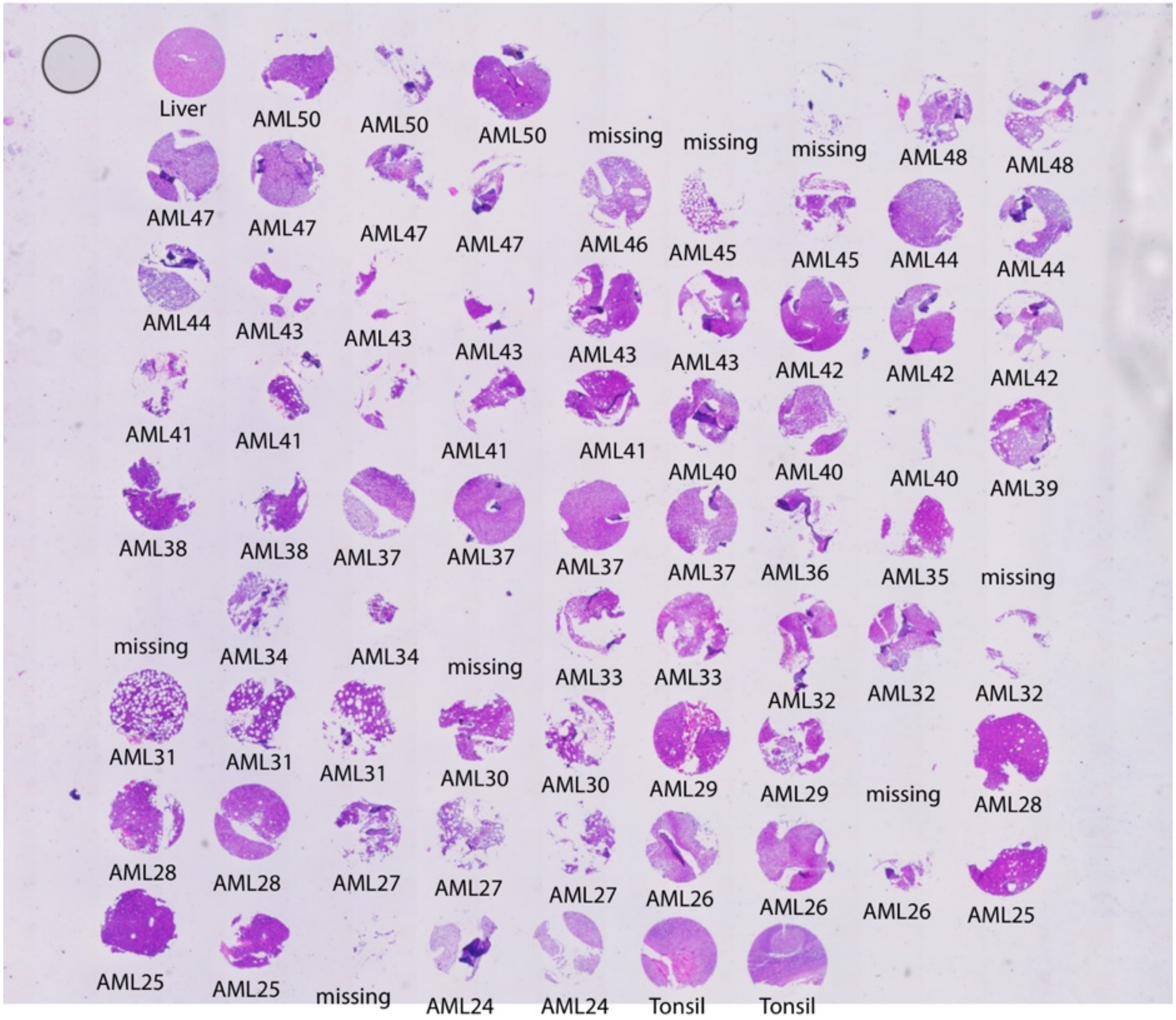

NPM1-mutant AML TMA 3

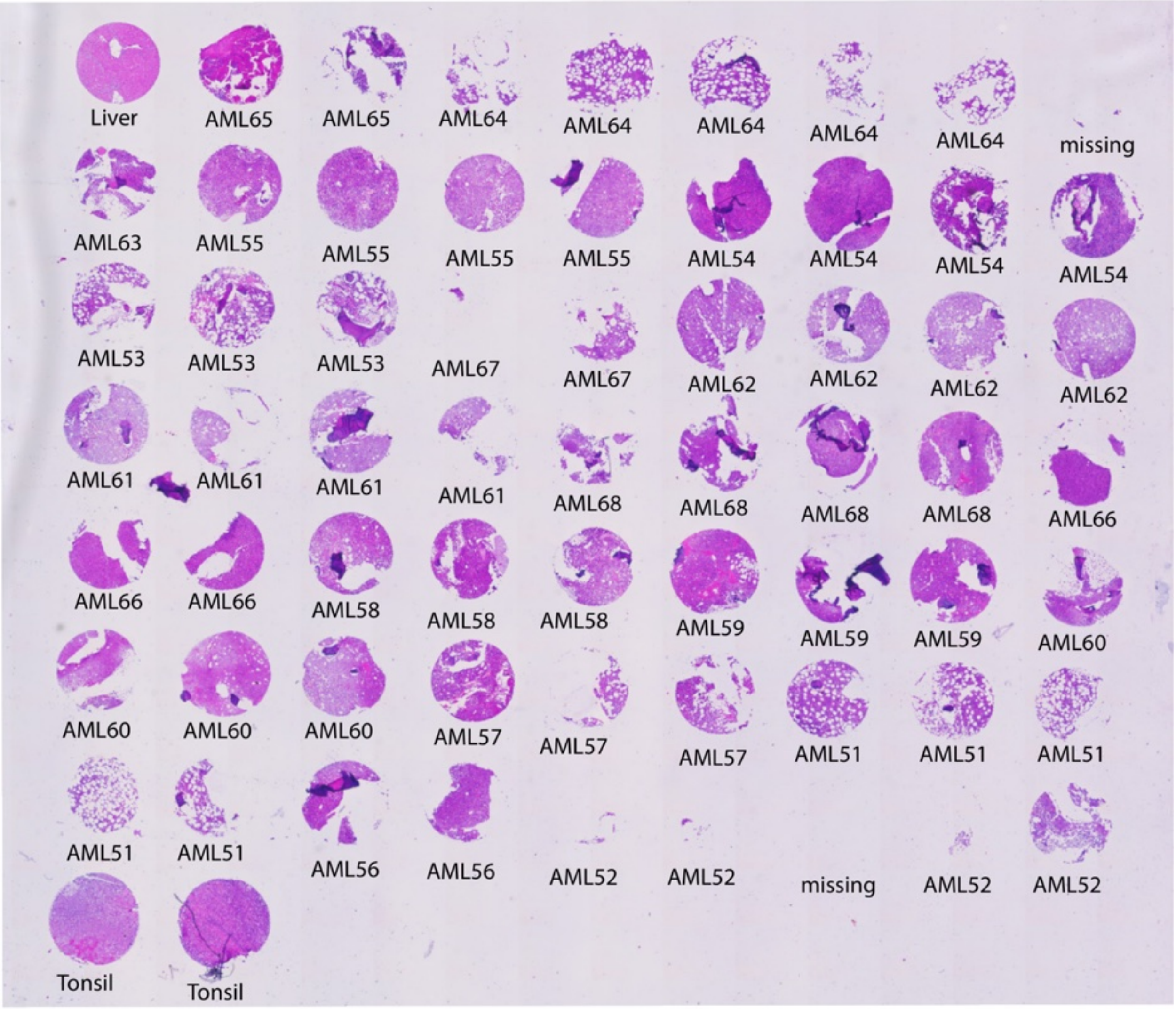


**Figure S1: Overview of H&E-stained tissue microarrays (TMAs).**

Five TMAs were H&E-stained and scanned after CODEX imaging had been performed. Corresponding normal bone marrow (NBM) cohort individual and AML patient IDs are annotated. TMA cores missing in the CODEX imaging dataset due to tissue detachment or insufficient image resolution are indicated.

**Figure S2**

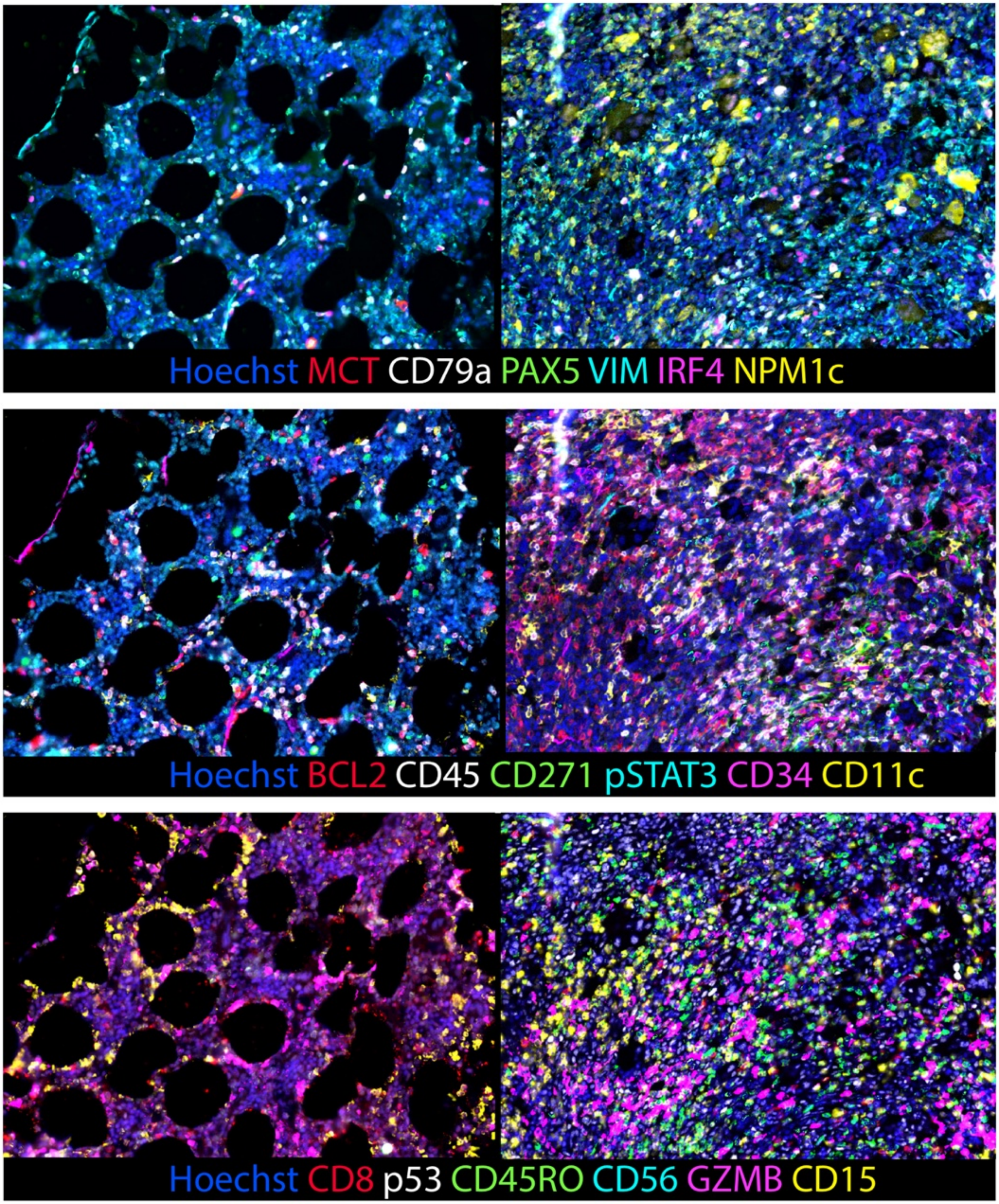
Hoechst MCT CD79a PAX5 VIM IRF4 NPM1c
Hoechst BCL2 CD45 CD271 pSTAT3 CD34 CD11c
Hoechst CD8 p53 CD45RO CD56 GZMB CD15

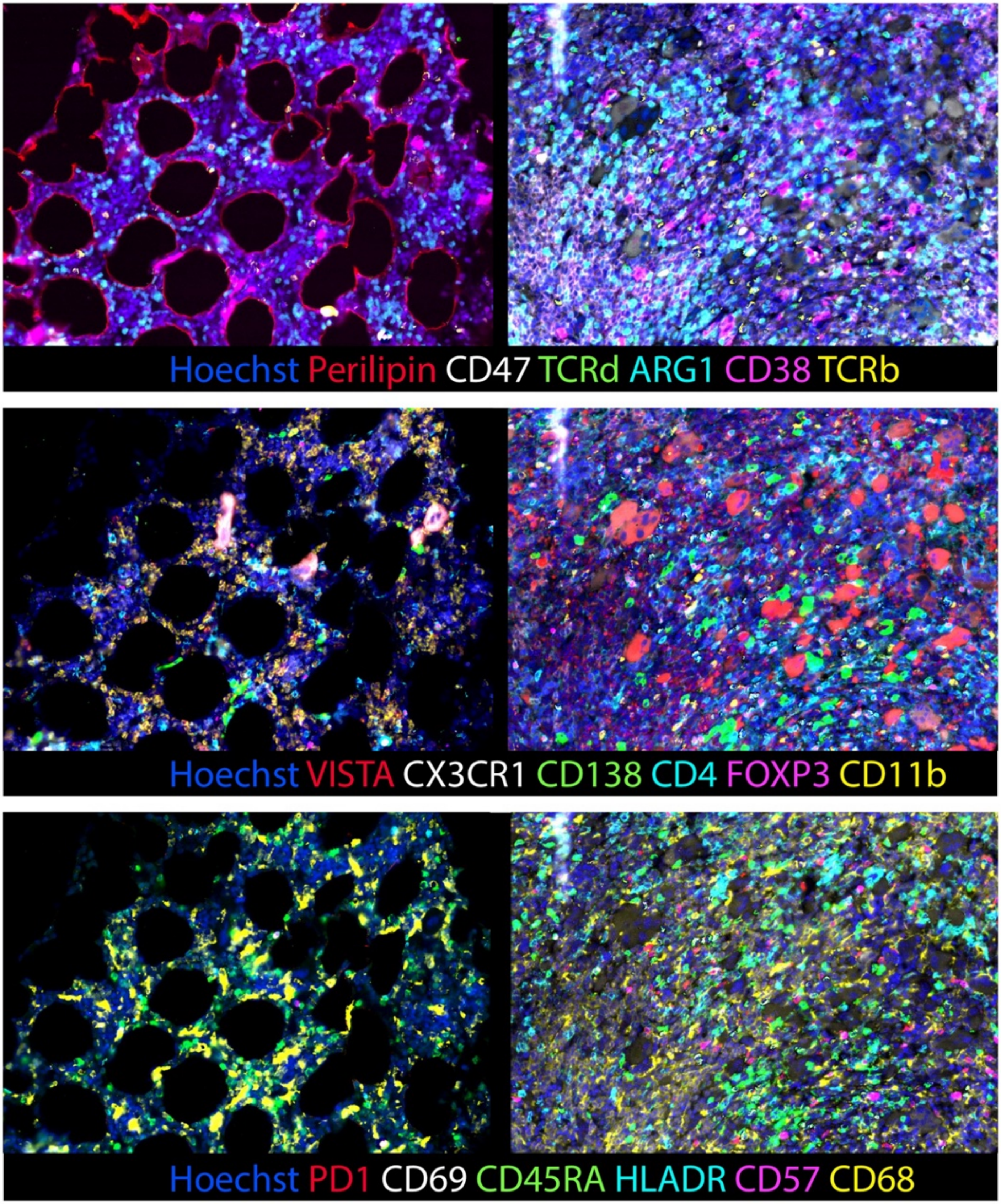
Hoechst Perilipin CD47 TCRd ARG1 CD38 TCRb
Hoechst VISTA CX3CR1 CD138 CD4 FOXP3 CD11b
Hoechst PD1 CD69 CD45RA HLADR CD57 CD68

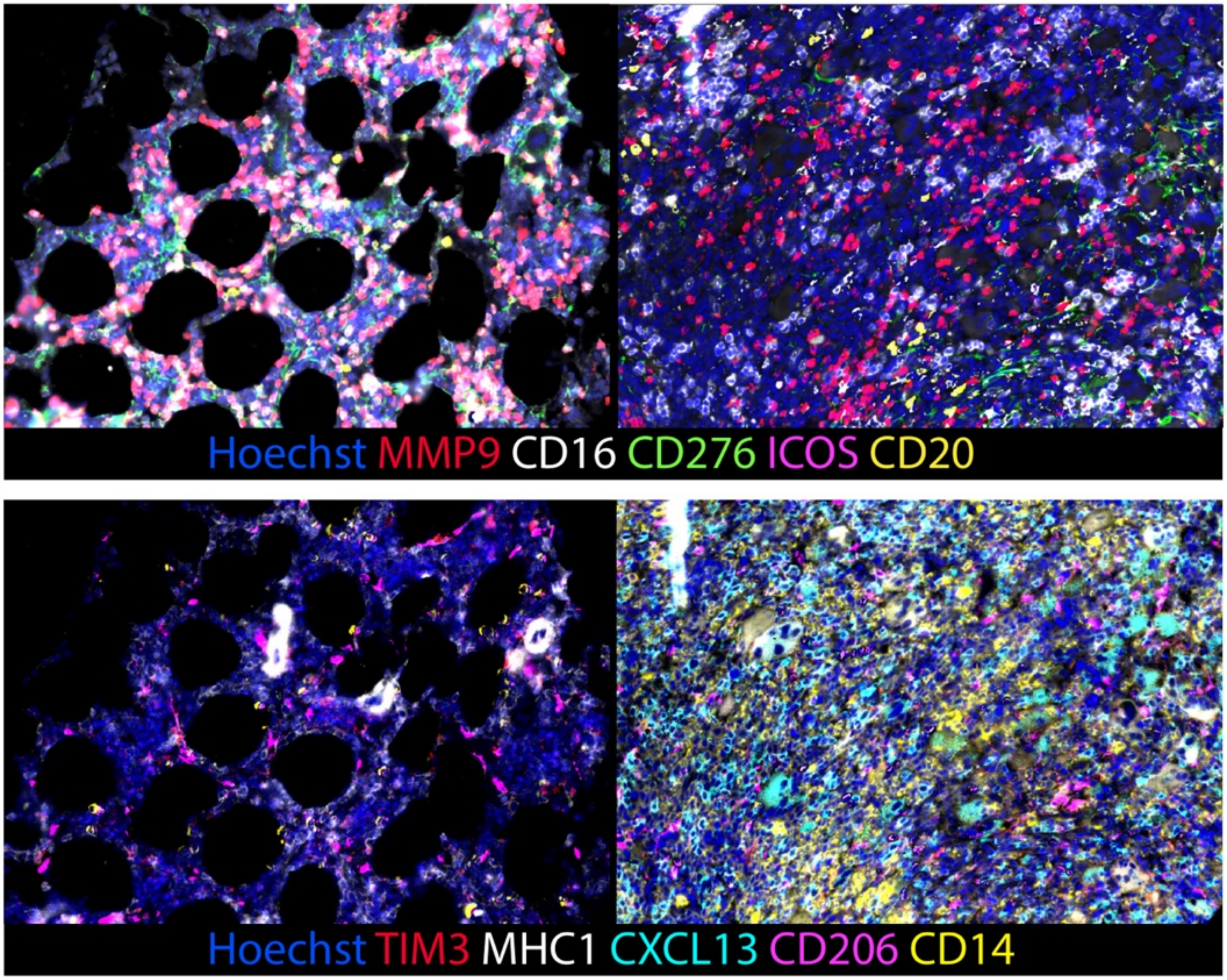
Hoechst MMP9 CD16 CD276 ICOS CD20
Hoechst TIM3 MHC1 CXCL13 CD206 CD14

**Figure S2: 7-marker CODEX overlays for all markers.**

CODEX overlay images showing 7 markers across all markers in the panel excluding the ones shown in Figure 1E, on exemplary normal bone marrow (NBM; image panels on the left) and AML (image panels on the right) regions.

#### Figure S3

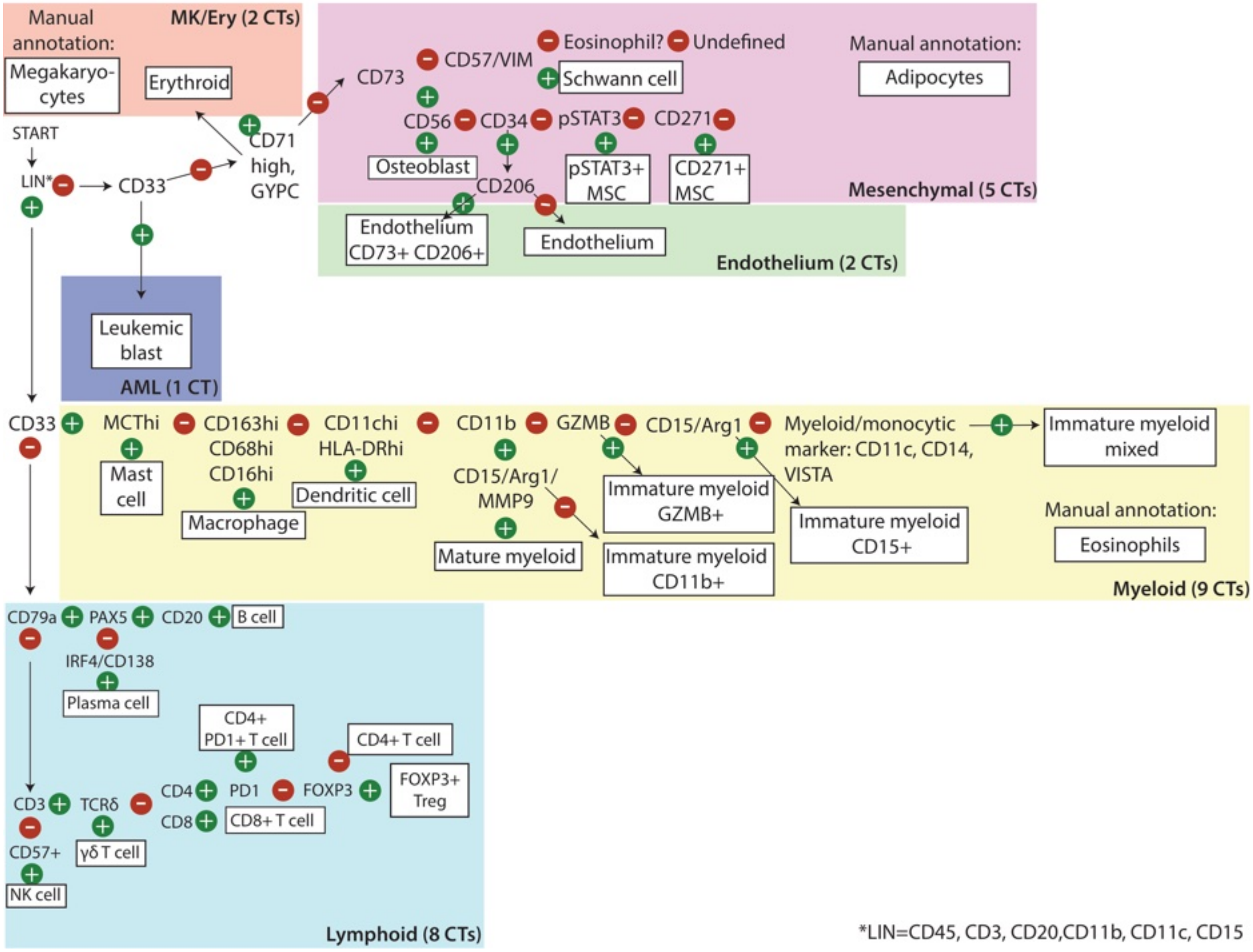

Manual annotation:
MK/Ery (2 CTs)
Megakaryo-cytes
Erythroid
START
LIN*
CD33
CD71 high, GYPC
Leukemic blast
AML (1 CT)
CD73
CD57/VIM
Eosinophil?
Undefined
Schwann cell
Manual annotation:
Adipocytes
CD56
CD34
pSTAT3
CD271
Osteoblast
pSTAT3+ MSC
CD271+ MSC
CD206
Mesenchymal (5 CTs)
Endothelium CD73+ CD206+
Endothelium
Endothelium (2 CTs)
CD33
MCThi
CD163hi CD68hi CD16hi
CD11chi HLA-DRhi
CD11b
GZMB
CD15/Arg1
Myeloid/monocytic marker: CD11c, CD14, VISTA
Immature myeloid mixed
Mast cell
Macrophage
Dendritic cell
CD15/Arg1/ MMP9
Immature myeloid GZMB+
Immature myeloid CD15+
Manual annotation:
Eosinophils
Mature myeloid
Immature myeloid CD11b+
Myeloid (9 CTs)
CD79a
PAX5
CD20
B cell
IRF4/CD138
Plasma cell
CD4+ PD1+ T cell
CD4+ T cell
CD3
TCRδ
CD4
PD1
FOXP3
FOXP3+ Treg
CD8
CD8+ T cell
CD57+
γδ T cell
NK cell
Lymphoid (8 CTs)
*LIN=CD45, CD3, CD20,CD11b, CD11c, CD15

**Figure S3: Cell type annotation rationale.**

Flow chart depicting cell type (CT) annotation rationale based on marker expression. CTs are depicted in white boxes and grouped by lineage (colored boxes).

**Figure S4**

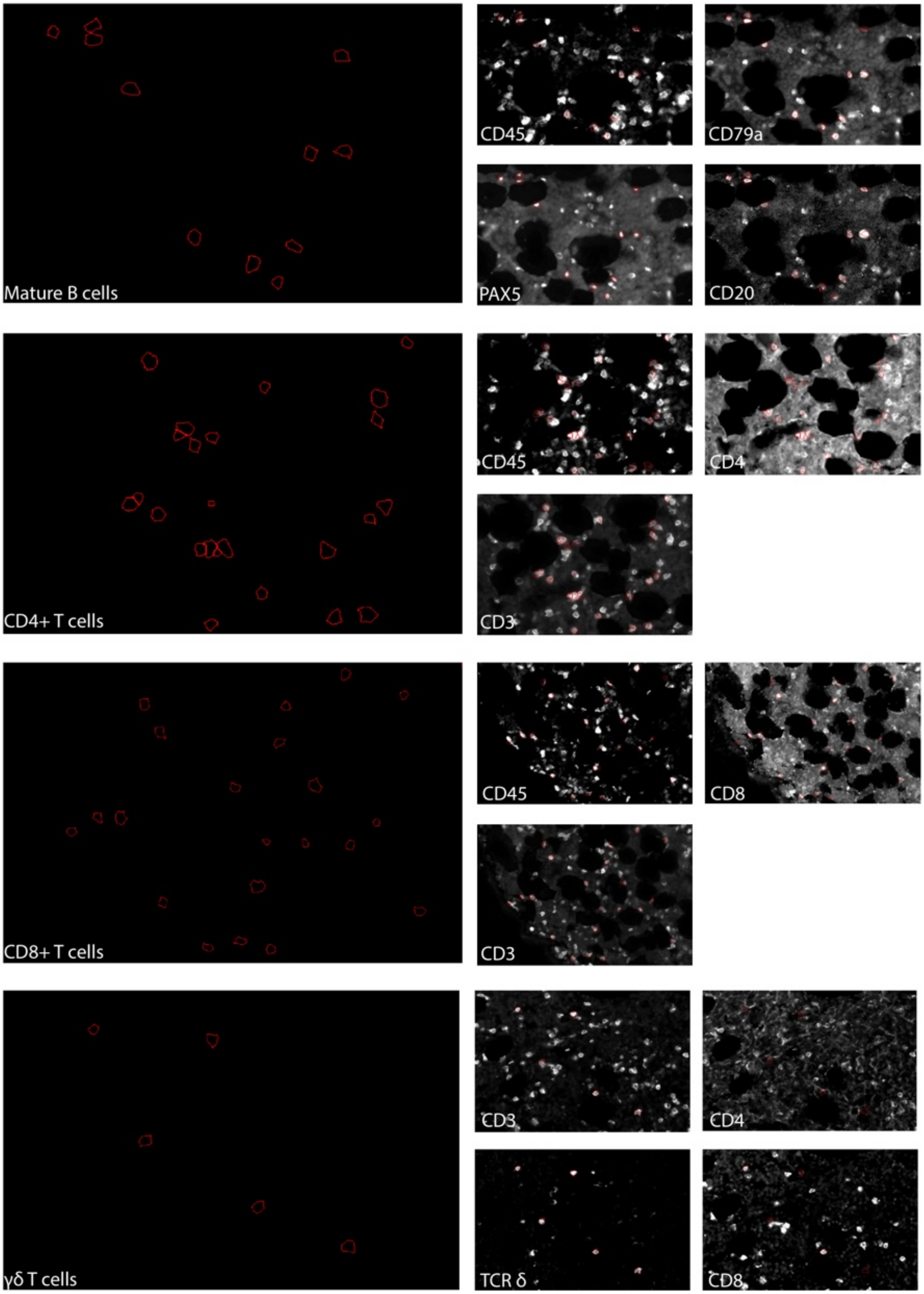
Mature B cells
CD45
CD79a
PAX5
CD20
CD4+ T cells
CD45
CD4
CD3
CD8+ T cells
CD45
CD8
CD3
γδ T cells
CD3
CD4
TCR δ
CD8

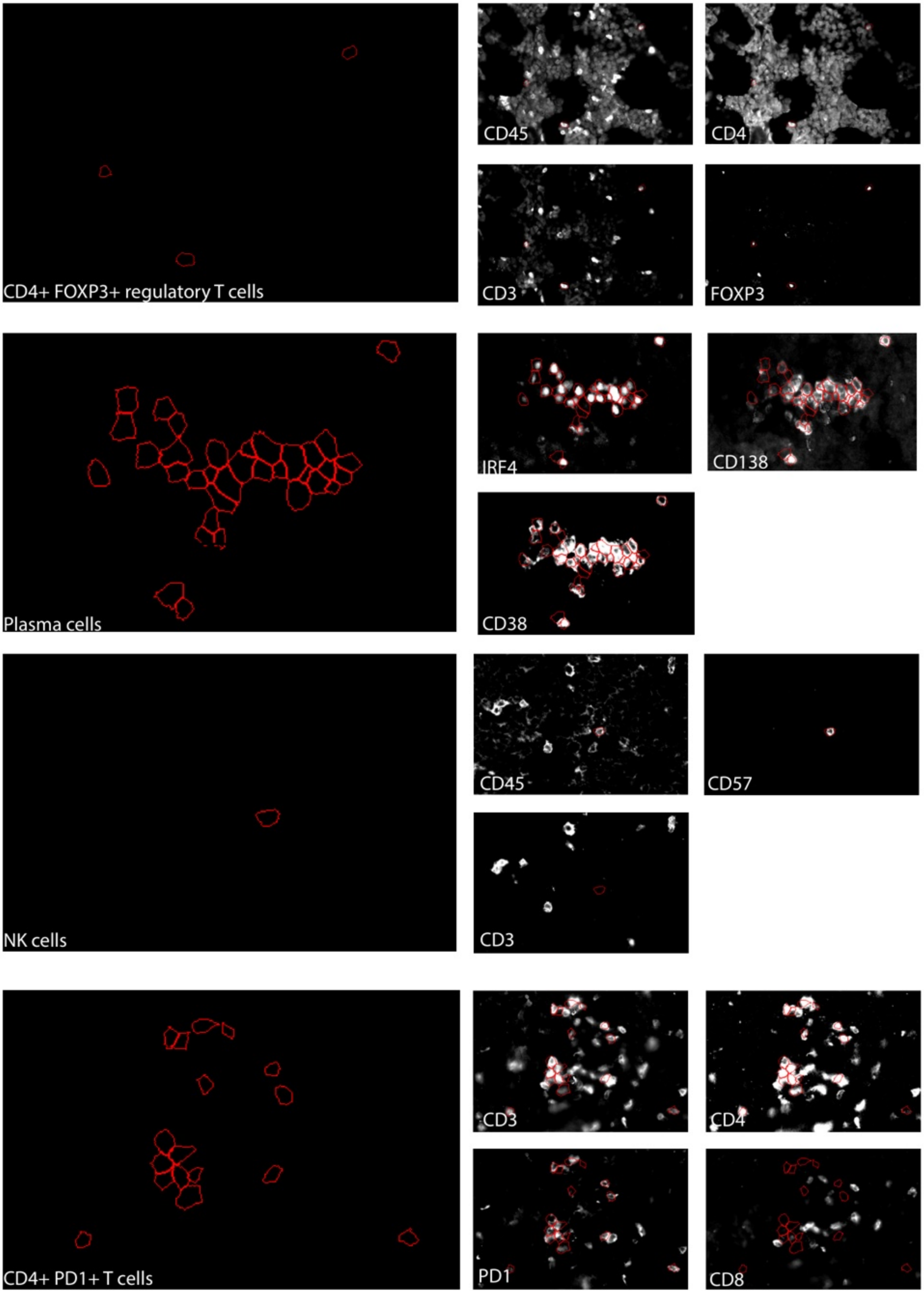
CD4+ FOXP3+ regulatory T cells
CD45
CD4
CD3
FOXP3
Plasma cells
IRF4
CD138
CD38
NK cells
CD45
CD57
CD3
CD4+ PD1+ T cells
CD3
CD4
PD1
CD8

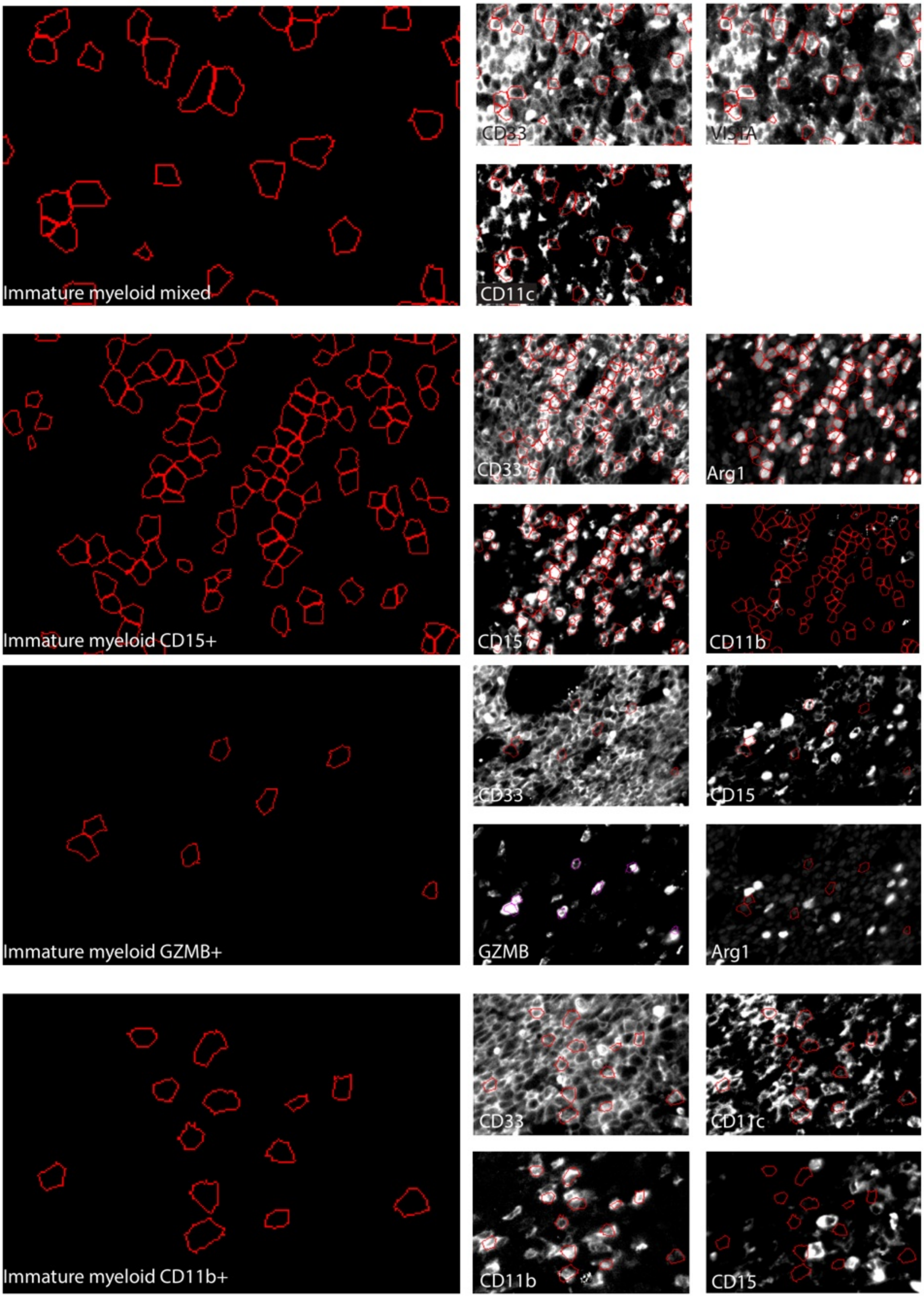
Immature myeloid mixed
CD33
VISTA
CD11c
Immature myeloid CD15+
CD33
Arg1
CD15
CD11b
Immature myeloid GZMB+
CD33
CD15
GZMB
Arg1
Immature myeloid CD11b+
CD33
CD11c
CD11b
CD15

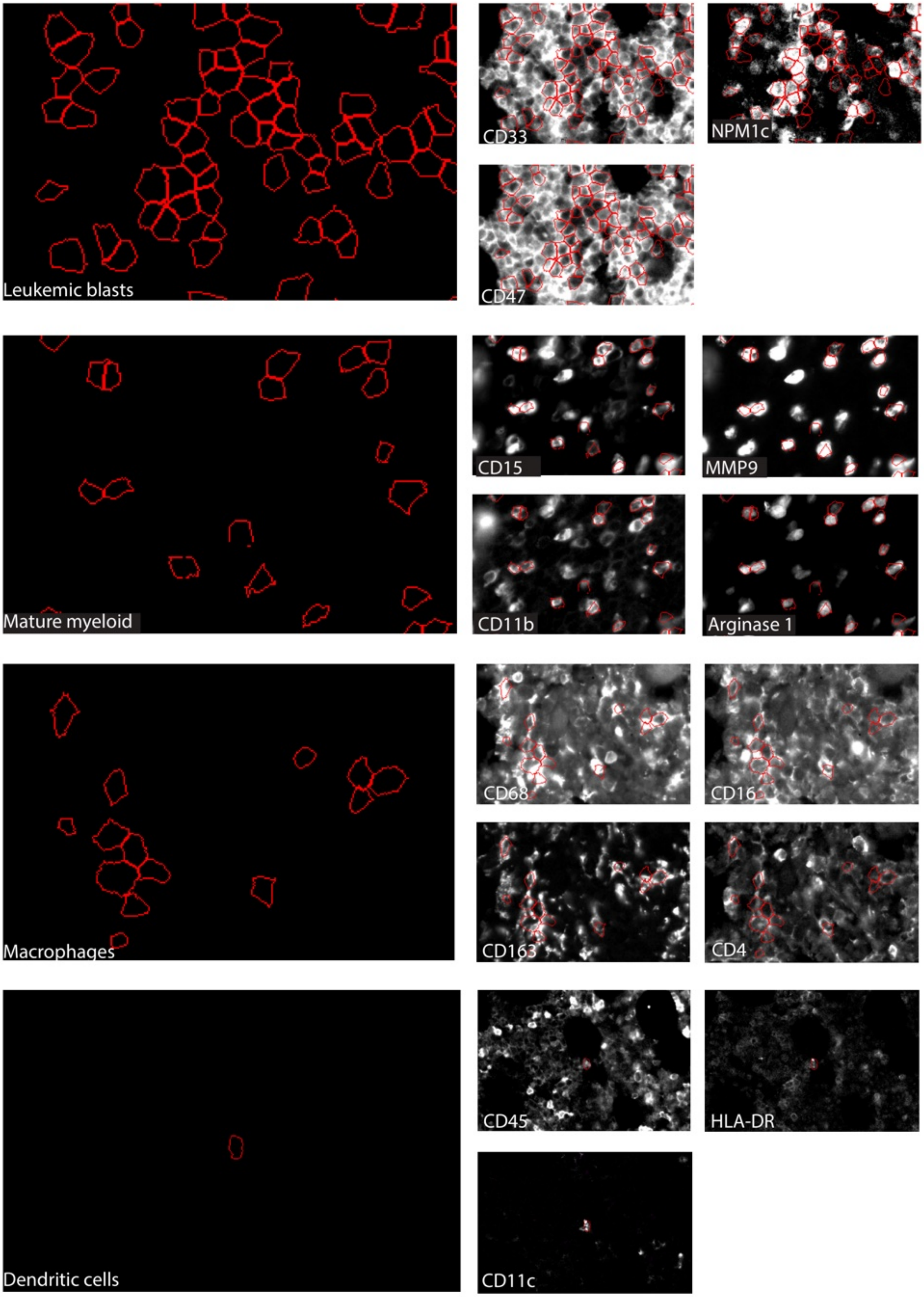
Leukemic blasts
CD33
NPM1c
CD47
Mature myeloid
CD15
MMP9
CD11b
Arginase 1
Macrophages
CD68
CD16
CD163
CD4
Dendritic cells
CD45
HLA-DR
CD11c

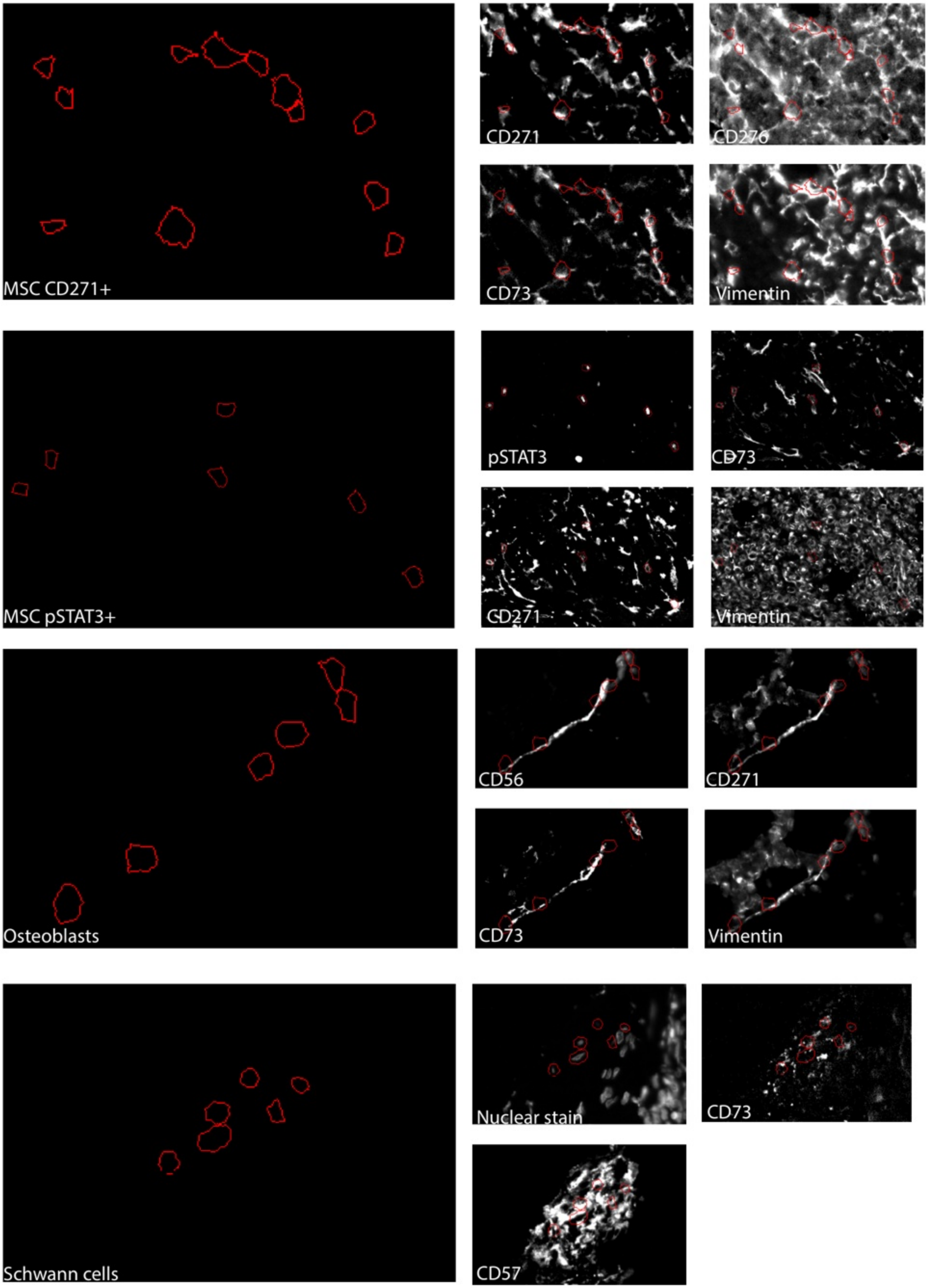
MSC CD271+
CD271
CD276
CD73
Vimentin
MSC pSTAT3+
pSTAT3
CD73
CD271
Vimentin
Osteoblasts
CD56
CD271
CD73
Vimentin
Schwann cells
Nuclear stain
CD73
CD57

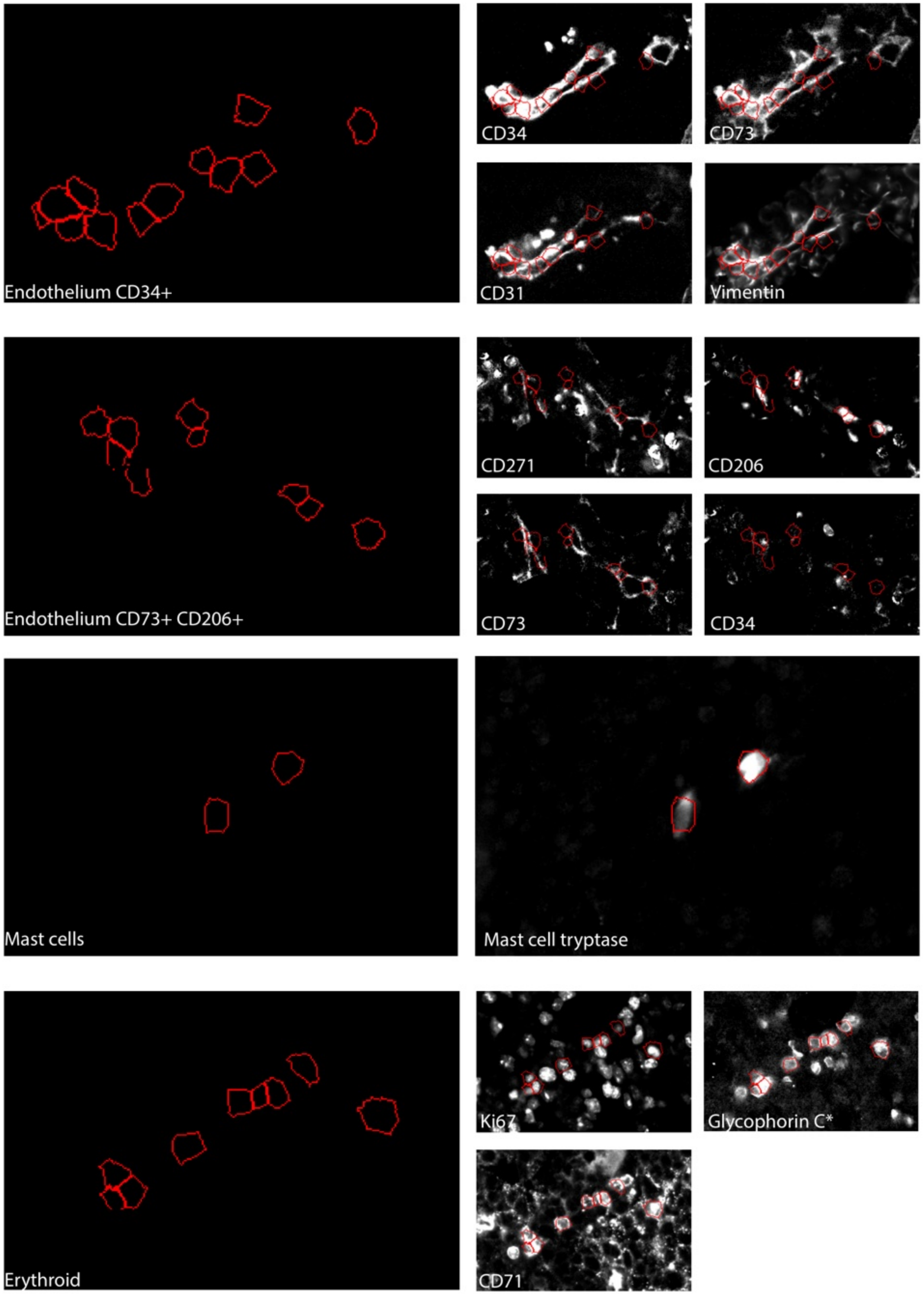
Endothelium CD34+
CD34
CD73
CD31
Vimentin
Endothelium CD73+ CD206+
CD271
CD206
CD73
CD34
Mast cells
Mast cell tryptase
Erythroid
Ki67
Glycophorin C*
CD71

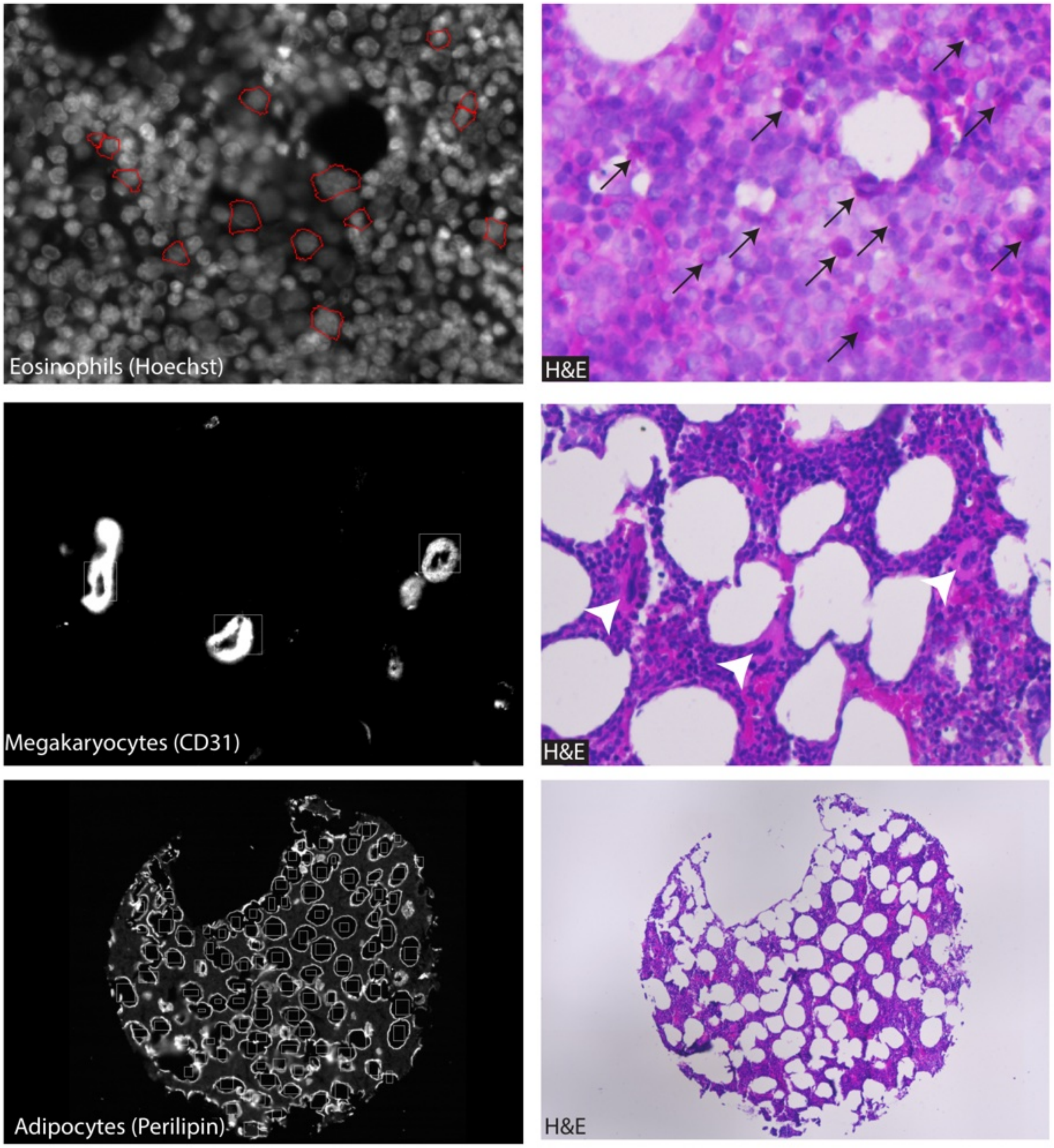

Eosinophils (Hoechst)
H&E
Megakaryocytes (CD31)
H&E
Adipocytes (Perilipin)
H&E

**Figure S4: Cell type annotation examples.**

Representative single-color images for cell type specific markers and H&E images used for annotation. Eosinophil clusters were identified by correlation with H&E, see example (eosinophils indicated by black arrows). Megakaryocytes (white arrow heads) and adipocytes were manually annotated by correlation with markers (CD31, perilipin) and with H&E.

#### Figure S5

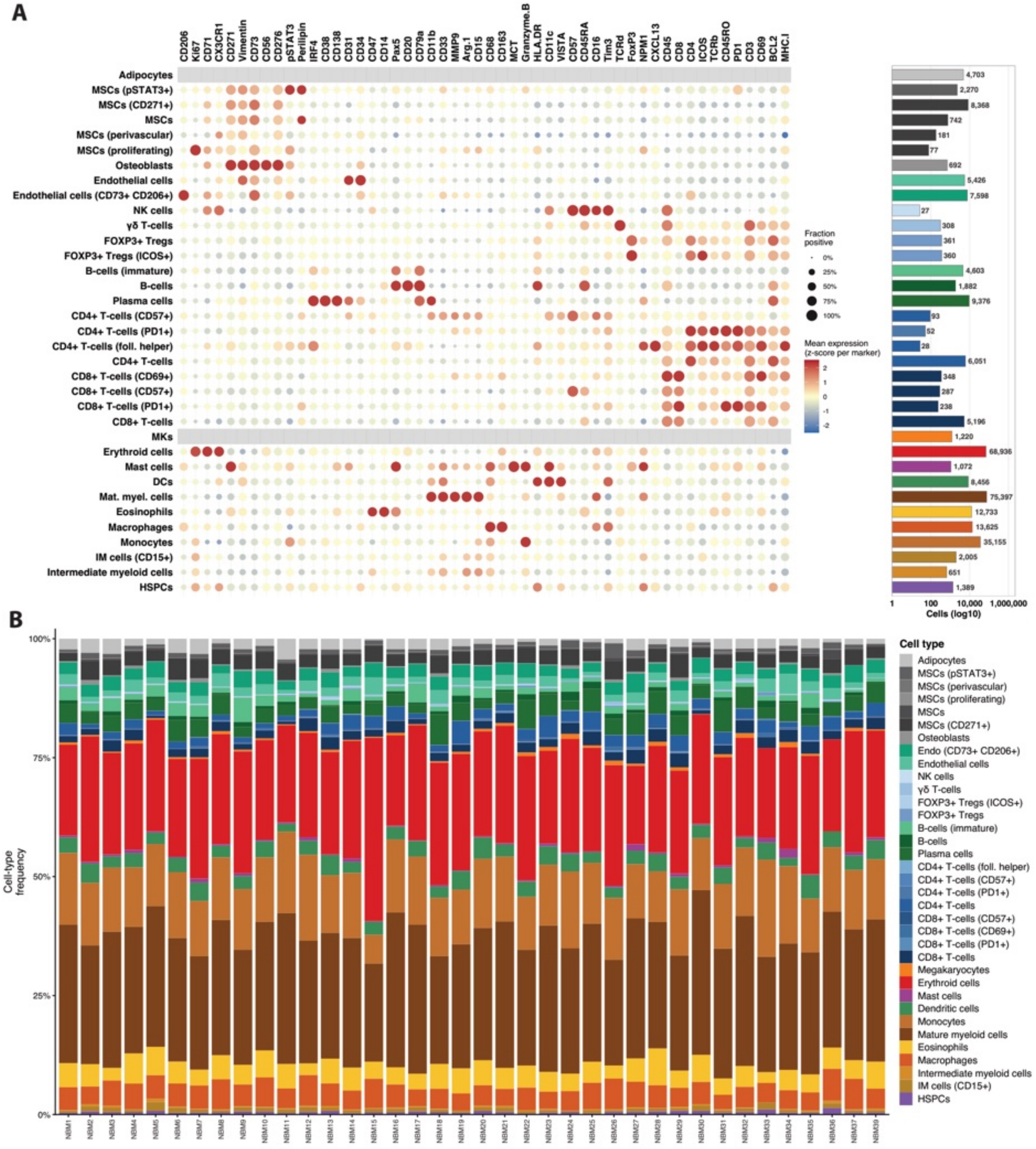
A
CD206
Ki67
CD71
CX3CR1
CD271
Vimentin
CD73
CD56
CD276
pSTAT3
Perilipin
IRF4
CD38
CD138
CD31
CD34
CD47
CD14
Pax5
CD20
CD79a
CD11b
CD33
MMP9
Arg.1
CD15
CD68
CD163
MCT
Granzyme.B
HLA.DR
CD11c
VISTA
CD57
CD45RA
CD16
Tim3
TCRd
FoxP3
NPM1
CXCL13
CD45
CD8
CD4
ICOS
TCRb
CD45RO
PD1
CD3
CD69
BCL2
MHC.I
Adipocytes
MSCs (pSTAT3+)
MSCs (CD271+)
MSCs
MSCs (perivascular)
MSCs (proliferating)
Osteoblasts
Endothelial cells
Endothelial cells (CD73+ CD206+)
NK cells
γδ T-cells
FOXP3+ Tregs
FOXP3+ Tregs (ICOS+)
B-cells (immature)
B-cells
Plasma cells
CD4+ T-cells (CD57+)
CD4+ T-cells (PD1+)
CD4+ T-cells (foll. helper)
CD4+ T-cells
CD8+ T-cells (CD69+)
CD8+ T-cells (CD57+)
CD8+ T-cells (PD1+)
CD8+ T-cells
MKs
Erythroid cells
Mast cells
DCs
Mat. myel. cells
Eosinophils
Macrophages
Monocytes
IM cells (CD15+)
Intermediate myeloid cells
HSPCs
Fraction positive
0%
25%
50%
75%
100%
Mean expression (z-score per marker)
2
1
0
-1
-2
4,703
2,270
8,368
742
181
77
692
5,426
7,598
27
308
361
360
4,603
1,882
9,376
93
52
28
6,051
348
287
238
5,196
1,220
68,936
1,072
8,456
75,397
12,733
13,625
35,155
2,005
651
1,389
1
100
10,000
1,000,000
Cells (log10)
B
100%
75%
50%
25%
0%
Cell-type frequency
NBM1
NBM2
NBM3
NBM4
NBM5
NBM6
NBM7
NBM8
NBM9
NBM10
NBM11
NBM12
NBM13
NBM14
NBM15
NBM16
NBM17
NBM18
NBM19
NBM20
NBM21
NBM22
NBM23
NBM24
NBM25
NBM26
NBM27
NBM28
NBM29
NBM30
NBM31
NBM32
NBM33
NBM34
NBM35
NBM36
NBM37
NBM39
Cell type
Adipocytes
MSCs (pSTAT3+)
MSCs (perivascular)
MSCs (proliferating)
MSCs
MSCs (CD271+)
Osteoblasts
Endo (CD73+ CD206+)
Endothelial cells
NK cells
γδ T-cells
FOXP3+ Tregs (ICOS+)
FOXP3+ Tregs
B-cells (immature)
B-cells
Plasma cells
CD4+ T-cells (foll. helper)
CD4+ T-cells (CD57+)
CD4+ T-cells (PD1+)
CD4+ T-cells
CD8+ T-cells (CD57+)
CD8+ T-cells (CD69+)
CD8+ T-cells (PD1+)
CD8+ T-cells
Megakaryocytes
Erythroid cells
Mast cells
Dendritic cells
Monocytes
Mature myeloid cells
Eosinophils
Macrophages
Intermediate myeloid cells
IM cells (CD15+)
HSPCs

**Figure S5: Cell type annotation for normal bone marrow cohort.**

A: Marker expression per cell type. B: Stacked bar chart depicting cell type frequencies for each individual.

#### Figure S6

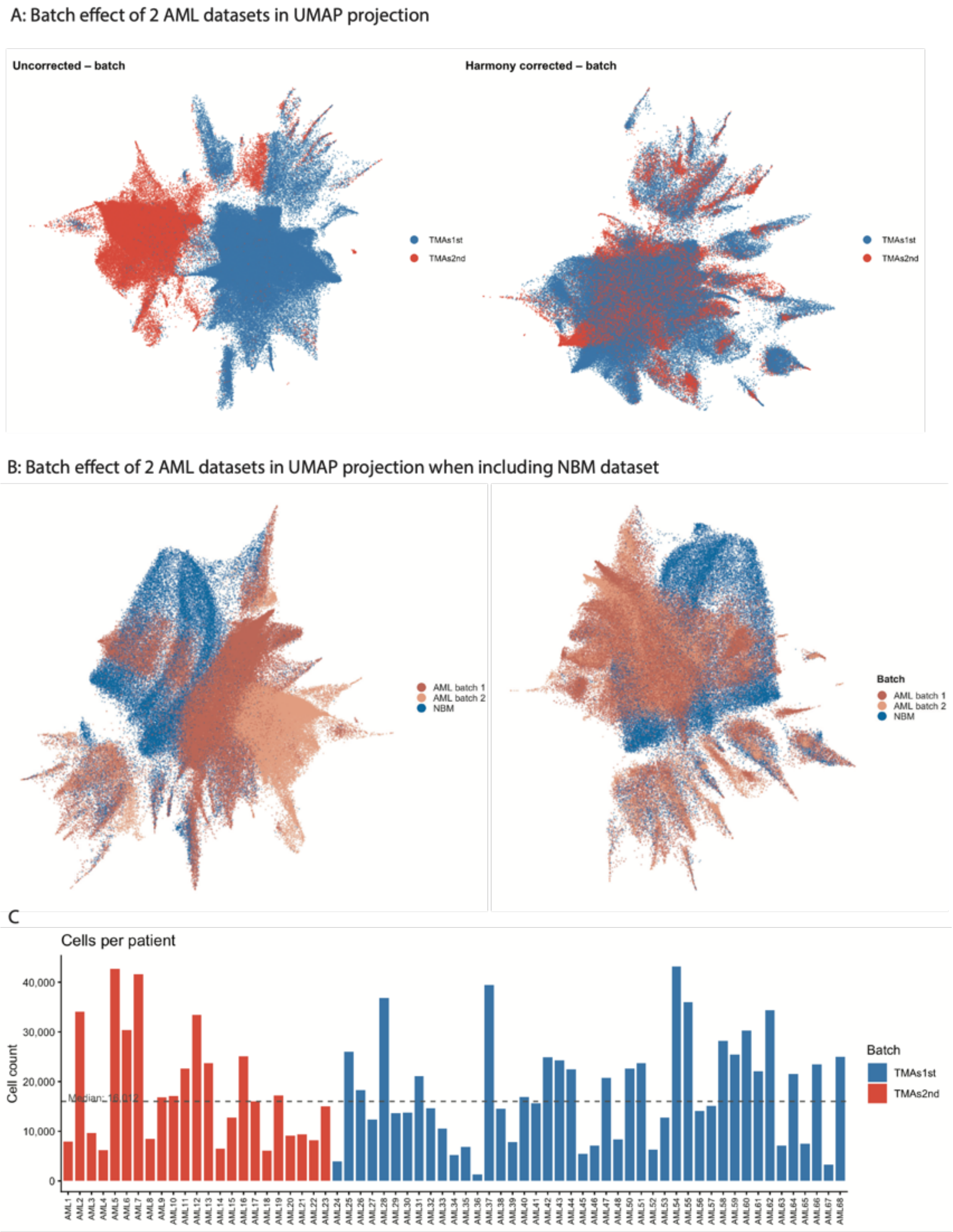
A: Batch effect of 2 AML datasets in UMAP projection
Uncorrected – batch
Harmony corrected – batch
TMAs1st
TMAs2nd
B: Batch effect of 2 AML datasets in UMAP projection when including NBM dataset
AML batch 1
AML batch 2
NBM
Batch
C
Cells per patient
Cell count
40,000
30,000
20,000
10,000
0
Batch
TMAs1st
TMAs2nd

**Figure S6: Imaging batches for AML dataset.**

The AML dataset was imaged in two batches (TMA1st: NPM1-mutant AML TMAs 2 & 3; TMA2nd: NPM1-mutant AML TMA 1). A: Application of batch-correction to AML cell projection in marker space. B: Diminished overall batch effect on UMAP projection if AML and normal bone marrow (NBM) datasets are projected together. C: Cell count per patient across two AML imaging batches.

**Figure S7**

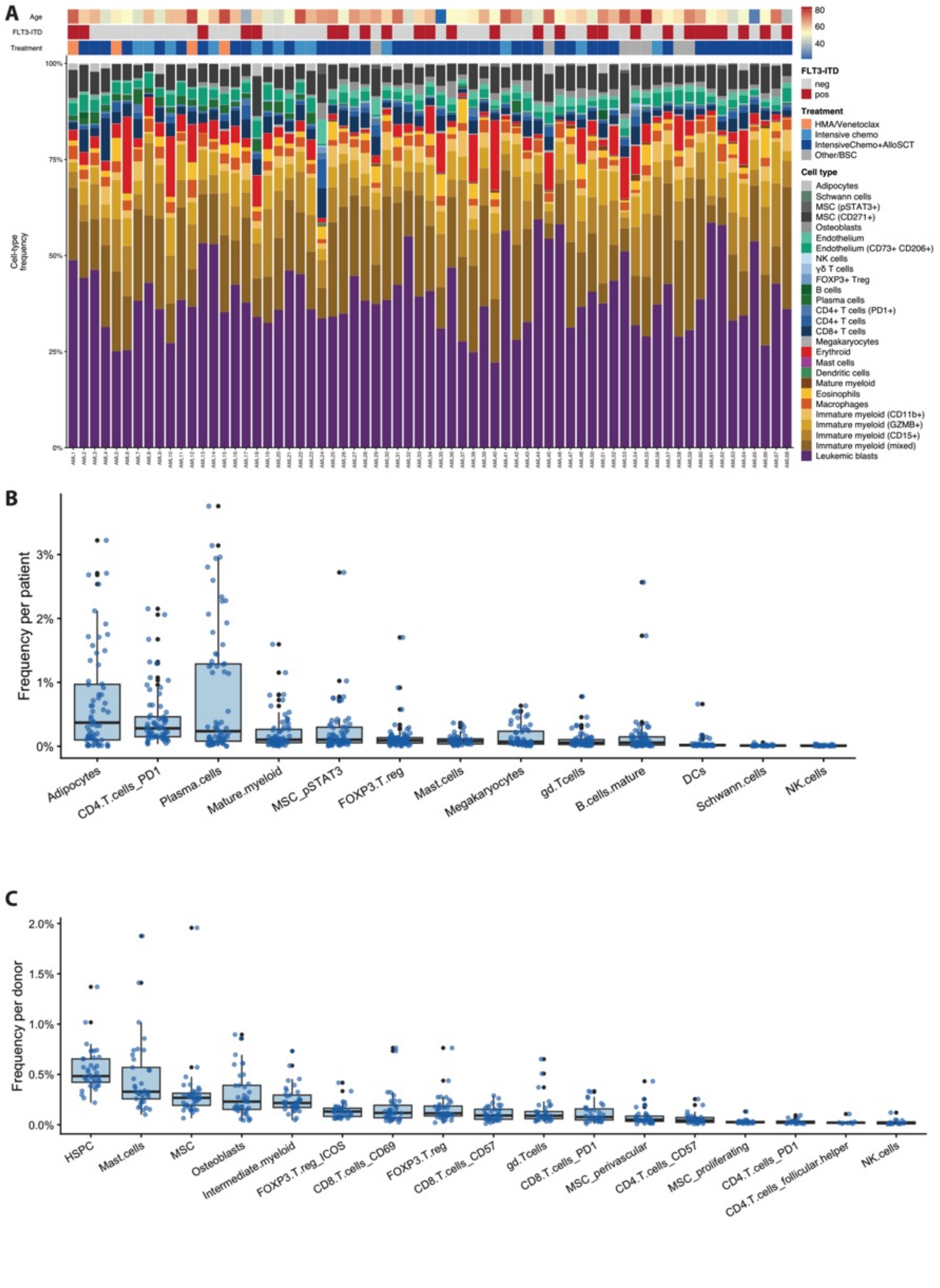
A
Age
FLT3-ITD
Treatment
Cell-type frequency
FLT3-ITD
neg
pos
Treatment
HMA/Venetoclax
Intensive chemo
IntensiveChemo+AlloSCT
Other/BSC
Cell type
Adipocytes
Schwann cells
MSC (pSTAT3+)
MSC (CD271+)
Osteoblasts
Endothelium
Endothelium (CD73+ CD206+)
NK cells
γδ T cells
FOXP3+ Treg
B cells
Plasma cells
CD4+ T cells (PD1+)
CD4+ T cells
CD8+ T cells
Megakaryocytes
Erythroid
Mast cells
Dendritic cells
Mature myeloid
Eosinophils
Macrophages
Immature myeloid (CD11b+)
Immature myeloid (GZMB+)
Immature myeloid (CD15+)
Immature myeloid (mixed)
Leukemic blasts
B
Frequency per patient
Adipocytes
CD4.T.cells_PD1
Plasma.cells
Mature.myeloid
MSC_pSTAT3
FOXP3.T.reg
Mast.cells
Megakaryocytes
gd.Tcells
B.cells.mature
DCs
Schwann.cells
NK.cells
C
Frequency per donor
HSPC
Mast.cells
MSC
Osteoblasts
Intermediate.myeloid
FOXP3.T.reg_ICOS
CD8.T.cells_CD69
FOXP3.T.reg
CD8.T.cells_CD57
gd.Tcells
CD8.T.cells_PD1
MSC_perivascular
CD4.T.cells_CD57
MSC_proliferating
CD4.T.cells_PD1
CD4.T.cells_follicular.helper
NK.cells

**Figure S7: Cell type frequency and detection of rare cell types.**

A: Stacked bar chart depicting cell type frequencies for each *NPM1*-mutant AML patient along with age at diagnosis, *FLT3*-ITD status, and primary treatment. B: Frequency of rare cell types per AML patient. C: Frequency of rare cell types per individual in the normal bone marrow (NBM) cohort.

#### Figure S8

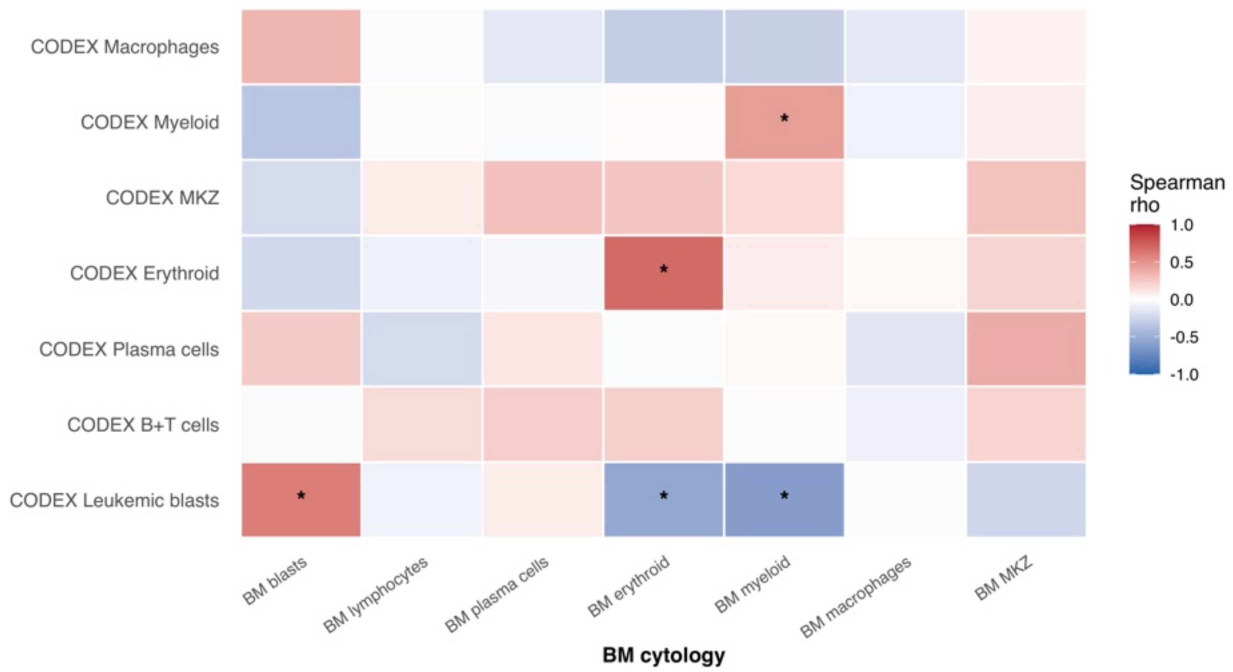


**Figure S8: Correlation in frequencies of CODEX cell types and clinical cytology from simultaneously obtained BM aspirates.**

Heatmap showing correlation between per-patient CODEX cell type fraction and corresponding percentages of cell types from clinical cytology results of diagnostic bone marrow aspirates. Cell types were grouped to harmonize between differences in CODEX annotation and cytology cell type nomenclature. Megakaryocytes (MKZ) were reported on an ordinal scale in clinical cytology results. Spearman rank correlations were computed for each CODEX × cytology pair. P-values were adjusted with the Benjamini-Hochberg method. Only patients with available cytology data (n=43) contributed to each correlation.

**Figure S9**

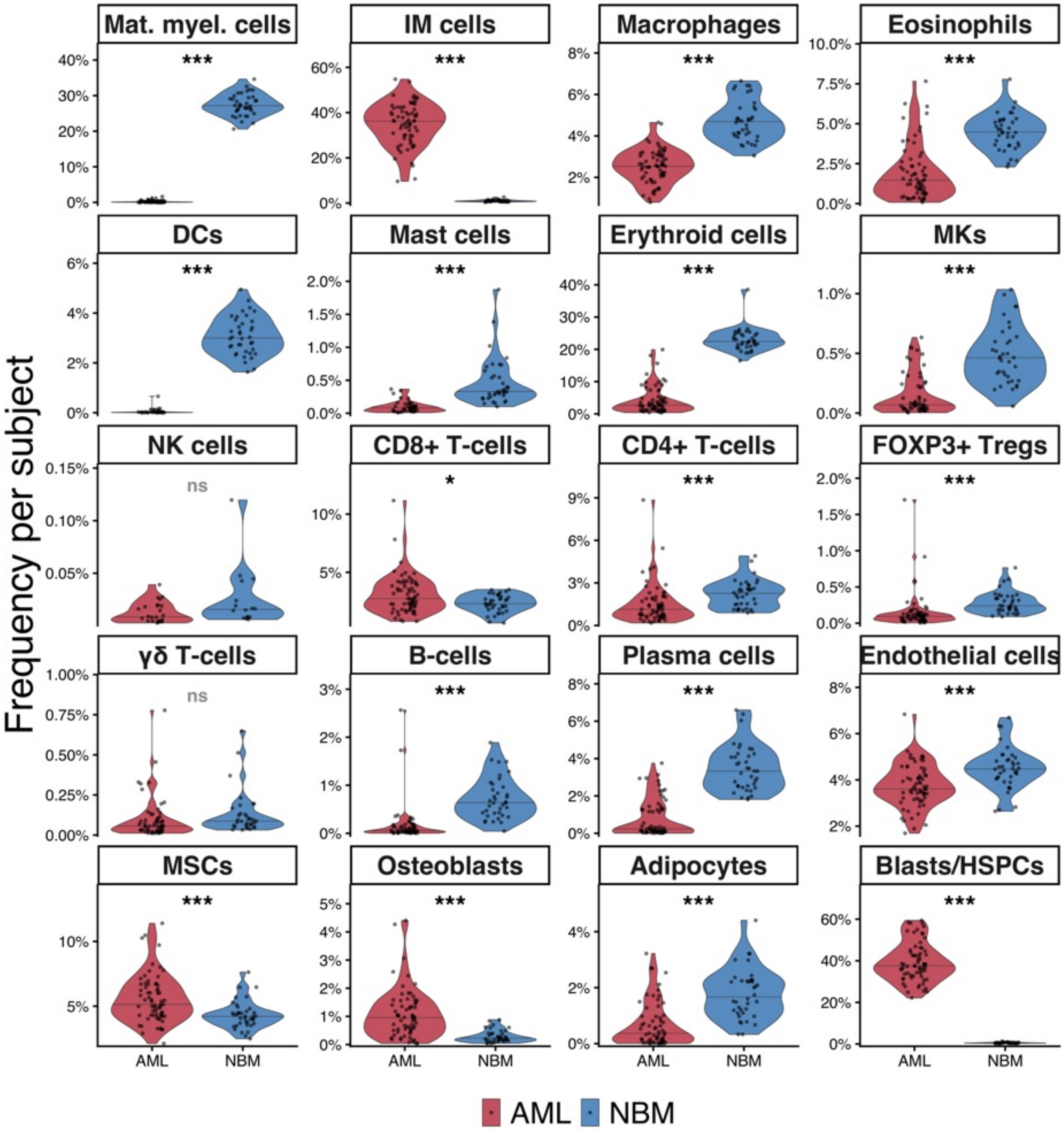
Frequency per subject
Mat. myel. cells
***
IM cells
***
Macrophages
***
Eosinophils
***
DCs
***
Mast cells
***
Erythroid cells
***
MKs
***
NK cells
ns
CD8+ T-cells
*
CD4+ T-cells
***
FOXP3+ Tregs
***
γδ T-cells
ns
B-cells
***
Plasma cells
***
Endothelial cells
***
MSCs
***
Osteoblasts
***
Adipocytes
***
Blasts/HSPCs
***
AML
NBM
AML NBM

**Figure S9: AML vs normal bone marrow cell type frequency.**

Per-subject cell type frequencies were computed for cell types present in both AML and normal bone marrow (NBM) datasets and visualized as violin plots. Significance of frequency differences was assessed by two-sided Wilcoxon rank-sum test with Benjamini–Hochberg FDR correction. Annotation symbols: *** FDR < 0.001, ** < 0.01, * < 0.05, ns = not significant.

#### Figure S10

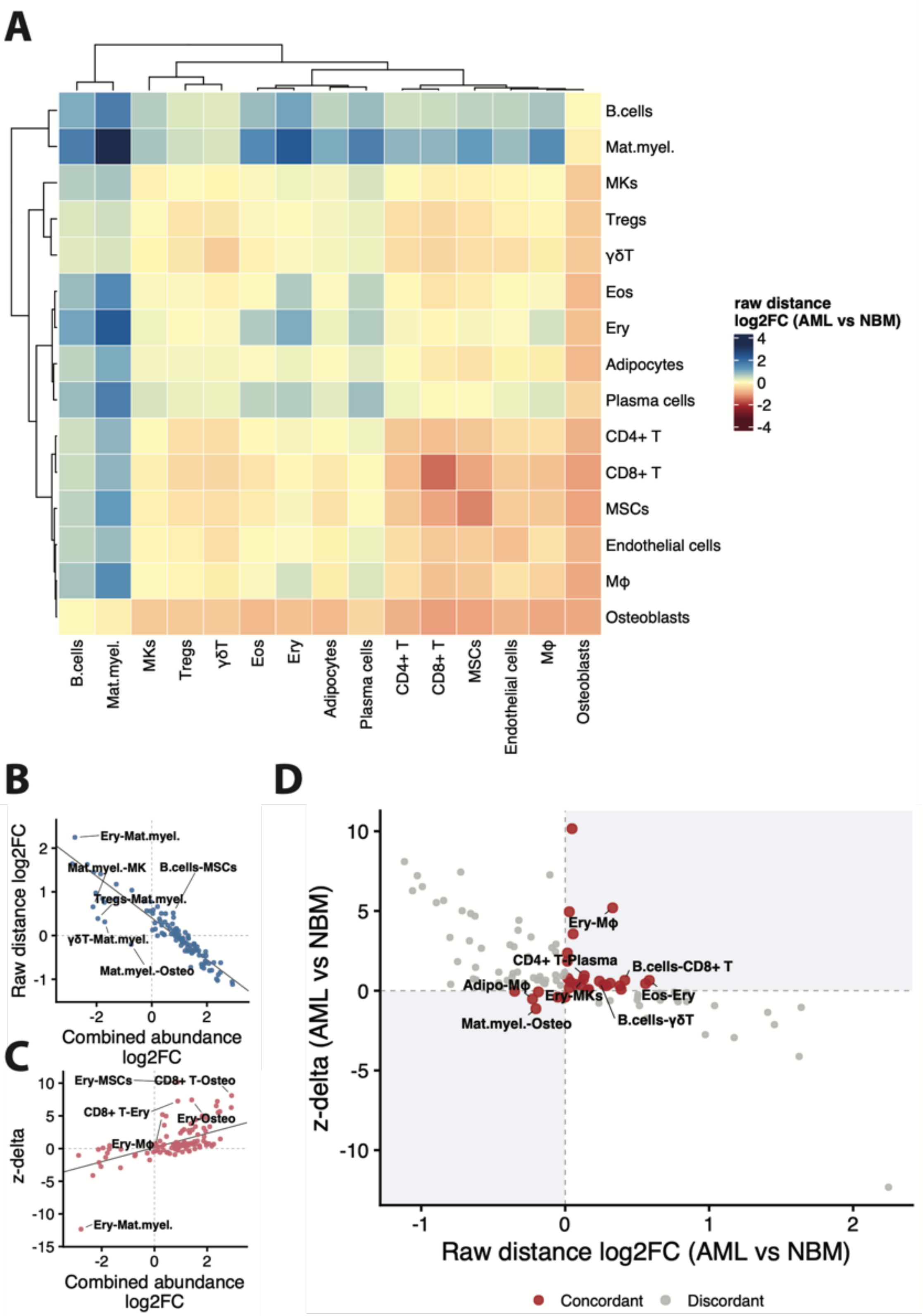
A
B.cells
Mat.myel.
MKs
Tregs
γδT
Eos
Ery
Adipocytes
Plasma cells
CD4+ T
CD8+ T
MSCs
Endothelial cells
Mφ
Osteoblasts
raw distance
log2FC (AML vs NBM)
B
Raw distance log2FC
Combined abundance log2FC
Ery-Mat.myel.
Mat.myel.-MK
B.cells-MSCs
Tregs-Mat.myel.
γδT-Mat.myel.
Mat.myel.-Osteo
C
z-delta
Ery-MSCs
CD8+ T-Osteo
CD8+ T-Ery
Ery-Osteo
Ery-Mφ
Ery-Mat.myel.
D
z-delta (AML vs NBM)
Raw distance log2FC (AML vs NBM)
Ery-Mφ
CD4+ T-Plasma
B.cells-CD8+ T
Adipo-Mφ
Ery-MKs
Eos-Ery
Mat.myel.-Osteo
B.cells-γδT
Concordant
Discordant

**Figure S10: Density-correction of shifts in pairwise cell type distance between the normal bone marrow and AML datasets.**

A: Heatmap visualizing $log_2$ fold-change ($log_2FC$) in raw mean-minimum nearest-neighbor distance between AML and normal bone marrow (NBM) for each cell type (CT) pair.

B: Scatter plot of raw distance $log_2FC$ versus combined abundance $log_2FC$ for each CT pair ($r=-0.91$, $R^2=0.83$); points furthest from the fitted trend are labeled. Combined abundance represents the log2 of the mean fraction of each CT pair.

C: Density-corrected proximity was assessed using a region-conditional permutation test and quantified as a z-score (Supplemental Methods). To compare shifts in pairwise density-corrected proximity between NBM and AML, the difference between z-scores (z-delta = z_AML − z_NBM) was computed. Negative z-delta indicates the CT pair is more attracted relative to chance in AML than in NBM. Scatter plot of z-delta versus combined abundance $log_2FC$ for each CT pair, confounded in the opposite direction to panel B ($r=0.61$, $R^2=0.37$).

D: Scatter plot of z-delta versus raw distance $log_2FC$. Shaded quadrants indicate concordant change in both metrics; 30 of 105 cell-type pairs (red) show concordant change, with the 8 largest-effect pairs labeled.

#### Figure S11

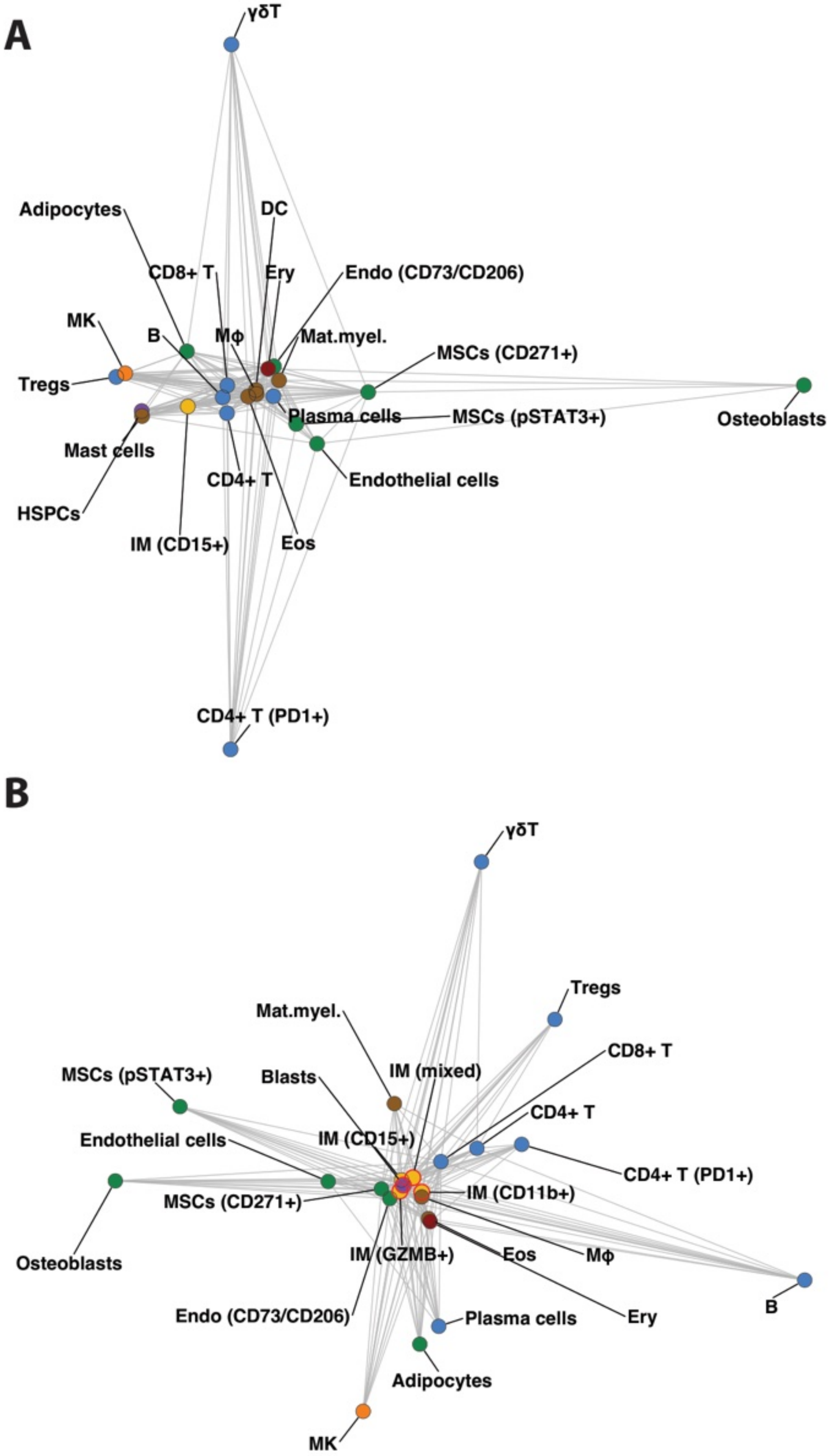
A
γδT
Adipocytes
DC
CD8+ T
Ery
Endo (CD73/CD206)
MK
B
Mϕ
Mat.myel.
MSCs (CD271+)
Tregs
Plasma cells
MSCs (pSTAT3+)
Osteoblasts
Mast cells
CD4+ T
Endothelial cells
HSPCs
IM (CD15+)
Eos
CD4+ T (PD1+)
B
γδT
Tregs
Mat.myel.
IM (mixed)
CD8+ T
MSCs (pSTAT3+)
Blasts
CD4+ T
Endothelial cells
IM (CD15+)
CD4+ T (PD1+)
IM (CD11b+)
MSCs (CD271+)
Osteoblasts
IM (GZMB+)
Eos
Mϕ
Endo (CD73/CD206)
Ery
B
Plasma cells
Adipocytes
MK

**Figure S11: Multidimensional-scaling proximity map of normal bone marrow and AML cell types**

Multidimensional-scaling proximity map of all cell types (CTs) present in the normal bone marrow (NBM) dataset (A), and all CTs in the AML dataset (B), embedded from pairwise raw mean-minimum nearest-neighbor distance (non-metric MDS, isoMDS, k=2). Grey edges connect each CT to up to its 4 nearest spatial neighbors (or any additional pair within 25% of that distance). Stress: NBM 35.6%, AML 27.9%. Panels share a common physical scale, so relative spread/distance is comparable between NBM and AML. Absolute position and orientation of CT nodes are not comparable, as each cohort was embedded independently.

#### Figure S12

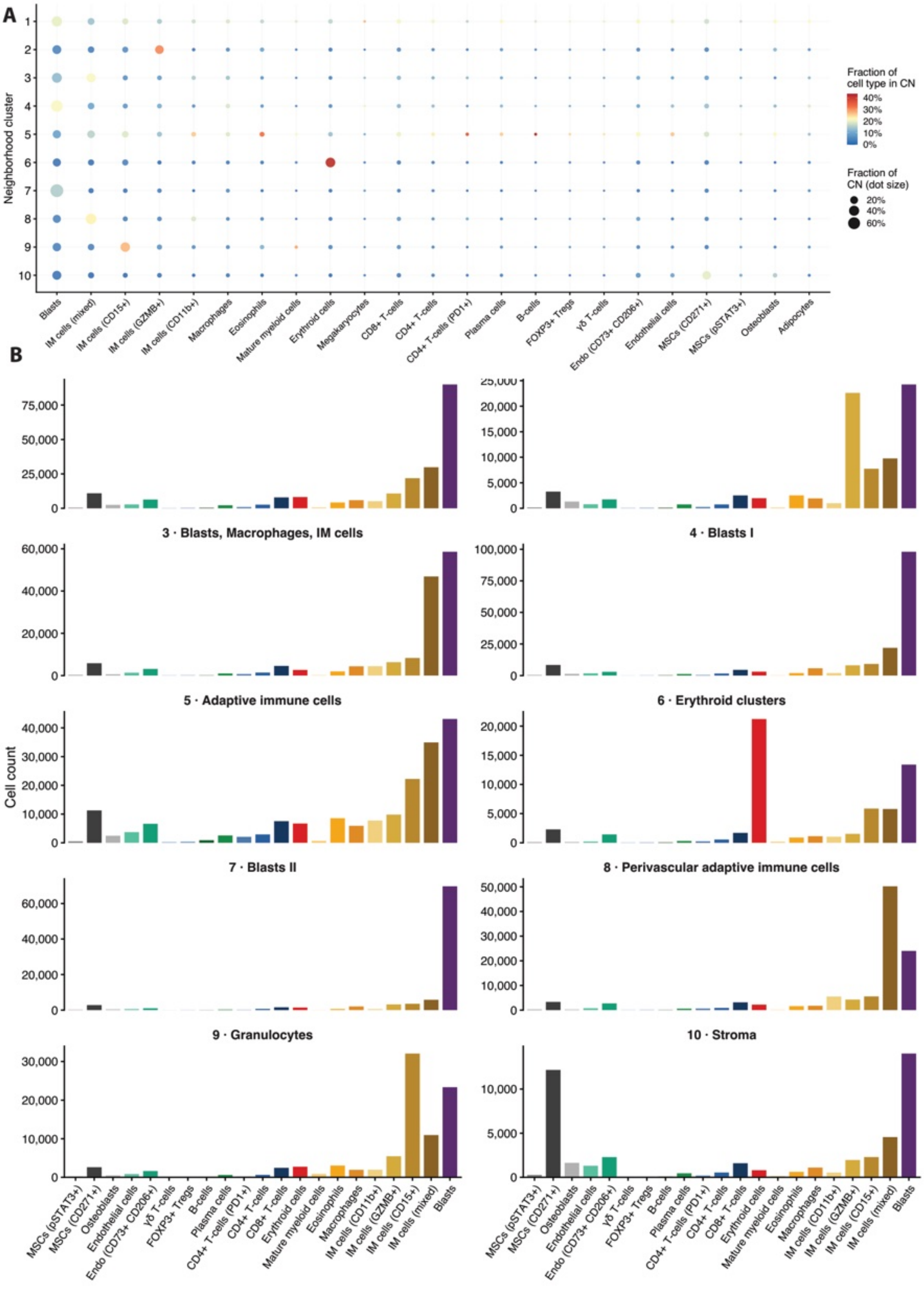
A
B
Neighborhood cluster
Fraction of cell type in CN
Fraction of CN (dot size)
Blasts
IM cells (mixed)
IM cells (CD15+)
IM cells (GZMB+)
IM cells (CD11b+)
Macrophages
Eosinophils
Mature myeloid cells
Erythroid cells
Megakaryocytes
CD8+ T-cells
CD4+ T-cells
CD4+ T-cells (PD1+)
Plasma cells
B-cells
FOXP3+ Tregs
γδ T-cells
Endo (CD73+ CD206+)
Endothelial cells
MSCs (CD271+)
MSCs (pSTAT3+)
Osteoblasts
Adipocytes
3 · Blasts, Macrophages, IM cells
4 · Blasts I
5 · Adaptive immune cells
6 · Erythroid clusters
7 · Blasts II
8 · Perivascular adaptive immune cells
9 · Granulocytes
10 · Stroma
Cell count

**Figure S12: AML neighborhood cluster cell type fractions.** A: Dot plot showing cell type (CT) fraction of all cells in a given cellular neighborhood (CN) (dot size); and fraction of cells from a given CT found in a given CN (dot color). B: Bar charts showing CT counts per CN.

**Figure S13**

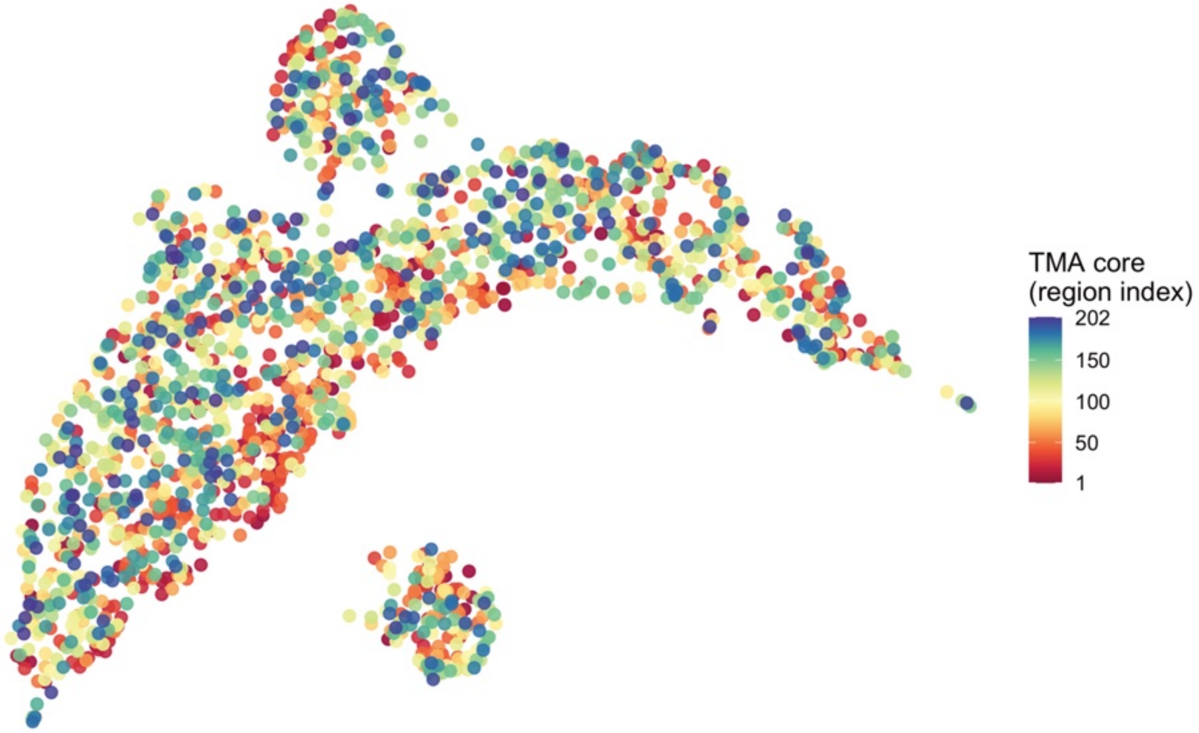


**Figure S13: Neighborhood unit UMAP with superimposed TMA regions.**

UMAP showing distribution of cellular neighborhood (CN) units projected on cell type (CT) composition space. CN units were defined as the entirety of cells belonging to one CN cluster in one TMA core. Dots represent single CN units and are color-labeled by the TMA core corresponding to this CN unit.

#### Figure S14

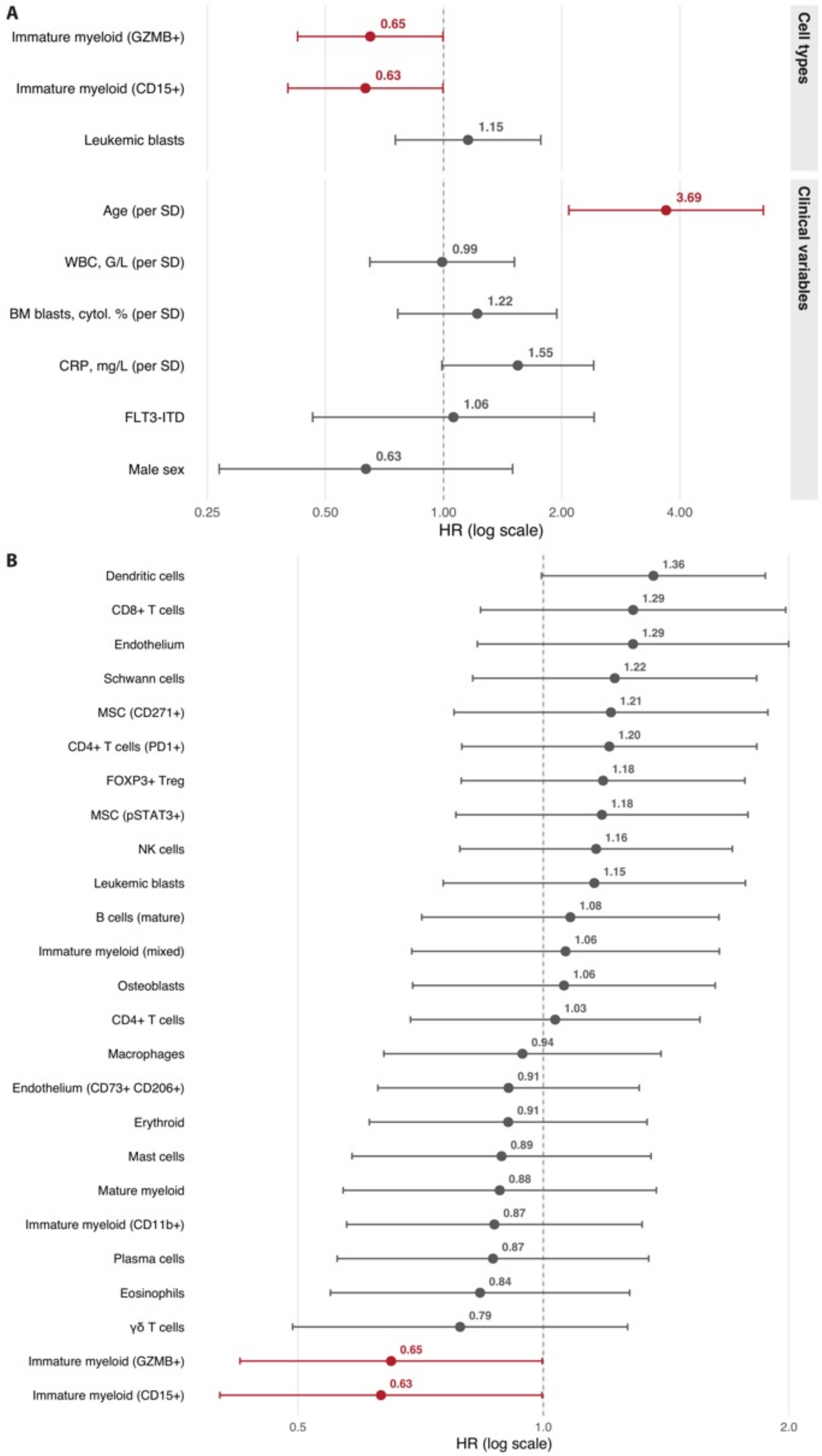
A
Immature myeloid (GZMB+)
0.65
Immature myeloid (CD15+)
0.63
Leukemic blasts
1.15
Age (per SD)
3.69
WBC, G/L (per SD)
0.99
BM blasts, cytol. % (per SD)
1.22
CRP, mg/L (per SD)
1.55
FLT3-ITD
1.06
Male sex
0.63
Cell types
Clinical variables
0.25
0.50
1.00
2.00
4.00
HR (log scale)
B
Dendritic cells 1.36
CD8+ T cells 1.29
Endothelium 1.29
Schwann cells 1.22
MSC (CD271+) 1.21
CD4+ T cells (PD1+) 1.20
FOXP3+ Treg 1.18
MSC (pSTAT3+) 1.18
NK cells 1.16
Leukemic blasts 1.15
B cells (mature) 1.08
Immature myeloid (mixed) 1.06
Osteoblasts 1.06
CD4+ T cells 1.03
Macrophages 0.94
Endothelium (CD73+ CD206+) 0.91
Erythroid 0.91
Mast cells 0.89
Mature myeloid 0.88
Immature myeloid (CD11b+) 0.87
Plasma cells 0.87
Eosinophils 0.84
γδ T cells 0.79
Immature myeloid (GZMB+) 0.65
Immature myeloid (CD15+) 0.63
0.5
1.0
2.0
HR (log scale)

**Figure S14: Univariate association of baseline clinical variables and cell types with survival.**

Forest plots displaying univariate hazard ratio (HR) and 95% confidence interval for association of clinical variable and cell type (CT) frequency with overall survival (OS). A: Selected CTs and clinical variables. B: All CTs.

#### Figure S15

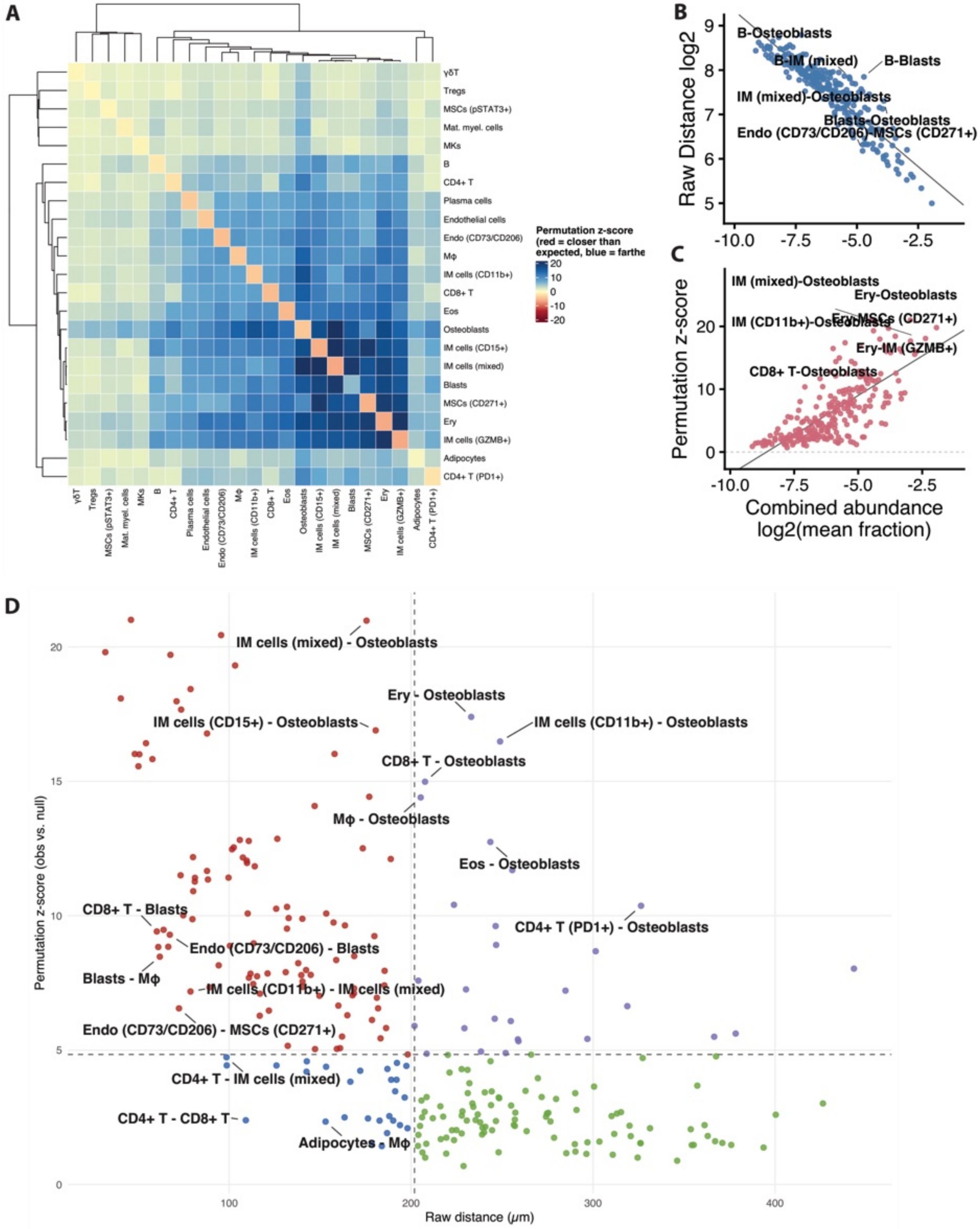
A
Permutation z-score (red = closer than expected, blue = farthe
γδT
Tregs
MSCs (pSTAT3+)
Mat. myel. cells
MKs
B
CD4+ T
Plasma cells
Endothelial cells
Endo (CD73/CD206)
Mφ
IM cells (CD11b+)
CD8+ T
Eos
Osteoblasts
IM cells (CD15+)
IM cells (mixed)
Blasts
MSCs (CD271+)
Ery
IM cells (GZMB+)
Adipocytes
CD4+ T (PD1+)
B
Raw Distance log2
B-Osteoblasts
B-IM (mixed)
B-Blasts
IM (mixed)-Osteoblasts
Blasts-Osteoblasts
Endo (CD73/CD206)-MSCs (CD271+)
C
Permutation z-score
IM (mixed)-Osteoblasts
Ery-Osteoblasts
IM (CD11b+)-Osteoblasts
Ery-MSCs (CD271+)
Ery-IM (GZMB+)
CD8+ T-Osteoblasts
Combined abundance log2(mean fraction)
D
Permutation z-score (obs vs. null)
Raw distance (μm)
IM cells (mixed) - Osteoblasts
Ery - Osteoblasts
IM cells (CD15+) - Osteoblasts
IM cells (CD11b+) - Osteoblasts
CD8+ T - Osteoblasts
Mφ - Osteoblasts
Eos - Osteoblasts
CD8+ T - Blasts
CD4+ T (PD1+) - Osteoblasts
Endo (CD73/CD206) - Blasts
Blasts - Mφ
IM cells (CD11b+) - IM cells (mixed)
Endo (CD73/CD206) - MSCs (CD271+)
CD4+ T - IM cells (mixed)
CD4+ T - CD8+ T
Adipocytes - Mφ

**Figure S15: Density-corrected proximity of AML cell types.**

Raw mean-minimum nearest-neighbor distance and density-corrected proximity, assessed using a region-conditional permutation test and quantified as a z-score, were calculated for all AML cell type (CT) pairs (Supplemental Methods).

A: Permutation z-score heatmap for AML cell type pairs, including self-association. Red = closer than expected under the density-matched null; blue = farther than expected.

B: Scatter plot of $\log_2$(raw distance) versus combined abundance ($\log_2$ of mean fraction of CT pair) for each CT pair (r=-0.91, $R^2$=0.83); labeled points are the pairs farthest from the fitted trend.

C: Scatter plot of permutation z-score versus combined abundance for each CT pair (r=0.74, $R^2$=0.55), confounded in the opposite direction to panel B.

D: Raw distance versus permutation z-score for each CT pair, classified into four quadrants by median split on each. Labeled pairs are those with concordant deviation from both metrics' abundance-predicted values.

#### Figure S16

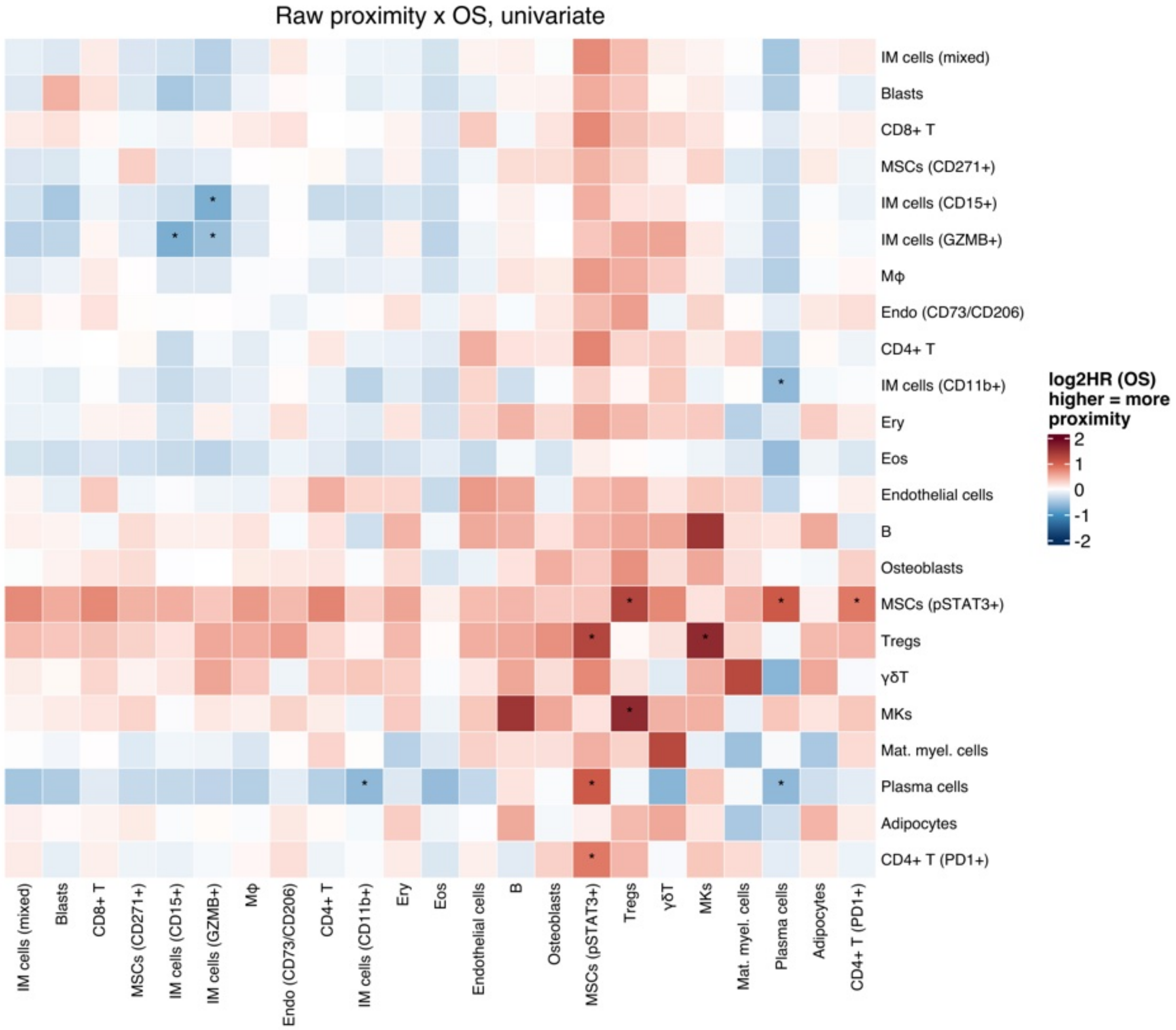
Raw proximity x OS, univariate
IM cells (mixed)
Blasts
CD8+ T
MSCs (CD271+)
IM cells (CD15+)
IM cells (GZMB+)
Mϕ
Endo (CD73/CD206)
CD4+ T
IM cells (CD11b+)
Ery
Eos
Endothelial cells
B
Osteoblasts
MSCs (pSTAT3+)
Tregs
γδT
MKs
Mat. myel. cells
Plasma cells
Adipocytes
CD4+ T (PD1+)
log2HR (OS) higher = more proximity
2
1
0
-1
-2

**Figure S16: Association of pairwise raw distance with overall survival.**

Heatmap of $\log_2$(hazard ratio) for overall survival (OS) from univariate Cox proportional-hazards models, one per CT pair, using mean nearest-neighbor distance between the two cell types as a continuous, z-scored predictor. Asterisk (*) denotes $p < 0.05$. Cell type pairs with fewer than 15 patients contributing non-missing distance data were excluded.

#### Figure S17

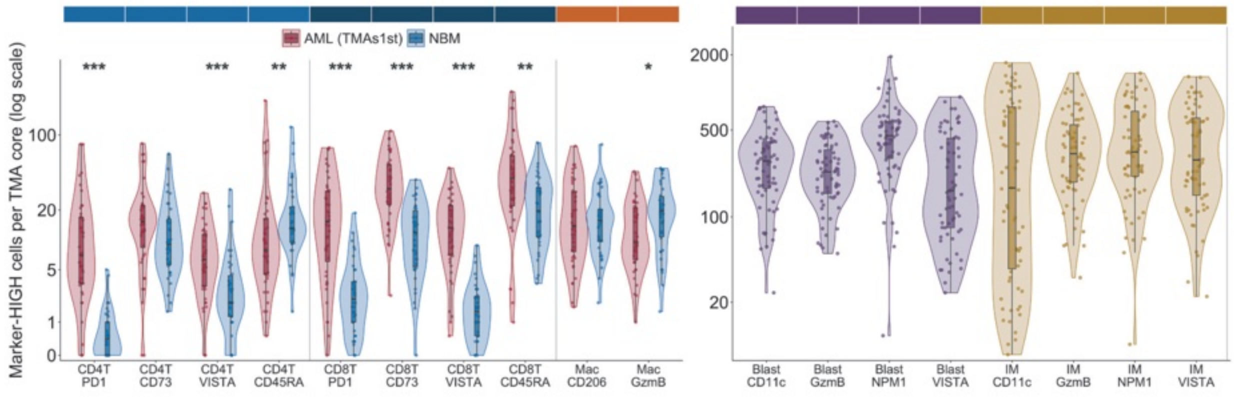


**Figure S17: Marker expression across patients in immune cells, immature myeloid cells, and leukemic blasts.**

Marker expression analysis, restricted to one imaging batch containing 2 AML TMAs and the NBM batch to minimize marker intensity batch effects. Marker-high fractions were gated per batch using gaussian mixed models.

#### Figure S18

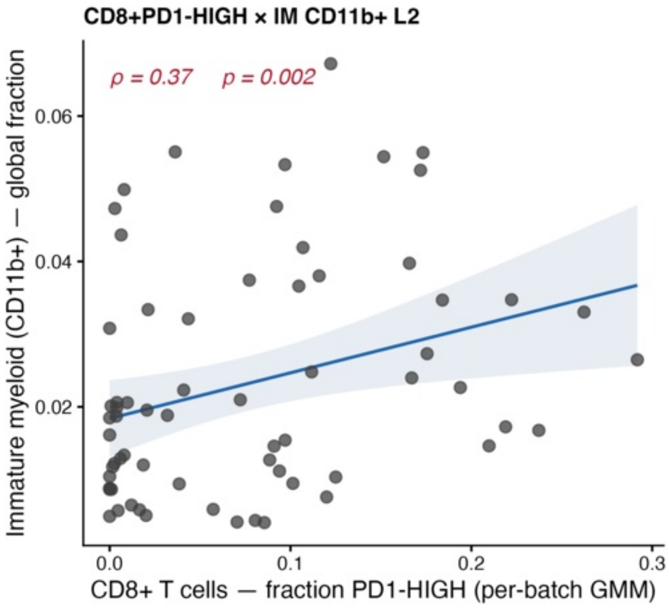


**Figure S18: Correlation between frequency of PD1-high CD8+ T-cells and CD11b+ immature myeloid cells in AML bone marrow per patient.**

Correlation of the fraction of CD11b+ immature myeloid cells and PD1-high CD8+ T-cells per patient using spearman rho. PD1-high CD8+ T-cell populations were gated per batch using gaussian mixed models.

#### Figure S19

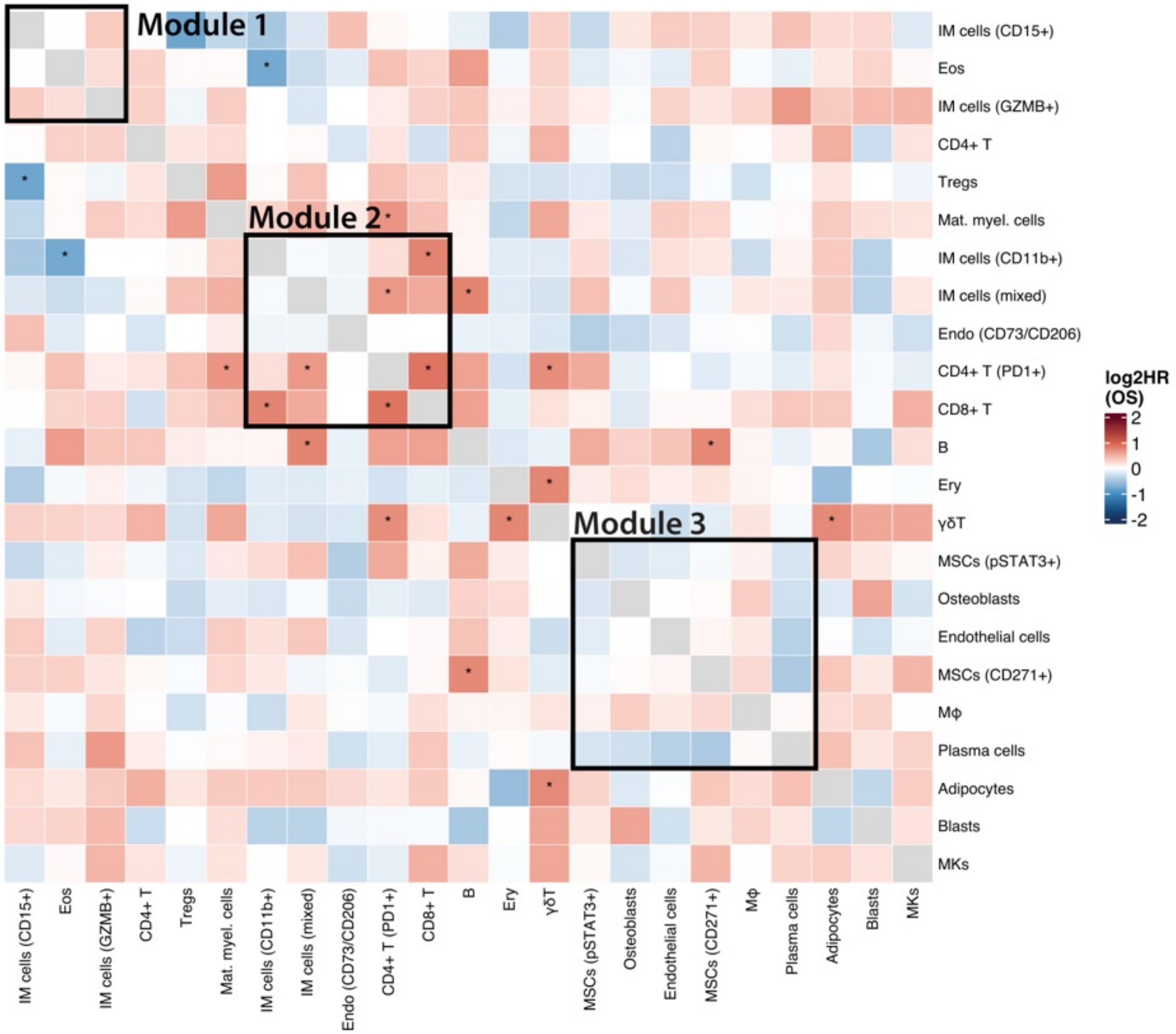
Module 1
Module 2
Module 3
IM cells (CD15+)
Eos
IM cells (GZMB+)
CD4+ T
Tregs
Mat. myel. cells
IM cells (CD11b+)
IM cells (mixed)
Endo (CD73/CD206)
CD4+ T (PD1+)
CD8+ T
B
Ery
γδT
MSCs (pSTAT3+)
Osteoblasts
Endothelial cells
MSCs (CD271+)
Mϕ
Plasma cells
Adipocytes
Blasts
MKs
log2HR (OS)
2
1
0
-1
-2

**Figure S19: Association of pairwise co-enrichment with overall survival.**

Heatmap of $\log_2$(hazard ratio) for overall survival from univariate Cox proportional-hazards models, one per cell type (CT) pair, using the per-patient Spearman correlation ($\rho$) of the two CTs' raw fractions across shared cellular neighborhoods ($\geq$4 neighborhoods required per patient) as a continuous, z-scored predictor. Asterisk (*) denotes $p < 0.05$. Black boxes indicate modules of cell types with mutual co-enrichment (Modules 1–3). CT pairs with fewer than 15 patients contributing non-missing co-enrichment data were excluded.

#### Figure S20

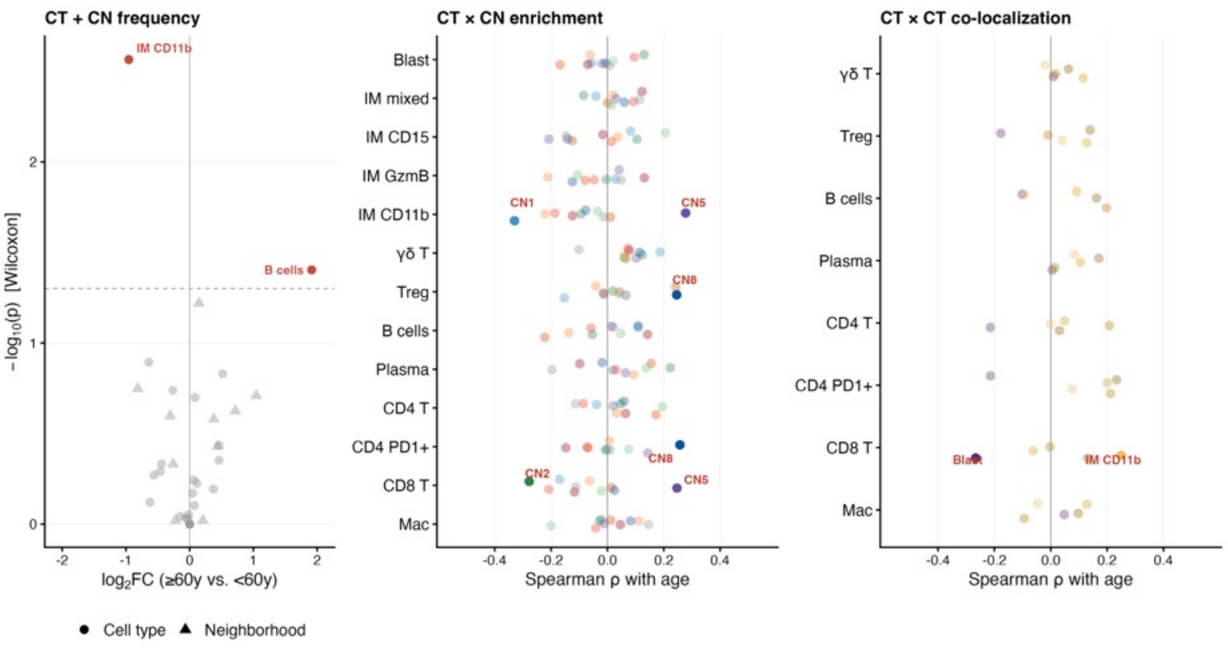


**Figure S20: Correlation of cell type frequencies, neighborhood frequencies, cell type in neighborhood enrichment, and pairwise cell type co-enrichment with age.**

Left, volcano plot showing enrichment of cell types (CTs) and cellular neighborhoods (CNs) in patients younger vs. older than 60 years. Center, correlation of CT in CN enrichment with age for immune, immature myeloid and leukemic blast CTs. Right, correlation of CT co-enrichment with age for immune CTs (x axis) and immature myeloid and leukemic blast CTs (dots).

**Figure S21**

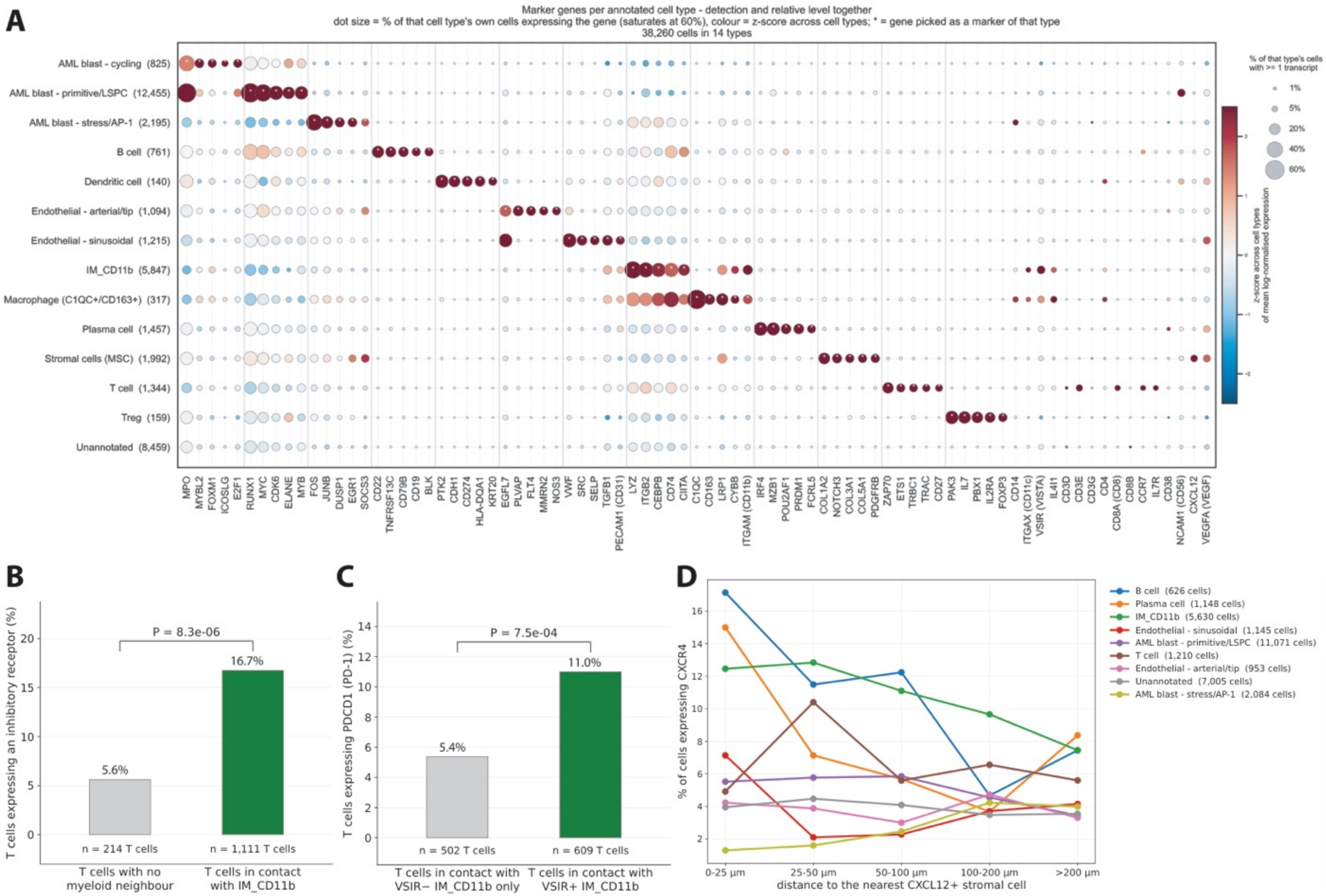
A
Marker genes per annotated cell type - detection and relative level together
dot size = % of that cell type's own cells expressing the gene (saturates at 60%), colour = z-score across cell types; * = gene picked as a marker of that type
38,260 cells in 14 types
AML blast - cycling (825)
AML blast - primitive/LSPC (12,455)
AML blast - stress/AP-1 (2,195)
B cell (761)
Dendritic cell (140)
Endothelial - arterial/tip (1,094)
Endothelial - sinusoidal (1,215)
IM_CD11b (5,847)
Macrophage (C1QC+/CD163+) (317)
Plasma cell (1,457)
Stromal cells (MSC) (1,992)
T cell (1,344)
Treg (159)
Unannotated (8,459)
MPO MYBL2 FOXM1 ICOSLG E2F1 RUNX1 MYC CDK6 ELANE MYB FOS JUNB DUSP1 EGR1 SOCS3 CD22 TNFRSF13C CD79B CD19 BLK PTK2 CDH1 CD274 HLA-DQA1 KRT20 EGFL7 PLVAP FLT4 MMRN2 NOS3 VWF SRC SELP TGFB1 PECAM1 (CD31) LYZ ITGB2 CEBPB CD74 CIITA C1QC CD163 LRP1 CYBB ITGAM (CD11b) IRF4 MZB1 POU2AF1 PRDM1 FCRL5 COL1A2 NOTCH3 COL3A1 COL5A1 PDGFRB ZAP70 ETS1 TRBC1 TRAC CD27 PAX3 IL7 PBX1 IL2RA FOXP3 CD14 ITGAX (CD11c) VSIR (VISTA) IL4I1 CD3D CD3E CD3G CD4 CD8A (CD8) CD8B CCR7 IL7R CD38 NCAM1 (CD56) CXCL12 VEGFA (VEGF)
% of that type's cells with >= 1 transcript
1%
5%
20%
40%
60%
z-score across cell types of mean log-normalised expression
B
P = 8.3e-06
T cells expressing an inhibitory receptor (%)
5.6%
16.7%
n = 214 T cells
n = 1,111 T cells
T cells with no myeloid neighbour
T cells in contact with IM_CD11b
C
P = 7.5e-04
T cells expressing PDCD1 (PD-1) (%)
5.4%
11.0%
n = 502 T cells
n = 609 T cells
T cells in contact with VSIR− IM_CD11b only
T cells in contact with VSIR+ IM_CD11b
D
% of cells expressing CXCR4
0-25 µm
25-50 µm
50-100 µm
100-200 µm
>200 µm
distance to the nearest CXCL12+ stromal cell
B cell (626 cells)
Plasma cell (1,148 cells)
IM_CD11b (5,630 cells)
Endothelial - sinusoidal (1,145 cells)
AML blast - primitive/LSPC (11,071 cells)
T cell (1,210 cells)
Endothelial - arterial/tip (953 cells)
Unannotated (7,005 cells)
AML blast - stress/AP-1 (2,084 cells)

**Figure S21: Spatial transcriptome cell type annotation.**

A: MERFISH spatial transcriptomic profiling was performed using the 815-gene Immuno-Oncology panel on 22 bone-marrow trephine biopsies from the *NPM1*-mutant AML cohort. A total of 306,445 cells were initially detected, of which 38,260 cells passed the predefined quality-control criteria and were included in downstream analyses. Four of 22 biopsies failed the quality-control criteria, 18 of 22 contributed at least one cell, and 12 of 22 provided enough cells for spatial statistical analyses. Cell types (CTs) with at least 50 cells were ranked based on Wilcoxon test against all other types. The top 5 genes were retained per CT, resulting in overall 65 genes. 16 genes were selected as the corresponding canonical CODEX markers (CD14, ITGAX, VSIR, IL4I1, CD3D, CD3E, CD3G, CD4, CD8A, CD8B, CCR7, IL7R, CD38, NCAM1, CXCL12, VEGFA). These markers are also shown in the heatmap (regardless of their rank) because they were used in the CT annotation step. Color is the z-score across CTs, i.e. representing not absolute expression but relative enrichment.

B: T-cells in interface contact with IM_CD11b cells express an inhibitory receptor (any of *PDCD1*, *CTLA4*, *HAVCR2*, *LAG3*, *TIGIT*, *ENTPD1*, *CD244*) more frequently than T-cells with no myeloid neighbor (16.7% of 1,111 vs 5.6% of 214).

C: Among T-cells in interface contact with IM_CD11b cells, those with at least one *VSIR*+ IM_CD11b neighbor express *PDCD1* more frequently (11.0% of 609, i.e. 67 cells, 5.1% of all 1,325 T-cells) than those whose IM_CD11b neighbors are all *VSIR*− (5.4% of 502).

D: For each distance range from the nearest *CXCL12*+ stromal cell (0-25, 25-50, 50-100, 100-200, over 200 µm), the proportion of cells expressing *CXCR4* was plotted. The same measurement was repeated against *CXCL12* negative stromal cells to distinguish a ligand specific proximity to MSCs from a general proximity.

#### SUPPLEMENTAL TABLES

#### Table S1: NPM1-mutant AML cohort.

| UPN | FLT3 ITD | Sex | Age | Age at diagnosis | % Blasts cytology | % Blasts histology | Year of first diagnosis | 1st line | alloHSCT | Response | Relapse | Death | Status |
|---|---|---|---|---|---|---|---|---|---|---|---|---|---|
| AML01 | yes | male | ≥ 60 | 75 | 96 | NA | 2022 | HMA/Ven | no | no CR post induction | no | yes | Relapse mortality |
| AML02 | yes | male | ≥ 60 | 60 | 98 | NA | 2022 | Intensive induction | yes | no CR post induction | yes | yes | Relapse mortality |
| AML03 | no | female | ≥ 60 | 63 | 66.5 | NA | 2022 | Intensive induction | yes | no CR post induction | no | no | Alive w/o relapse |
| AML04 | no | female | < 60 | 47 | 69.5 | NA | 2022 | Intensive induction | yes | no CR post induction | no | no | Alive w/o relapse |
| AML05 | no | male | < 60 | 55 | 15 | 18 | 2021 | HMA/Ven | yes | CR post induction | no | no | Alive w/o relapse |
| AML06 | no | male | < 60 | 53 | NA | NA | 2022 | Intensive induction | yes | CR post induction | no | no | Alive w/o relapse |
| AML07 | no | female | < 60 | 47 | 55 | 95 | 2022 | Intensive induction | no | CR post induction | no | no | Alive w/o relapse |
| AML08 | no | female | ≥ 60 | 62 | 71 | 71 | 2021 | Intensive induction | no | no CR post induction | no | no | Alive w/o relapse |
| AML09 | no | male | < 60 | 48 | 69.5 | 75 | 2021 | Intensive induction | yes | CR post induction | yes | yes | Relapse mortality |
| AML10 | no | female | ≥ 60 | 62 | 32 | 20 | 2021 | Intensive induction | no | no CR post induction | no | no | Alive w/o relapse |
| AML11 | no | female | ≥ 60 | 62 | 90 | 90 | 2021 | Intensive induction | yes | no CR post induction | no | no | Alive w/o relapse |
| AML12 | no | male | ≥ 60 | 79 | 58.5 | NA | 2021 | HMA/Ven | no | no CR post induction | no | no | Alive w/o relapse |
| AML13 | yes | female | < 60 | 54 | 90 | 80 | 2021 | Intensive induction | yes | no CR post induction | yes | no | Alive w/ relapse |
| AML14 | no | male | ≥ 60 | 61 | 90 | NA | 2020 | Intensive induction | no | no CR post induction | no | no | Alive w/o relapse |
| AML15 | no | male | ≥ 60 | 78 | 51.5 | NA | 2019 | HMA/Ven | no | no CR post induction | no | yes | Relapse mortality |
| AML16 | no | female | ≥ 60 | 62 | 89.5 | 90 | 2019 | Intensive induction | yes | CR post induction | no | no | Alive w/o relapse |
| AML17 | yes | female | < 60 | 37 | 72 | NA | 2019 | Intensive induction | yes | no CR post induction | no | no | Alive w/o relapse |
| AML18 | yes | female | < 60 | 59 | 37 | 15 | 2018 | Intensive induction | no | CR post induction | no | no | Alive w/o relapse |
| AML19 | no | male | < 60 | 48 | 41.5 | NA | 2018 | Intensive induction | yes | CR post induction | yes | no | Alive w/ relapse |
| AML20 | no | female | < 60 | 46 | 20.5 | 10 | 2018 | Intensive induction | no | no CR post induction | no | no | Alive w/o relapse |
| AML21 | no | female | < 60 | 51 | 70 | NA | 2018 | Intensive induction | yes | CR post induction | yes | no | Alive w/ relapse |
| AML22 | no | male | ≥ 60 | 77 | 86.5 | 100 | 2017 | Intensive induction | no | no CR post induction | no | yes | Relapse mortality |
| AML23 | no | female | ≥ 60 | 63 | 73 | 50 | 2017 | Intensive induction | no | CR post induction | no | no | Alive w/o relapse |
| AML24 | no | male | ≥ 60 | 67 | 56.5 | 20 | 2012 | Intensive induction | yes | no CR post induction | no | yes | Non-relapse mortality |
| AML25 | yes | male | ≥ 60 | 68 | NA | NA | 2012 | Intensive induction | yes | | yes | yes | Relapse mortality |
| AML26 | yes | male | < 60 | 50 | 28 | NA | 2013 | Intensive induction | yes | CR post induction | no | no | Alive w/o relapse |
| AML27 | no | female | ≥ 60 | 67 | 69 | NA | 2013 | Intensive induction | yes | no CR post induction | yes | yes | Relapse mortality |
| AML28 | yes | male | ≥ 60 | 63 | 83 | NA | 2013 | Intensive induction | yes | no CR post induction | yes | no | Alive w/ relapse |
| AML29 | no | male | ≥ 60 | 66 | NA | 15 | 2013 | HMA | yes | | yes | yes | Relapse mortality |
| AML30 | yes | female | < 60 | 58 | 17 | NA | 2013 | Intensive induction | no | no CR post induction | yes | yes | Relapse mortality |
| AML31 | no | male | ≥ 60 | 73 | 17 | NA | 2013 | Intensive induction | yes | no CR post induction | no | yes | Non-relapse mortality |
| AML32 | no | male | < 60 | 46 | 78.5 | 10 | 2014 | Intensive induction | yes | CR post induction | yes | no | Alive w/ relapse |
| AML33 | yes | male | < 60 | 58 | 94.5 | NA | 2014 | Intensive induction | yes | no CR post induction | no | no | Alive w/o relapse |
| AML34 | yes | male | ≥ 60 | 63 | NA | 50 | 2014 | Intensive induction | yes | no CR post induction | no | yes | Non-relapse mortality |
| AML35 | no | male | < 60 | 22 | 16.5 | 15 | 2014 | Intensive induction | yes | no CR post induction | yes | no | Alive w/ relapse |
| AML36 | yes | male | < 60 | 53 | 23.5 | NA | 2015 | Intensive induction | yes | CR post induction | yes | no | Alive w/ relapse |
| AML37 | no | female | < 60 | 52 | NA | NA | 2015 | Intensive induction | yes | CR post induction | yes | no | Alive w/ relapse |
| AML38 | no | male | < 60 | 55 | 12.5 | NA | 2015 | Intensive induction | yes | CR post induction | no | no | Alive w/o relapse |
| AML39 | no | female | ≥ 60 | 64 | 91 | 80 | 2015 | Intensive induction | yes | CR post induction | yes | no | Alive w/ relapse |
| AML40 | yes | female | < 60 | 52 | 3 | NA | 2015 | Intensive induction | yes | CR post induction | no | no | Alive w/o relapse |
| AML41 | no | female | ≥ 60 | 75 | 77.5 | 90 | 2015 | Intensive induction | no | no CR post induction | no | yes | Non-relapse mortality |
| AML42 | no | male | ≥ 60 | 69 | 5 | 40 | 2016 | Intensive induction | yes | no CR post induction | yes | yes | Relapse mortality |
| AML43 | no | female | < 60 | 46 | 61.5 | NA | 2016 | Intensive induction | yes | CR post induction | yes | yes | Relapse mortality |
| AML44 | yes | male | < 60 | 49 | NA | NA | 2016 | Intensive induction | yes | no CR post induction | yes | yes | Relapse mortality |
| AML45 | no | female | ≥ 60 | 76 | 39 | 80 | 2016 | HMA | no | | no | yes | Relapse mortality |
| AML46 | yes | male | < 60 | 52 | 73.5 | NA | 2016 | Intensive induction | yes | no CR post induction | no | no | Alive w/o relapse |
| AML47 | no | male | < 60 | 57 | 5 | NA | 2017 | Intensive induction | yes | CR post induction | yes | no | Alive w/ relapse |
| AML48 | no | female | ≥ 60 | 60 | 25 | NA | 2017 | Intensive induction | no | CR post induction | yes | no | Alive w/ relapse |
| AML50 | yes | male | < 60 | 49 | 59 | NA | 2017 | Intensive induction | yes | no CR post induction | no | no | Alive w/o relapse |
| AML51 | yes | male | ≥ 60 | 69 | 35 | NA | 2015 | Intensive induction | yes | no CR post induction | no | no | Alive w/o relapse |
| AML52 | no | female | ≥ 60 | 63 | NA | 9 | 2019 | Intensive induction | yes | CR post induction | yes | no | Alive w/ relapse |
| AML53 | no | female | ≥ 60 | 76 | 39 | 80 | 2016 | HMA | no | | no | yes | Relapse mortality |
| AML54 | yes | male | ≥ 60 | 66 | 86 | NA | 2017 | HMA | yes | no CR post induction | no | yes | Non-relapse mortality |
| AML55 | no | female | ≥ 60 | 83 | NA | NA | 2020 | HMA | no | | no | no | Alive w/o relapse |
| AML56 | no | male | < 60 | 59 | 58 | NA | 2018 | Intensive induction | no | | no | yes | Non-relapse mortality |
| AML57 | yes | female | < 60 | 54 | 12 | NA | 2013 | Intensive induction | yes | CR post induction | no | no | Alive w/o relapse |
| AML58 | no | male | < 60 | 57 | NA | NA | 2015 | HMA | yes | | no | no | Alive w/o relapse |
| AML59 | yes | female | ≥ 60 | 70 | 70 | NA | 2017 | HMA | yes | | yes | yes | Relapse mortality |
| AML60 | yes | male | ≥ 60 | 65 | 80 | NA | 2022 | Intensive induction | yes | CR post induction | yes | yes | Relapse mortality |
| AML61 | yes | female | < 60 | 52 | 95 | 85 | 2020 | Intensive induction | yes | CR post induction | no | no | Alive w/o relapse |
| AML62 | yes | male | < 60 | 56 | 89 | NA | 2020 | Intensive induction | yes | | yes | no | Alive w/ relapse |
| AML63 | no | female | < 60 | 46 | 51.5 | NA | 2007 | Intensive induction | yes | | no | no | Alive w/o relapse |
| AML64 | yes | female | < 60 | 51 | 30 | 15 | 2011 | Intensive induction | yes | CR post induction | yes | no | Alive w/ relapse |
| AML65 | no | male | < 60 | 27 | 77.5 | NA | 2011 | Intensive induction | yes | CR post induction | yes | no | Alive w/ relapse |
| AML66 | yes | female | < 60 | 53 | 0 | 35 | 2017 | Intensive induction | yes | | yes | yes | Relapse mortality |
| AML67 | no | male | ≥ 60 | 72 | 70 | 80 | 2016 | Intensive induction | yes | CR post induction | yes | no | Alive w/ relapse |
| AML68 | yes | female | < 60 | 41 | NA | NA | 2015 | Intensive induction | yes | CR post induction | no | no | Alive w/o relapse |

AML49 missing due to tissue detachment of TMA core from slide prior to imaging.

**Table S2: Normal bone marrow cohort.**

| UPN | Sex | Age | Indication_biopsy | Diagnosis |
|---|---|---|---|---|
| NBM01 | Female | 71 | Suspected lymphoma | NA |
| NBM02 | Male | 51 | Suspected lymphoma | Follicular lymphoma |
| NBM03 | Female | 47 | Leukopenia | NA |
| NBM04 | Male | 62 | Suspected lymphoma | MALT lymphoma |
| NBM05 | Female | 58 | Suspected lymphoma | NA |
| NBM06 | Male | 37 | Suspected lymphoma | NA |
| NBM07 | Male | 81 | Anemia | NA |
| NBM08 | Male | 71 | Suspected lymphoma | Follicular lymphoma |
| NBM09 | Female | 69 | Anemia, thrombocytopenia | NA |
| NBM10 | Female | 37 | Suspected lymphoma | Hodgkin's lymphoma |
| NBM11 | Male | 72 | Suspected lymphoma | Non-Hodgkin's lymphoma |
| NBM12 | Male | 72 | Suspected lymphoma | NA |
| NBM13 | Female | 69 | Suspected lymphoma | Follicular lymphoma |
| NBM14 | Male | 60 | Suspected lymphoma | Follicular lymphoma |
| NBM15 | Female | 59 | Anemia | Autoimmune hemolysis |
| NBM16 | Female | 69 | Suspected lymphoma | Marginal zone lymphoma |
| NBM17 | Male | 57 | Suspected lymphoma | Follicular lymphoma |
| NBM18 | Male | 75 | Suspected lymphoma | DLBCL |
| NBM19 | Male | 42 | Suspected lymphoma | Retroperitoneal fibrosis |
| NBM20 | Male | 41 | Suspected lymphoma | NLPHL |
| NBM21 | Male | 65 | Suspected lymphoma | Follicular lymphoma |
| NBM22 | Male | 61 | Suspected lymphoma | Follicular lymphoma |
| NBM23 | Male | 79 | Suspected lymphoma | DLBCL |
| NBM24 | Male | 24 | Suspected lymphoma | PCNSL |
| NBM25 | Female | 40 | Suspected lymphoma | NSCLC |
| NBM26 | Male | 77 | Suspected lymphoma | PCNSL |
| NBM27 | Female | 63 | Suspected lymphoma | Marginal zone lymphoma |
| NBM28 | Male | 59 | Suspected lymphoma | Vitreoretinal lymphoma |
| NBM29 | Female | 82 | Suspected lymphoma | Vitreoretinal lymphoma |
| NBM30 | Female | 82 | Suspected lymphoma | NA |
| NBM31 | Female | 81 | Suspected lymphoma | Marginal zone lymphoma |
| NBM32 | Female | 56 | Suspected lymphoma | MGUS |
| NBM33 | Female | 78 | Suspected lymphoma | DLBCL |
| NBM34 | Female | 54 | Suspected lymphoma | Follicular lymphoma |
| NBM35 | Female | 59 | Suspected lymphoma | Marginal zone lymphoma |
| NBM36 | Female | 74 | Suspected lymphoma | Vitreoretinal lymphoma |
| NBM37 | Male | 50 | Suspected lymphoma | DLBCL |
| NBM39 | Male | 59 | Suspected lymphoma | DLBCL |

DLBCL, diffuse large B-cell lymphoma. NLPHL, Nodular lymphocyte-predominant Hodgkin's lymphoma. PCNSL, primary central nervous system lymphoma. NSCLC, non small-cell lung cancer. MGUS, monoclonal gammopathy of unclear significance. NBM38 missing due to tissue detachment of TMA core from slide prior to imaging.

**Table S3: Antibodies for CODEX staining.**

| Antigen | Clone | Manufacturer | Catalog No. | CODEX oligo | Working dilution | Exposure time (ms) |
|---|---|---|---|---|---|---|
| BCL2 | 124 | Cell Marque | custom | 41 | 1:50 | 150 |
| Ki67 | B56 | BD Biosciences | 556003 | 6 | 1:200 | 150 |
| NPM1 | poly | ThermoFisher | PA1-46356 | 67 | 1:200 | 300 |
| FOXP3 | 236A/E7 | Invitrogen | 14-4777-80 | 2 | 1:50 | 500 |
| IRF4 | IRF4.3E4 | Biolegend | 646402 | 56 | 1:200 | 150 |
| CD33 | 44M12D3 | Novus Biologicals | NBP2-22377 | 14 | 1:50 | 250 |
| CD31 | C31.3+..+.. | Novus Biologicals | NBP2-47785 | 68 | 1:50 | 150 |
| CD45 | 2B11 + PD7/26 | Novus Biologicals | NBP2-34528 | 145 | 1:25 | 300 |
| CD25 | 4C9 | Cell Marque | custom / 125M-14 | 24 | 1:50 | 150 |
| CD56 | MRQ-42 | Cell Marque | custom | 29 | 1:50 | 300 |
| CD57 | HCD57 | Biolegend | 322325 | 30 | 1:100 | 250 |
| CD69 | poly | Novus Biologicals | AF2359 | 36 | 1:50 | 500 |
| CD34 | QBEnd/10 | Novus Biologicals | NBP2-47909-0.1mg | 38 | 1:50 | 300 |
| CD20 | rIGEL/773 | Novus Biologicals | NBP2-54591 | 48 | 1:50 | 300 |
| ARG-1 | poly | Novus | | 52 | 1:200 | 150 |
| CD15 | MMA (BD) | BD Biosciences | 559045 | 113 | 1:100 | 150 |
| HLA-DR | EPR3692 | Abcam | ab215985 | 65 | 1:50 | 300 |
| MHC-I | EMR8-5 | Abcam | ab70328 | 15 | 1:200 | 300 |
| MMP9 | L51/82 (Biolegend) | Biolegend | 819701 | 143 | 1:100 | 250 |
| MCT | AA1 (BioLegend) | Biolegend | 369402 | 112 | 1:50 | 150 |
| CD11c | EP1347Y | Abcam | ab216655 | 49 | 1:50 | 150 |
| ICOS | D1K2T | Cell Signaling Technology | custom | 74 | 1:100 | 500 |
| CXCL13 | poly | R&D Systems | AF801 | 75 | 1:100 | 500 |
| CX3CR1 | poly | Novus Biologicals | NBP1-76949 | 7 | 1:200 | 150 |
| CD47 | poly (RD) | R&D Systems | AF4670 | 105 | 1:50 | 150 |
| CD73 | poly (Novus) | R&D Systems | AF5795 | 107 | 1:50 | 150 |
| Granzyme B | EPR20129-217 | Abcam | ab219803 | 81 | 1:200 | 100 |
| CD163 | EDHu-1 | Novus Biologicals | NB110-40686 | 45 | 1:50 | 250 |
| VISTA | D1L2G | Cell Signaling Technology | custom | 79 | 1:50 | 300 |
| pSTAT3 | 4 | BD Biosciences | custom | 5RR | 1:50 | 500 |
| GATA3 | L50-823 | Cell Marque | custom | 60 | 1:50 | 300 |
| p53 | DO-7 | Santa Cruz | sc-47698 | 25 | 1:50 | 300 |
| PAX5 | D7H5X | Cell Signaling Technology | 12709BF | 42RR | 1:100 | 150 |
| CD3 | MRQ-39 | Cell Marque | custom | 77RR | 1:100 | 300 |
| NKG2D | 1D11 | Novus | NB100-65956 | 32 | 1:50 | 300 |
| CD45RO | UCH-L1 | BioLegend | 304202 | 61 | 1:50 | 250 |
| CD8 | C8/144B | Santa Cruz Biotechnology | sc-53212 | 8 | 1:25 | 150 |
| CD71 | MRQ-48 | Cell Marque | custom / 171M-94 | 3 | 1:800 | 30 |
| CTLA4 | E2V1Z | Cell Signaling Technology | 53560S | 62 | 1:100 | 50 |
| CD4 | EPR6855 | Abcam | ab181724 | 20 | 1:50 | 250 |
| CD271 | 74902 | R&D Systems | MAB367 | 21 | 1:50 | 300 |
| PD1 | D4W2J | Cell Signaling Technology | 86163BF | 23 | 1:50 | 500 |
| CD7 | MRQ-56 | Cell Marque | custom | 63RR | 1:50 | 150 |
| Vimentin | RV202 (BD) | BD Biosciences | 550513 | 110 | 1:50 | 150 |
| CD45RA | HI100 | Biolegend | 304102 | 109 | 1:50 | 500 |
| CD68 | KP-1 | Biolegend | 916104 | 53 | 1:200 | 75 |
| CD16 | D1N9L | Cell Signaling Technology | custom | 26 | 1:100 | 150 |
| TCRb | G-11 (sc-5277) | Santa Cruz Biotechnology | sc-5277 | 116 | 1:50 | 500 |
| TCRd | H41 | Santa Cruz Biotechnology | sc-100289 | 121 | 1:50 | 300 |
| CD38 | EPR4106 | Abcam | ab176886 | 66 | 1:100 | 150 |
| CD138 | MI15 | BioLegend | 356502 | 127 | 1:50 | 150 |
| CD5 | NBP2-34583 (Novus | Novus Biologicals | NBP2-34583-0.1mg | 148RR | 1:50 | 150 |
| CD79a | JBC117 | Novus Biologicals | NBP2-34652-0.1mg | 111 | 1:50 | 300 |
| CD14 | poly (CST) | Cell Signaling Technology | 43878SF | 128 | 1:50 | 300 |
| CD11b | EPR1344 | Abcam | ab209970 | 28 | 1:50 | 150 |
| CD206 | poly | BioLegend / R&D | AF2534 | 55 | 1:200 | 150 |
| CD276 | poly (R&D) | Abcam | ab105922 | 76 | 1:50 | 150 |
| TIM3 | poly | R&D Systems | AF2365 | 46 | 1:50 | 150 |
| Perilipin | poly | Sigma Aldrich | P1873 | 70 | 1:200 | 150 |

#### Table S4: Oligos for CODEX multicycle.

| Oligo No. | Antibody oligo sequence (5'-3') | Fluorescent oligonucleotide sequences (5'-3') | Blocking component 4 (BC4) oligos (5'-3') |
|---|---|---|---|
| 2 | /mal/ATGGTTTAGGACTAC | /5fluorophoreN/GTAGTCCTAAACCAT | GTAGTCCTAAACCAT |
| 3 | /mal/TACTCCTCGCCG | /5fluorophoreN/CGGCGAGGAGTA | CGGCGAGGAGTA |
| 5 | /mal/TCTCCCATTAGTCGG | /5fluorophoreN/CCGACTAATGGGAGA | CCGACTAATGGGAGA |
| 6 | /mal/TGGATGTGTTACGAT | /5fluorophoreN/ATCGTAACACATCCA | ATCGTAACACATCCA |
| 7 | /mal/CGCTAAGATATTCTAAG | /5fluorophoreN/CTTAGAATATCTTAGCG | CTTAGAATATCTTAGCG |
| 8 | /mal/CGCAGATGAATATTC | /5fluorophoreN/GAATATTCATCTGCG | GAATATTCATCTGCG |
| 14 | /mal/AGATTCAGTCTCTCG | /5fluorophoreN/CGAGAGACTGAATCT | CGAGAGACTGAATCT |
| 15 | /mal/CTGTAATAGGCACTA | /5fluorophoreN/TAGTGCCTATTACAG | TAGTGCCTATTACAG |
| 20 | /mal/ATGTAGGATGGTCTC | /5fluorophoreN/GAGACCATCCTACAT | GAGACCATCCTACAT |
| 21 | /mal/CGTGCCGTTTCAC | /5fluorophoreN/GTGAAACGGCACG | GTGAAACGGCACG |
| 23 | /mal/GGTTTCCTCAGACAC | /5fluorophoreN/GTGTCTGAGGAAACC | GTGTCTGAGGAAACC |
| 24 | /mal/ATAAGGGCTCATTGT | /5fluorophoreN/ACAATGAGCCCTTAT | ACAATGAGCCCTTAT |
| 26 | /mal/CACTTGTCTAACCAA | /5fluorophoreN/TTGGTTAGACAAGTG | TTGGTTAGACAAGTG |
| 28 | /mal/TGGTCCACTAACGTA | /5fluorophoreN/TACGTTAGTGGACCA | TACGTTAGTGGACCA |
| 29 | /mal/ATAGGGCATTTGAAG | /5fluorophoreN/CTTCAAATGCCCTAT | CTTCAAATGCCCTAT |
| 30 | /mal/CACATGAGCGAATCA | /5fluorophoreN/TGATTCGCTCATGTG | TGATTCGCTCATGTG |
| 32 | /mal/TACCAAATCCTGATG | /5fluorophoreN/CATCAGGATTTGGTA | CATCAGGATTTGGTA |
| 36 | /mal/ACCTACACAATGCTA | /5fluorophoreN/TAGCATTGTGTAGGT | TAGCATTGTGTAGGT |
| 38 | /mal/GCGTCTACTTATAAG | /5fluorophoreN/CTTATAAGTAGACGC | CTTATAAGTAGACGC |
| 41 | /mal/TGTATGAGTAGTAATCT | /5fluorophoreN/AGATTACTACTCATACA | AGATTACTACTCATACA |
| 42 | /mal/TCTAAGTCAGAGAGC | /5fluorophoreN/GCTCTCTGACTTAGA | GCTCTCTGACTTAGA |
| 45 | /mal/GCCAGAATGCCA | /5fluorophoreN/TGGCATTCTGGC | TGGCATTCTGGC |
| 46 | /mal/GACTCGAACCTGAG | /5fluorophoreN/CTCAGGTTCGAGTC | CTCAGGTTCGAGTC |
| 48 | /mal/GCACGGCAAAGTG | /5fluorophoreN/CACTTTGCCGTGC | CACTTTGCCGTGC |
| 49 | /mal/ATAACGCCTCGTATC | /5fluorophoreN/GATACGAGGCGTTAT | GATACGAGGCGTTAT |
| 52 | /mal/ACCAGGTGATTGCAT | /5fluorophoreN/ATGCAATCACCTGGT | ATGCAATCACCTGGT |
| 55 | /mal/AGGTCAACTCGCAC | /5fluorophoreN/GTGCGAGTTGACCT | GTGCGAGTTGACCT |
| 56 | /mal/GGTCACATGGTCGTT | /5fluorophoreN/AACGACCATGTGACC | AACGACCATGTGACC |
| 60 | /mal/ACAAAACTGCTGTCG | /5fluorophoreN/CGACAGCAGTTTTGT | CGACAGCAGTTTTGT |
| 61 | /mal/TCTTATTCCCGAATA | /5fluorophoreN/TATTCGGGAATAAGA | TATTCGGGAATAAGA |
| 62 | /mal/TAGGGGAACAGGTTG | /5fluorophoreN/CAACCTGTTCCCCTA | CAACCTGTTCCCCTA |
| 63 | /mal/AATTAGCTTAAGAGAGT | /5fluorophoreN/ACTCTCTTAAGCTAATT | ACTCTCTTAAGCTAATT |
| 65 | /mal/GATAAATATTTTACAGAGT | /5fluorophoreN/ACTCTGTAAAATATTTATC | ACTCTGTAAAATATTTATC |
| 66 | /mal/TACGTGCTTTGGTT | /5fluorophoreN/AACCAAAGCACGTA | AACCAAAGCACGTA |
| 67 | /mal/GACGACGAAGGC | /5fluorophoreN/GCCTTCGTCGTC | GCCTTCGTCGTC |
| 68 | /mal/CTTCTTGTGGAACC | /5fluorophoreN/GGTTCCACAAGAAG | GGTTCCACAAGAAG |
| 70 | /mal/AACCAAACTGACCG | /5fluorophoreN/CGGTCAGTTTGGTT | CGGTCAGTTTGGTT |
| 74 | /mal/ATTTGCTTCGACGA | /5fluorophoreN/TCGTCGAAGCAAAT | TCGTCGAAGCAAAT |
| 75 | /mal/CGCTTGGGTGTTTA | /5fluorophoreN/TAAACACCCAAGCG | TAAACACCCAAGCG |
| 76 | /mal/CTGTCGGTCGTCA | /5fluorophoreN/TGACGACCGACAG | TGACGACCGACAG |
| 77 | /mal/ATTTCAACAAATATTGTT | /5fluorophoreN/AACAATATTTGTTGAAAT | AACAATATTTGTTGAAAT |
| 79 | /mal/GCCGGAAGTGGT | /5fluorophoreN/ACCACTTCCGGC | ACCACTTCCGGC |
| 81 | /mal/CAAGGAACTACCGA | /5fluorophoreN/TCGGTAGTTCCTTG | TCGGTAGTTCCTTG |
| 105 | /mal/ATTTCCAAGGGC | /5fluorophoreN/GCCCTTGGAAAT | GCCCTTGGAAAT |
| 107 | /mal/CTTTACAACGGG | /5fluorophoreN/CCCGTTGTAAAG | CCCGTTGTAAAG |
| 109 | /mal/TACCGTGGTCAA | /5fluorophoreN/TTGACCACGGTA | TTGACCACGGTA |
| 110 | /mal/CCAACTGGGTTA | /5fluorophoreN/TAACCCAGTTGG | TAACCCAGTTGG |
| 111 | /mal/GCTGGCTTCAAA | /5fluorophoreN/TTTGAAGCCAGC | TTTGAAGCCAGC |
| 112 | /mal/ATTCTGCAAGGC | /5fluorophoreN/GCCTTGCAGAAT | GCCTTGCAGAAT |
| 113 | /mal/CCGGTTGCTAAA | /5fluorophoreN/TTTAGCAACCGG | TTTAGCAACCGG |
| 116 | /mal/CCCGTTTGAAGA | /5fluorophoreN/TCTTCAAACGGG | TCTTCAAACGGG |
| 121 | /mal/TTTACAAGCCGG | /5fluorophoreN/CCGGCTTGTAAA | CCGGCTTGTAAA |
| 127 | /mal/TTTGAACGAGCC | /5fluorophoreN/GGCTCGTTCAAA | GGCTCGTTCAAA |
| 128 | /mal/TTCCAGAGGTCA | /5fluorophoreN/TGACCTCTGGAA | TGACCTCTGGAA |
| 143 | /mal/TTGACCAGAGCT | /5fluorophoreN/AGCTCTGGTCAA | AGCTCTGGTCAA |
| 145 | /mal/ATGCTGGATCCA | /5fluorophoreN/TGGATCCAGCAT | TGGATCCAGCAT |
| 148 | /mal/ATACCTTGGGCA | /5fluorophoreN/TGCCCAAGGTAT | TGCCCAAGGTAT |

**Table S5: CODEX multicycle layout.**

| Cyc | Channel 1 | Exposure Time (ms) | Alexa488 | Exposure Time (ms) | ATTO550 | Oligo | Clone | Dilution | Exposure Time (ms) | Alexa647 | Oligo | Clone | Dilution | Exposure Time (ms) |
|---|---|---|---|---|---|---|---|---|---|---|---|---|---|---|
| 1 | Hoechst 33342 | 4 | Blank | 2 | Blank | Blank | - | - | 150 | Blank | Blank | - | - | 150 |
| 2 | Hoechst 33342 | 4 | Empty | 2 | BCL2 | A41 | 124 | 1:50 | 150 | pSTAT3 | A5RR | 4 | 1:50 | 500 |
| 3 | Hoechst 33342 | 4 | Empty | 2 | Ki67 | A6 | B56 | 1:200 | 150 | GATA3 | A60 | L50-823 | 1:50 | 300 |
| 4 | Hoechst 33342 | 4 | Empty | 2 | NPM1 | A67 | poly | 1:200 | 300 | p53 | A25 | DO-7 | 1:50 | 300 |
| 5 | Hoechst 33342 | 4 | Empty | 2 | FOXP3 | A2 | 236A/E7 | 1:50 | 500 | PAX5 | A42RR | D7H5X | 1:100 | 150 |
| 6 | Hoechst 33342 | 4 | Empty | 2 | IRF4 | A56 | IRF4.3E4 | 1:200 | 150 | CD3 | A77RR | MRQ-39 | 1:100 | 300 |
| 7 | Hoechst 33342 | 4 | Empty | 2 | CD33 | A14 | 44M12D3 | 1:50 | 250 | NKG2D | A32 | 1D11 | 1:50 | 300 |
| 8 | Hoechst 33342 | 4 | Empty | 2 | CD31 | A68 | C31.3+..+.. | 1:50 | 150 | CD45RO | A61 | UCH-L1 | 1:50 | 250 |
| 9 | Hoechst 33342 | 4 | Empty | 2 | CD45 | 145 | 2B11 + PD7/. | 1:25 | 300 | CD8 | A8 | C8/144B | 1:25 | 150 |
| 10 | Hoechst 33342 | 4 | Empty | 2 | CD25 | A24 | 4C9 | 1:50 | 150 | CD71 | A3 | MRQ-48 | 1:800 | 30 |
| 11 | Hoechst 33342 | 4 | Blank | 2 | CD56 | A29 | MRQ-42 | 1:50 | 300 | CTLA4 | A62 | E2V1Z | 1:100 | 50 |
| 12 | Hoechst 33342 | 4 | Empty | 2 | CD57 | A30 | HCD57 | 1:100 | 250 | CD4 | A20 | EPR6855 | 1:50 | 250 |
| 13 | Hoechst 33342 | 4 | Empty | 2 | CD69 | A36 | poly | 1:50 | 500 | CD271 | A21 | 74902 | 1:50 | 300 |
| 14 | Hoechst 33342 | 4 | Empty | 2 | CD34 | A38 | QBEnd/10 | 1:50 | 300 | PD1 | A23 | D4W2J | 1:50 | 500 |
| 15 | Hoechst 33342 | 4 | Empty | 2 | CD20 | A48 | rIGEL/773 | 1:50 | 300 | CD7 | A63RR | MRQ-56 | 1:50 | 150 |
| 16 | Hoechst 33342 | 4 | Empty | 2 | ARG-1 | A52 | poly | 1:200 | 150 | Vimentin | 110 | RV202 (BD) | 1:50 | 150 |
| 17 | Hoechst 33342 | 4 | Empty | 2 | CD15 | 113 | MMA (BD) | 1:100 | 150 | CD45RA | 109 | HI100 | 1:50 | 500 |
| 18 | Hoechst 33342 | 4 | Empty | 2 | HLA-DR | A65 | EPR3692 | 1:50 | 300 | CD68 | A53 | KP-1 | 1:200 | 75 |
| 19 | Hoechst 33342 | 4 | Empty | 2 | MHC-I | A15 | EMR8-5 | 1:200 | 300 | CD16 | A26 | D1N9L | 1:100 | 150 |
| 20 | Hoechst 33342 | 4 | Empty | 2 | MMP9 | 143 | L51/82 (Biol | 1:100 | 250 | TCRb | 116 | G-11 (sc-527 | 1:50 | 500 |
| 21 | Hoechst 33342 | 4 | Empty | 2 | MCT | 112 | AA1 (BioLeg | 1:50 | 150 | TCRd | 121 | H41 | 1:50 | 300 |
| 22 | Hoechst 33342 | 4 | Empty | 2 | CD11c | 49 | EP1347Y | 1:50 | 150 | CD38 | A66 | EPR4106 | 1:100 | 150 |
| 23 | Hoechst 33342 | 4 | Empty | 2 | ICOS | A74 | D1K2T | 1:100 | 500 | CD138 | 127 | MI15 | 1:50 | 150 |
| 24 | Hoechst 33342 | 4 | Empty | 2 | CXCL13 | A75 | poly | 1:100 | 500 | CD5 | 148RR | NBP2-34583 | 1:50 | 150 |
| 25 | Hoechst 33342 | 4 | Empty | 2 | CX3CR1 | A7 | poly | 1:200 | 150 | CD79a | 111 | JBC117 | 1:50 | 300 |
| 26 | Hoechst 33342 | 4 | Empty | 2 | CD47 | 105 | poly (RD) | 1:50 | 150 | CD14 | 128 | poly (CST) | 1:50 | 300 |
| 27 | Hoechst 33342 | 4 | Empty | 2 | CD73 | 107 | poly (Novus) | 1:50 | 150 | CD11b | A28 | EPR1344 | 1:50 | 150 |
| 28 | Hoechst 33342 | 4 | Empty | 2 | Granzyme B | A81 | EPR20129-2: | 1:100 | 100 | CD206 | A55 | poly | 1:200 | 150 |
| 29 | Hoechst 33342 | 4 | Empty | 2 | CD163 | A45 | EDHu-1 | 1:50 | 250 | CD276 | 76 | poly (R&D) | 1:50 | 150 |
| 30 | Hoechst 33342 | 4 | Empty | 2 | VISTA | A79 | D1L2G | 1:50 | 300 | TIM3 | 46 | poly | 1:50 | 150 |
| 31 | Hoechst 33342 | 4 | Empty | 2 | Empty | Empty | - | - | 150 | Perilipin | A70 | poly | 1:200 | 150 |
| 32 | Hoechst 33342 | 4 | Empty | 2 | Empty | Empty | - | - | 150 | DRAQ5 | | | | 150 |
| 33 | Hoechst 33342 | 4 | Empty | 2 | Blank | Blank | - | - | 150 | Blank | Blank | - | - | 150 |

**Table S6: Cell types and marker expression patterns**

| Category | Cell type | Antigen expression pattern |
|---|---|---|
| Lymphoid | B-cells | LIN+ CD45+ PAX5+ CD79a+ CD20+ |
| Lymphoid | CD4+ T-cells | LIN+ CD45+ CD3+ CD4+ CD8- |
| Lymphoid | CD4+ T-cells (PD1+) | LIN+ CD45+ CD3+ CD4+ CD8- PD1+ |
| Lymphoid | CD8+ T-cells | LIN+ CD45+ CD3+ CD4+ CD8- |
| Myeloid | DCs | LIN+ CD11c+ HLA-DR+ |
| Stromal | Endothelium | LIN- CD34hi CD31+ |
| Stromal | Endothelium (CD73+ CD206+) | LIN- CD34lo CD73+ CD206hi |
| Myeloid | Eosinophils | Identified by unspecific Fc-binding, correlatec |
| Erythroid | Erythroid | LIN- CD33- CD34- CD71hi Ki67hi |
| Lymphoid | FOXP3+ Tregs | LIN+ CD45+ CD3+ CD4+ FOXP3+ |
| Lymphoid | yd T-cells | LIN+ CD45+ CD3+ TCRd+ |
| Myeloid | Immature myeloid cells (CD11b+) | LIN+ CD33+ CD15lo CD11bhi |
| Myeloid | Immature myeloid cells (CD15+) | LIN+ CD33+ CD15+ Arg1+ CD11b- |
| Myeloid | Immature myeloid cells (GZMB+) | LIN+ CD33+ GZMBhi |
| Myeloid | Immature myeloid cells (mixed) | LIN+ CD33+ CD11c+ CD14+ VISTA+ CD15lo A |
| Myeloid | Blasts | LIN- CD33+ CD47+ |
| Myeloid | Macrophages | LIN+ CD3- CD4+ CD16+ CD68+ CD163+ |
| Myeloid | Mast cells | MCThigh |
| Myeloid | Mature myeloid cells | CD33+ CD11b+ CD15+ Arg1+ MMP9+ |
| Stromal | MSC (CD271+) | LIN- CD73+ CD271+ CD206lo |
| Stromal | MSC (pSTAT3+) | LIN- CD73+ CD271+ pSTAT3+ |
| Lymphoid | NK cells | LIN+ CD45+ CD3- CD57+ |
| Stromal | Osteoblasts | LIN- CD73+ CD271+ CD56hi |
| Lymphoid | Plasma cells | LIN+ CD79a+ CD38+ IRF4+ CD138+ CD20- |
| Stromal | Schwann cells | LIN- VIM+ CD57hi |

### SUPPLEMENTAL MATERIALS & METHODS

#### Patient cohort

176 patients diagnosed with acute myeloid leukemia (AML) with a *Nucleophosmin 1* (*NPM1*) mutation were treated at our center between 2005 and 2023. Archival formalin-fixed paraffin-embedded (FFPE) trephine biopsies from 67 patients at first diagnosis were available. Clinicopathological data including long-term follow-up from all patients were extracted from clinical and pathological reports (**Table S1**). For correlative analysis, a cohort of 38 age- and sex-matched individuals with normal bone marrow (NBM) who received trephine biopsy for lymphoma staging at first diagnosis or suspected hematological disorder was composed and clinically annotated. Only samples without any pathological findings upon comprehensive diagnostic hematopathology assessment were included. Informed consent was obtained in accordance with the Declaration of Helsinki. This research was approved by the Ethics Committee of the University of Tübingen (099/2023BO2).

#### Construction of tissue microarrays (TMAs)

FFPE tissue blocks were retrieved from the tissue archive at the Department of Pathology, University Hospital Tübingen, Germany. For the study of *NPM1*-mutant AML, three independent tissue microarrays (TMAs) were generated, containing one to six cores per patient (median, 3 cores per patient). TMA cores containing representative bone marrow (BM) regions were annotated on corresponding hematoxylin and eosin (H&E)-stained sections under the supervision of a board-certified surgical pathologist (C.M.S.). All TMAs contain 1 mm diameter cores and were assembled using a TMA Grand Master automated tissue microarrayer (3DHistech). TMAs were sectioned at 4 µm thickness, mounted on Superfrost slides (Firma Langenbrinck) and stored at 4°C until analysis. TMA sections were stained and processed for CODEX (see below). After CODEX imaging, TMA sections were stained with H&E, and digitized using a Pannoramic P250 digital slide scanner (3DHistech). For NBM, two independent TMAs were generated and sectioned following the same procedures. Tonsil cores were included in all TMAs and co-imaged and analyzed as technical controls.

#### Buffers and solutions

All buffers were prepared according to the published CODEX protocols[1–3], sterile-filtered through 0.2 µm membranes (500 mL units), and held at room temperature (RT) unless noted.

Staining solution 1 (S1): 5 mM EDTA (Sigma), 0.5% w/v bovine serum albumin (BSA, Sigma) and 0.02% w/v NaN3 (Sigma) in PBS (Thermo Fisher Scientific); storage at 4°C.

Staining solution 2 (S2): 61 mM $Na_2HPO_4 \cdot 7 H_2O$ (Sigma), 39 mM $NaH_2PO_4$ (Sigma) and 250 mM NaCl (Sigma) in a 1:0.7 v/v solution of S1 and doubly-distilled $H_2O$ ($ddH_2O$); final pH 6.8- 7.0.

Staining solution 4 (S4): 0.5 M NaCl in S1.

TE buffer: 10 mM Tris pH 8.0 and 1 mM EDTA in $ddH_2O$.

Tris stock (50 mM, pH 7.2 at RT) for buffer C: Trizma HCl and Trizma base combined in $ddH_2O$ per the manufacturer's mixing table (Sigma).

Buffer C (for conjugation): 150 mM NaCl, 2 mM Tris, pH 7.2 and 1 mM EDTA in $ddH_2O$.

CODEX 2.0 buffer (H2): 150 mM NaCl, 10 mM Tris pH 7.5 (Teknova), 10 mM $MgCl_2 \cdot 6 H_2O$ (Sigma), 0.1% w/v Triton X-100 (Sigma) and 0.02% w/v $NaN_3$ in $ddH_2O$.

Blocking reagent 1 (B1): 1 mg/ml mouse IgG (Sigma) in S2.

Blocking reagent 2 (B2): 1 mg/ml rat IgG (Sigma) in S2.

Blocking reagent 3 (B3): Sheared salmon sperm DNA, 10 mg/ml in $H_2O$ (Thermo Fisher, #AM9680).

Blocking component 4 (BC4) and BC5: all 79 non-modified CODEX barcode oligonucleotides pooled in TE buffer, 0.5 mM each.

BS3 stock: BS3 (Thermo Fisher, #21580) dissolved to 200 mg/mL in anhydrous DMSO (Sigma), aliquoted at 15 µL and held at −20 °C.

TCEP solution: 0.5 M 2.5 mM Tris (2-carboxyethyl) phosphine hydrochloride (TCEP; Merck / Supelco, #646547-10G ) in $ddH_2O$, pH 7.0.

Rendering buffer: 20% DMSO (v/v) in H2 buffer.

Bleaching solution: 4.5% (w/v) $H_2O_2$ and 20 mM NaOH in PBS.

**CODEX DNA-conjugated antibodies**

Mono- and polyclonal antibodies (purified, carrier-free; Table S3) were coupled to maleimide-modified DNA oligonucleotides (Biomers; Table S4) following established protocols[1,2]. In brief, antibody was concentrated on an Amicon Ultra 50-kDa centrifugal filter unit (Millipore Sigma / Merck, #UFC505096). Interchain disulfides were then reduced for 30 min at RT in PBS supplemented with 2.5 mM TCEP (Merck / Supelco, #646547-10G) and 2.5 mM EDTA, pH 7.0. Each barcode oligonucleotide was dissolved in buffer C adjusted to 400 mM NaCl, applied to the retained antibody on the filter at an oligonucleotide:antibody mass ratio of 2:1, and allowed to react for 2 h at RT. No less than 100 µg of antibody was used per conjugation. Unbound oligonucleotide was removed by washing, and the conjugate was recovered in a PBS-based antibody stabilizer (Candor Bioscience, #131 050) supplemented with 500 mM NaCl, 5 mM EDTA and 0.1% (w/v) $NaN_3$ (Sigma-Aldrich, #S8032).

#### CODEX antibody screening, validation and titration

Antibodies were first screened and validated in CODEX single-stainings on tonsil tissue or a multi-tissue TMA, with cross-validation by manual IHC as described previously[1]. Briefly, after antibody staining and fixation, 100 nM of fluorescent DNA probe (Biomers) was added to the tissue in rendering buffer, containing 0.7 mg/ml sheared salmon sperm DNA, and was incubated at RT for 5-10 min, followed by washing with rendering buffer and S4 buffer. Coverslips were mounted onto glass slides, dried in the dark and imaged on a BZ-X810 inverted fluorescence microscope (Keyence). All validation was performed by or under the supervision of a board-certified surgical pathologist (C.M.S.) and confirmed with online databases (https://www.proteinatlas.org/, https://www.pathologyoutlines.com/) and published literature. For each validated conjugate, the staining dilution was optimized by titration, starting from 1:100 and adjusted according to signal-to-noise.

#### Immunohistochemistry

Sections were cut to 2.5 μm thickness and placed on TOMO® adhesive glass slides (Roche). H&E stained sections were obtained from each FFPE block. Deparaffinization, rehydration, and Heat-induced epitope retrieval were performed on a Ventana Benchmark XT tissue stainer (Roche) using the proprietary protocol, blocking solution, antibody diluent, washing agent and detection kit from Roche. Antibodies were diluted in 100 μL antibody diluent at dilutions ranging from 1:10 to 1:10,000 and sections were stained for two hours. After washing, bound antibodies were visualized using the proprietary substrate chromogen detection kit according to the manufacturer's instructions. Sections were counterstained with hematoxylin, followed by dehydration, mounting, and digitization using a Pannoramic P250 digital slide scanner (3DHistech).

#### CODEX multiplex tissue staining

For deparaffinization, slides were baked at 70°C for at least 1 h, followed by immersion in fresh xylene for 30 min. Sections were rehydrated in descending concentrations of ethanol (100% twice, 96%, 70%, 50%, 30%, ddH2O twice; each step for 5 min). Heat-induced epitope retrieval was performed in a pressure cooker (MD1000, Rommelsbacher) using Dako target retrieval solution, pH 9 (1x; Agilent, #S2367) at 108°C for 20 min. After cooling to RT for 30 min, autofluorescence quenching by photochemical bleaching was performed as reported previously[4]. Slides were submerged in bleaching solution in Petri dishes for a total of 90 min, with transfer to a dish with fresh bleaching solution after 45 min. For the complete duration, dishes were sandwiched between two broad spectrum LED light sources (Aibecy A5 Ultra Bright 25,000 Lux LED Pads). This was followed by 3 x 3 min washing in 1 x PBS and 10 min in 1x TBS wash buffer with Tween 20 (Cell Marque) on a shaker. Tissues were encircled with a UV-curable liquid plastic barrier (Bondic), and nonspecific binding was blocked for 90 min at RT using 100 µL of blocking buffer [S2 buffer containing B1 (1:20), B2 (1:20), B3 (1:20), and BC4 and BC5 (1:15)]. For each slide, DNA-conjugated antibodies were added to 50 µL of blocking buffer on a 50-kDa filter unit, concentrated by spinning at 12,000 x *g* for 8 min, and resuspended in blocking buffer to a final volume of 150 µL. This antibody cocktail was then added to the slide, and staining was performed in a sealed humidity chamber overnight at 4°C on a shaker. After staining, slides were washed for 4 min in S2 and fixed in S4 containing 1.6% paraformaldehyde for 10 min, followed by three washes in PBS. Then, slides were incubated in 100% ice-cold methanol for 5 min, followed by three washes in PBS. Fresh BS3 fixative was prepared immediately before final fixation by thawing and diluting one 15 µL aliquot of BS3 in 1 mL PBS. Final fixation was performed at RT for 20 min, followed by three washes in PBS. Thereafter, slides were stored in S4 in a conical tube at 4°C for up to two weeks, or further processed for imaging.

**CODEX multicycle reaction and image acquisition**

Staining and multicycle imaging were performed with in-house-prepared buffers (see above) rather than the proprietary Akoya reagent kit, as described previously[1,2]. For imaging using the PhenoCycler Fusion 2.0 instrument (Akoya Biosciences / Quanterix), flow cells were assembled by stamping adhesive flow cell covers (Akoya Biosciences / Quanterix) onto slides with stained tissue after removal of the liquid plastic barrier and gentle wiping of the slide surface to be attached to the adhesive cover. Stamping was performed using the proprietary device from Akoya Biosciences / Quanterix. In case of prior storage in S4, slides were washed in 1x PBS for 10 min before flow cell assembly. To prevent trapping of bubbles, flow cells were submerged in a conical tube containing H2 buffer immediately. The flow cells were then first wiped with Kimwipe soaked in ddH2O and again wiped with a wipe soaked in 70% ethanol to remove dust and particles from the slide bottom. Next, flow cells were mounted on the flow cell slide carrier, the carrier was loaded onto the automated stage, followed by priming and rinsing according to the steps specified by the PhenoCycler Fusion 2.0 control software. Trapped bubbles were gently dislocated from the flow cells by angulation, tapping, or if necessary, manual flushing according to manufacturer recommendations. Exposure times, z-plane number and focus were determined automatically by the PhenoCycler Fusion 2.0 control software. Imaging was performed with a 20×/0.75 NA objective, and data were exported in QPTIFF format.

Fluorescent oligonucleotides (Table S4; concentration: 400 nM) were aliquoted in Corning™ black 96-well plates in 250 µL H2 buffer containing Hoechst nuclear stain (1:400) and 0.5

mg/ml sheared salmon sperm DNA, according to the multi-cycle reaction panels. Black plates were sealed with aluminum sealing film (VWR Scientific) and kept at RT during the multi-cycle reaction. The final concentration of fluorescent oligonucleotides in the tissue-containing imaging well corresponded to 80 nM (1:5 dilution in rendering buffer).

Data were acquired in two AML batches (first batch containing NPM1-mutant AML TMA 1, second batch containing NPM1-mutant AML TMAs 2 and 3) and one normal bone marrow (NBM) batch (containing NBM TMAs 1 and 2).

**Computational image processing**

Raw TIFF files were deconvolved and background-subtracted on the Enable Medicine cloud platform (https://www.enablemedicine.com/). Per-antibody staining quality was inspected region by region in the Enable Medicine Visualizer (https://app.enablemedicine.com/portal/visualizer). Nuclei were segmented with DeepCell/Mesmer[5] on the Hoechst channel within the same platform. To approximate cell boundaries, nuclear masks were expanded over nine iterations; at each iteration, unassigned pixels were probabilistically allocated to an adjacent mask, weighted by how many of the eight surrounding pixels already carried that label. For every whole-cell mask, mean marker intensities and centroid coordinates were tabulated and exported as CSV files.

**Unsupervised clustering and cluster validation**

Cells within each region were normalized across individual markers using single-z normalization. Cells were retained if at least three markers, including Hoechst and DRAQ5, had a z-score > 0, while cells with more than 35 markers with a z-score > 0 were excluded. Clustering was performed using the Seurat workflow with the following modifications. Region-wise single-z-score normalization was applied across markers prior to scaling, and the number of principal components (PCs) used for downstream analysis was determined using a data-driven elbow criterion.

Leiden clustering of the NBM dataset was performed at a resolution of 5, and the resulting clusters were manually annotated based on visual inspection of marker expression using the Enable Medicine visualizer, informed by normalized marker expression levels. The AML dataset from the first set of TMAs was filtered in the same manner as the NBM dataset, and marker expression was z-score normalized within each region. CTs identified in both datasets were subsequently combined and processed together using Seurat, including scaling, PC-analysis-based dimensionality reduction, and Leiden clustering at a resolution of 1. For each Leiden cluster in the combined NBM/AML dataset, the most abundant previously annotated healthy CT was used to pre-annotate the cluster. Pre-annotated clusters were subsequently assigned to coarse CT categories comprising lymphoid, myeloid, erythroid, mesenchymal, and artifact populations. Sub-clustering was performed for each of these pre-annotated coarse CT groups for multiple rounds and fine clusters (two levels, l1 and l2) were annotated based on marker expression and visual inspection.

Clusters that retained mixed CTs were subsequently re-annotated using supervised classification. Seurat's FindTransferAnchors function was applied separately at the coarse and fine l1 and l2 annotation levels, using successfully annotated cells as the reference and 15 PCs.

CT annotations were considered concordant when predictions agreed across all annotation levels or when the l1 and l2 predictions agreed on a CT that was compatible with the corresponding coarse lineage annotation based on CT hierarchy. Cells not fulfilling these concordance criteria were classified as unspecific.

Predicted CD4+ or CD8+ T-cells were reassigned as CD4+ T-cells if normalized CD4 expression exceeded normalized CD8 expression, and as CD8+ T-cells if normalized CD8 expression exceeded normalized CD4 expression. In AML, CD4+ T-cells with normalized FOXP3 expression >1 were assigned as FOXP3+ Tregs, confirmed by visual inspection of annotated cells. In addition, in the NBM dataset, predicted mature and immature B-cells were reassigned based on normalized CD20 expression, with cells showing CD20 expression $< 0.5$ assigned as immature B-cells and cells showing CD20 expression $> 0.5$ assigned as mature B-cells.

Cutoffs were selected from a range of candidate thresholds based on visual inspection. Predicted CTs were visually validated using the Enable Medicine visualizer, and CT annotations were adjusted where necessary.

Cells from a second imaging batch containing one AML TMA were filtered and normalized as described above. CTs were annotated using supervised classification based on the annotated cells from the first imaging batch containing 2 AML TMAs using the shared markers. CT predictions were considered concordant when the l1 and l2 predictions agreed on the CT and were compatible with the corresponding coarse lineage annotation based on the predefined CT hierarchy; otherwise, cells were classified as unspecific. Predicted CD4+ and CD8+ T-cells were reassigned as described above, and FOXP3 Tregs were identified within the CD4+ T-cell population using the same cutoff as described above. Cells classified as unspecific, macrophages, eosinophils, endothelial cells, osteoblasts, or CD4+ T-cells were further subclustered for one or multiple rounds and annotated based on marker expression and visual inspection using Enable Medicine to increase sensitivity and specificity of the final annotation. NBM cells were annotated at higher resolution than used in the final analysis. Because all analyses in this study are comparative with AML, only CTs matched to the AML annotation depth were used downstream.

##### Manual identification of adipocytes and megakaryocytes

Adipocytes and megakaryocytes (MKs) are poorly captured by nuclear segmentation because their nuclear signal is faint, irregular or multilobed, as noted in our previous work[6]. Both were therefore annotated by hand: each adipocyte and MK within every region was marked in the Enable Medicine Visualizer on the basis of its marker signal (perilipin and CD31, respectively) together with its Hoechst nuclear position. Since the panel contained no MK-restricted marker, MKs were identified by combining membranous CD31 positivity with their characteristic morphology, large cell size and a single large, lobulated or multilobed nucleus. Centroids of these annotations were computed and merged into the spatial dataset. Fragments of over-segmented MKs generated by clustering were excluded.

##### Cellular neighborhoods

Spatial neighborhoods were inferred as described previously[1]. Briefly, for each cell, the 20 nearest neighboring cells were identified within each region using Euclidean distances between spatial coordinates, excluding the query cell itself. Neighbors were mapped to their CT annotations, and each neighborhood was represented by a vector containing the number of neighboring cells of each annotated CT, with zero counts assigned to absent CTs. Neighborhood composition vectors were pooled across regions and clustered.

Partitioning around medoids (PAM) was performed on a random sample of 20,000 neighborhoods selected without replacement, using Euclidean distance and a predefined number of 10 clusters. All neighborhoods were subsequently assigned to their nearest medoid in CT composition space.

##### Quantification of pairwise CT proximity

For each tissue region, the raw physical distance between two CTs was quantified as the mean nearest-neighbor distance, computed symmetrically in both query directions (each type's cells to their nearest neighbor of the other type) and averaged between directions. For CT self-association, the same statistic used each cell's nearest other cell of the same type ($k=2$, since the first neighbor under this formulation is the cell itself, at distance 0). Per-pair values were aggregated across TMA cores by taking the mean of these per-region mean-minimum distances.

Density-corrected proximity between CT pairs was assessed using a region-conditional permutation test. For each tissue region, the observed statistic was the same bidirectional mean nearest-neighbor distance described above. For cross-type pairs, a null distribution was built by shuffling CT identity 500 times between the two focal types only, holding all cell positions fixed and preserving the true count of each type in every shuffle. For CT self-association, the null instead compared each type's self-distance to that of a random, identically-sized subset of the other (non-focal-type) cells, reflecting what would be expected under no self-clustering at all. Both constructions yield a null mean and standard deviation, and density correction was quantified as a z-score, z = (observed − null mean) / null SD. This z-score is the corrected proximity metric reported throughout. A CT (or, for cross-type pairs, both types) was only tested in a given TMA core if it had at least 3 cells present there; cores below this threshold were excluded before aggregation.

To compare raw proximity between datasets, raw physical-distance $\log_2$ fold-change was calculated. To compare density-corrected proximity, z-scores were computed identically and independently in each cohort, and the difference (z-delta = z_AML − z_NBM) was used as the density-corrected cross-cohort proximity contrast.

**Analysis of CT – CN enrichment and CT – CT co-enrichment and association with outcomes**

For analysis of enrichment for CTs in CNs, one ordinary least-squares model $y_{c,n} \sim y_c + b_n$ was fit separately for each (CT x CN) pair across patients, and enrichment beyond patient-level abundance was taken as that model's residual.

Counts from multiple cores of the same patient were pooled (summed) before computing fractions, not averaged per core. Here, $y_{c,n} = \log(N_{c,n} / N_n + 10^{-3})$ is the log-fraction of CT c among all cells in neighborhood n (per patient); $y_c = \log(N_c / N_{total} + 10^{-3})$ is that patient's systemic log-fraction of CT c, where $N_{total}$ is the patient's total cell count; and $b_n = \log(1 + N_n / K)$, where K is the patient's number of TMA cores, was included as a fitted covariate (not an offset) to adjust for neighborhood size. A fixed pseudocount of $10^{-3}$ accommodates zero counts. Only (CT × CN) pairs with ≥5 patients having non-zero counts were retained. Per-patient z-scored residuals were used as Cox predictors.

For analysis of co-localization of two CTs across CNs within a patient, the per-patient Spearman ρ of the two CTs' raw fractions (each CT's fraction of all cells within a given neighborhood) across ≥ 4 neighborhoods was computed as a continuous co-enrichment score. This score was z-scored and used as Cox predictor.

**Visualization of CT networks based on distance and co-enrichment**

Raw distance CT networks were visualized using non-metric multidimensional scaling proximity maps. CT node position was determined by non-metric multidimensional scaling (MASS::isoMDS, Kruskal stress minimization) on a pairwise dissimilarity matrix built from the raw mean nearest-neighbor distance between each pair of CTs (mean across TMA cores). Grey reference edges connect each CT to its four nearest neighbors by raw distance, plus any additional partner whose distance falls within 25% of that anchor's own minimum-to-maximum distance range beyond its closest neighbor.

Co-enrichment CT networks were visualized with force-directed proximity networks. CT pairs with co-enrichment $\rho > 0.10$ or $\rho < -0.10$ were retained as candidate edges. Node positions were computed with the Fruchterman-Reingold algorithm (ggraph, layout = "fr"), using only the positive-ρ edges (weight = ρ) to drive attraction, so that types with stronger positive co-enrichment are pulled closer together on the page; negative-ρ edges were excluded from the layout calculation and added afterward for display only. All retained edges are drawn, with color scaling the direction of co-enrichment (blue, negative; red, positive) and width scaling with $|\rho|$.

**MERFISH spatial transcriptomics**

BM trephine biopsies were profiled on the MERSCOPE platform (Vizgen) using an 815-gene immuno-oncology panel. Twenty-two samples from 22 patients entered the pipeline. Segmented objects were retained as cells if they contained at least 6 transcripts from at least 5 distinct genes. Of 306,445 objects, 38,260 (12.5%) passed. Eighteen of 22 patients retained at least one cell, and 12 retained ≥50 cells (pre-specified minimum for spatial analysis), so all spatial analyses used these 12 patients. The low recovery rate reflects the archival, decalcified FFPE source material rather than the filtering thresholds.

Raw transcript counts were normalized to 10,000 transcripts per cell, log1p-transformed, and scaled per gene without centering (zero_center = False), keeping all values at or above zero. Scaling was applied for clustering only, annotation used log-normalized values. Principal component analysis (25 components) was followed by Harmony batch correction across all cells simultaneously, with the per-patient tissue export as batch variable (each MERSCOPE run covered several patients, exported separately). A k = 15 neighborhood graph, UMAP (min_dist = 0.5, spread = 1.0) and Leiden clustering (resolution = 1.0) produced 44 clusters. All random seeds were fixed at 0. Marker genes were called per cluster by Wilcoxon rank-sum test on log-normalized values, retaining the top 10.

Each cluster was matched against 23 curated reference gene sets covering myeloid, lymphoid, endothelial and stromal populations, and AML blast states. A cluster was only annotated when at least 2 genes, having real transcripts, of its top 10 genes appeared in the reference gene set. Hypergeometric P – value using the 815 gene panel as a background was the tie breaker in case a cluster had the same number of overlapping genes with two or more reference gene sets. Clusters failing the conditions, remained unannotated and were excluded from the downstream analysis. Out of 44 clusters, 26 clusters met the conditions and got a label (13 high, 6 moderate, and 7 low confidence) while 18 were unannotated. After obtaining the labels of the clusters, we utilized the CODEX panel to further annotate the MERSCOPE panel-based clustering, merging and gating specific populations including macrophages and stromal cells. The final annotation comprises 13 labelled CTs and one unannotated population.

Transcript detection was constrained throughout by the archival decalcified material, which limited resolution of lineage markers and of some populations. *ITGAM* (CD11b) transcripts, for instance, were detected in approximately 11% of cells assigned to the IM_CD11b population; this population is therefore designated CD11b-enriched immature myeloid cells rather than uniformly CD11b-positive.

**Expression-based ligand-receptor scoring**

We curated a list of 49 ligand- receptor pairs (45 ligands and 40 receptors) spanning chemokine, checkpoint, co-stimulatory, niche, myeloid growth, vascular, suppressive and adhesion families. Each pair was scored as the product of the ligand's mean expression across all cells of a sender type and the receptor's mean expression across all cells of a receiver type. A null distribution was generated by shuffling CT labels across all cells 10,000 times. P values for all sender × receiver × pair combinations were corrected together by Benjamini-Hochberg ($q < 0.05$). This analysis uses expression values only and incorporates no spatial information.

**Interface-restricted ligand-receptor scoring**

A neighborhood radius was estimated as the 90th percentile of the distance to each object's 16th nearest neighbor. Across 12 patients with at least 50 cells, the radius range was from 23.2 to 50.3 micrometers with a median of 31.2 micrometers. For each pair of cells in contact (within the sample specific neighborhood radius), we identified the sender cells that had at least one receiver cell in their neighborhood and the receiver cells that had at least one sender cell in their neighborhood. The interaction between the neighboring ligands and receptors was scored by multiplying the mean log normalized expression value of the ligand and receptor. From this interface product, we subtracted the same product from non-neighboring cells so that the scores represent enrichment at the interface rather than overall expression. At least 5 cells were required for each of the four groups. Since ligands and receptors are expressed in both sender and receiver cells, the scores consider both directions. For each ligand-receptor pair, the per patient values were tested against zero with Wilcoxon signed rank test and P-values were corrected for multiple testing across all pairs with Benjamini-Hochberg procedure. No ligand-receptor interaction reached statistical significance at the interface after correction.

**Interface contact between T-cells and CD11b-enriched immature myeloid cells**

A T-cell was defined as in interface contact with the IM_CD11b population when at least one IM_CD11b centroid lay within the sample-specific neighborhood radius. Genes were called positive at ≥1 detected transcript. Two comparisons were made. First, the proportion of T-cells expressing any of seven inhibitory receptors (*PDCD1*, *CTLA4*, *HAVCR2*, *LAG3*, *TIGIT*, *ENTPD1*, *CD244*) was compared between T-cells with at least one IM_CD11b neighbor and T-cells with none. Second, among T-cells in contact with IM_CD11b cells, the proportion expressing *PDCD1* was compared between those with at least one *VSIR*+ IM_CD11b neighbor and those whose IM_CD11b neighbors were all *VSIR*−. Cells were pooled across patients, so these comparisons are descriptive and not adjusted for between-patient variation. The analysis establishes physical adjacency only. Transcript counts do not report membrane protein abundance or receptor engagement, and no inference about VISTA-PD-1 ligation is made.

**Statistical analysis and software**

Statistical analyses were performed with Python and R[7]. Data are presented as means ± SD. Group means were compared using t-tests for single comparisons, while one-way ANOVA was employed for multiple comparisons. Cox proportional-hazards regression tested association between each spatial metric (permutation z-score, adjusted-enrichment residual, or co-enrichment ρ) and overall survival. For every candidate predictor, a univariate model was fit. Predictors with fewer than 15 patients with complete survival data were excluded. Raw p-values are reported for results of Cox regression models. Statistical significance was defined as $p<0.05$. Symbols indicate significance levels as follows: *p<0.05, **p<0.01, ***p<0.001, ****p<0.0001. Details on statistical analysis, including the number of biological and technical replicates for each experiment, are provided in the figure legends.
